# Use of a counterselectable transposon to create markerless knockouts from a 18,432-clone ordered *M. bovis* BCG mutant resource

**DOI:** 10.1101/817007

**Authors:** Katlyn Borgers, Kristof Vandewalle, Annelies Van Hecke, Gitte Michielsen, Evelyn Plets, Loes van Schie, Sandrine Vanmarcke, Laurent Schindfessel, Nele Festjens, Nico Callewaert

## Abstract

Mutant resources are essential to improve our understanding of the biology of slow-growing mycobacteria, which include the causative agents of tuberculosis in various species, including humans. The generation of deletion mutants in slow-growing mycobacteria in a gene-by-gene approach in order to make genome-wide ordered mutant resources is still a laborious and costly approach; despite the recent development of improved methods. On the other hand, transposon mutagenesis in combination with Cartesian Pooling-Coordinate Sequencing allows the creation of large archived *Mycobacterium* transposon insertion libraries. However, such mutants contain selection marker genes with a risk of polar gene effects, which is undesired both for research and for use of these mutants as live attenuated vaccines. In this paper, a derivative of the Himar1 transposon is described, which allows the generation of clean, markerless knockouts from archived transposon libraries. By incorporating *FRT* sites for FlpE/*FRT*-mediated recombination and *I-SceI* sites for ISceIM-based transposon removal, we enable two thoroughly experimentally validated possibilities to create unmarked mutants from such marked transposon mutants. The *FRT* approach is highly efficient but leaves an *FRT* scar in the genome, whereas the *I-SceI* mediated approach can create mutants without any heterologous DNA in the genome. The combined use of CP-CSeq and this optimized transposon was applied in the BCG Danish 1331 vaccine strain (WHO reference 07/270), creating the largest ordered, characterized resource of mutants in a member of the *M. tb* complex (18,432 clones, mutating 83% of the non-essential *M. tb* homologues), from which clean knockouts can be generated.

**Importance:** While speeding up research for many fields of biology (e.g. yeast, plant, and *C. elegans*), genome-wide ordered mutant collections are still elusive in mycobacterial research. We developed methods to generate such resources in a time- and cost-effective manner, and developed a newly engineered transposon from which unmarked mutants can be efficiently generated. Our library in the WHO reference vaccine strain of *M. bovis* BCG Danish targets 83% of all non-essential genes and was made publicly available via the BCCM/ITM Mycobacteria Collection. This resource will speed up *Mycobacterium* research (e.g. drug resistance research, vaccine development) and paves the way to similar genome-wide mutant collections in other strains of the *M. tb* complex. The stretch to a full collection of mutants in all non-essential genes is now much shorter, with just 17% remaining genes to be targeted using gene-by-gene approaches, for which highly effective methods have recently also been described.

## Introduction

Tuberculosis (TB) is one of the main threats to public health in large areas of the world (1). Drug resistance is on the rise and an effective vaccine is still lacking. For many types of *Mycobacterium* research, e.g. into potential drug targets, and for live attenuated vaccine engineering, mycobacterial mutants are required. However, the genetic manipulation of slow-growing mycobacteria has long been problematic due to their slow growth, the low efficiency of classical transformation methods and the high frequency of illegitimate recombination. The lab of W. R. Jacobs Jr. has, however, optimized specialized transduction in such a way that we now have a feasible method to create specific deletion mutants in slow growing mycobacteria (2, 3). More recently, Murphy and others developed ORBIT (Oligonucleotide-Mediated Recombineering followed by Bxb1 Integrase Targeting) for the creation of mutants with targeted deletions, insertions, or fusions in the mycobacterial chromosome (4).

Nevertheless, even with the improved engineering methods, gene by gene mutation of the entire mycobacterial genome still takes a long time and a lot of effort and expense, and no publicly available mutant collections that approach completeness are available, forcing most investigators to invest in the generation of their own mutants of interest, time and again. On the other hand, transposon mutagenesis allows the creation of a wide variety of mutants in an isogenic mycobacterial background in a quick and efficient manner. However, it yields a complex mixture of transposon mutants. Such mutant mixtures are useful for analyzing competitive mutant behavior in bulk (5), but are not well-suited for extracting single mutants. Ordered transposon insertion libraries can be picked from such random mutant mixtures and screened for insertions in a locus of interest (6, 7), but this is laborious and needs to be repeated for every gene.

In a previous study, we described an easy-to-implement, cost- and time-effective, massively parallel sequencing based approach (called Cartesian Pooling-Coordinate Sequencing or CP-CSeq) to characterize ordered transposon libraries (8). These libraries consist of multiwell plates in which each well contains a transposon mutant colony picked from agar plates of the bulk transposon mutant library. The method combines a pooling strategy of a stack of 96-well plates along the Cartesian coordinates (x, y, z) with multiplex transposon sequencing and reports both on the identity as well as on the location of the transposon mutants in the plate stack. We illustrated the approach for a large 9,216 (96×96) clone ordered transposon insertion mutant library of a streptomycin-resistant lab strain of *M. bovis* Bacillus Calmette-Guérin (BCG) Pasteur. At that time, this library was the largest characterized resource of *M. bovis* BCG mutants and had identified clones with disruptions in 64% of the non-essential *M. tb* gene orthologs (8). A range of investigators have since made use of this resource to catalyze their research, which motivated us to further innovate to substantially improve on this proof-of-concept resource.

Although transposon insertion mutants are valuable for fundamental mycobacterial research, they also have a few drawbacks. This is mainly due to the presence of transcriptionally active transposon-associated sequences, especially an antibiotic selection marker, increasing the risk for polar effects. We have recently observed such an effect in a vaccine research project in our lab, where transposon insertion in the *sapM* gene resulted in a strong enhancement of expression of the upstream *upp* gene, necessitating a significant effort in complementation experiments and laborious generation of a ‘clean’ *sapM* knockout (KO) to exclude a role for this Upp upregulation in the observed enhanced vaccine efficacy of *sapM* mutation in BCG (9). Presence of antibiotic selection markers also precludes the use of such transposon mutants for further (pre)clinical development as vaccines. Hence, we had a strong incentive to solve this problem and to investigate how selection marker-free and even heterologous DNA-free mutants could be derived from transposon mutants.

Here, we describe the construction of a derivative of the Himar1 transposon to allow subsequent generation of unmarked KOs from marked transposon mutants. Amongst several other methodological options that were tested but found impracticable (including CRISPR-Cas9 inspired sequence deletion), we found two effective possibilities to create unmarked mutants: use of *FRT* sites for FlpE/*FRT*-mediated out-recombination of transcriptionally active transposon parts and *I-SceI* sites for I-SceI meganuclease (ISceIM)-based transposon removal. The first approach inherently leaves an FRT scar, whereas the last one has the potential of creating clean, markerless mutants. To make this practicable, a focused effort was needed to explore different genetic carriers for the unmarking enzymes, both in fast-growing (*M. smegmatis*) and in slow-growing mycobacteria (*M. bovis* BCG). Finally, we combined the use of this ‘unmarkable’ transposon and CP-CSeq in the clinically used BCG Danish 1331 vaccine strain (WHO reference strain, NIBSC 07/270), creating a validated characterized ordered transposon insertion library of 2×96×96 well plates or ~18,000 clones. The library can be used in its entirety, or in ordered sub-collections, e.g. for chemical genomics screens or *in vitro* tests of interactions with immune cells. Transposon mutants of interest can be retrieved, and subsequently unmarked to create ‘clean’ mutants to validate initial findings. Ultimately, these ‘clean’ mutants can even be used directly for (pre)clinical vaccine development.

## Results

### Himar1 transposon re-designed to enable subsequent mutant unmarking

Several site-specific recombination systems have already been used in mycobacteria to allow the removal of an antibiotic resistance cassette, including the γδ-resolvase/*res* (2, 10–13), the FlpE/*FRT* (14–16), the Cre/*loxP* (16, 17) and the Xer/*dif* system (18, 19). The γδ resolvase/*res* and FlpE/*FRT* system were the first ones shown to be functional in mycobacteria (16). We chose to implement the FlpE/*FRT* system from *Saccharomyces cerevisae,* as for this system a higher recombination frequency was reported compared to the γδ-resolvase system (>40% versus 3-5% in slow-growing mycobacteria) (15).

Another strategy to remove the antibiotic resistance cassette is by using meganucleases. These engineered versions of naturally occurring restriction enzymes typically have extended DNA recognition sequences (12-40 bp). The I-SceI meganuclease (ISceIM) is such an example; it recognizes a specific 18 bp sequence that statistically occurs only once every 10^10^-10^11^ bp. Most organisms’ genomes lack an endogenous *I-SceI* site, including most, if not all *Mycobacterium* species. The *I-SceI* system has been used to unmark mutants in several organisms (20–24), but not yet for unmarking of mycobacterial mutants. We saw the opportunity to use it for this purpose, as in *M. smegmatis*, the *I-SceI* system has already been used to study the repair mechanisms of double-stranded DNA breaks (DSBs) (25, 26). The created DSBs either resulted in death of the cell, or were repaired by non-homologous end joining (NHEJ), which added additional nucleotides prior to re-ligation, or resulted in deletions (26).

We designed a derivative of the original Himar1 transposon (27), in which we incorporated both the *FRT* and the *I-SceI* system to enable creation of unmarked mutants from marked transposon mutants (**Fig. 1a–b**). The *FRT* sites are present as direct repeats, so that the DNA sequence in between will be deleted upon site-specific recombination. The *I-SceI* sites are in inverted orientation, so that the overhanging ends are not compatible. This will avoid that a simple ligation can resolve the created DSB, since we want DSBs to be repaired by NHEJ, with its associated exonuclease-mediated shortening of the ends. This exonuclease activity can be extensive, and we hypothesized that it may sometimes degrade all of the transposon sequences prior to re-ligation. In addition, we introduced the negative selection marker SacB to allow for direct selection of unmarked clones on sucrose. A transposon plasmid, containing this optimized transposon and the hyperactive Himar1 transposase gene, was constructed (**Method S1**). Subsequently, we produced an optimized transposon phage using the improved shuttle phAE159 from the lab of W. R. Jacobs Jr., which has increased cloning capacity and efficient DNA delivery (2). This was essential, as the optimized transposon (~4 kbp) is 2 kbp larger than the original Himar1 transposon (~2 kbp). Furthermore, this larger size of the transposon would be predicted to lead to a ~74% decrease in transposition frequency (28). While we did not formally tested this, indeed, transposon mutagenesis of the BCG Danish strain yielded ~25,000 mutants for the new, larger transposon, under conditions that would yield ~100,000 mutants described in the literature for the original Himar1 transposon (29, 30), and as we have indeed observed earlier in our own lab (8).

**Figure 1.**
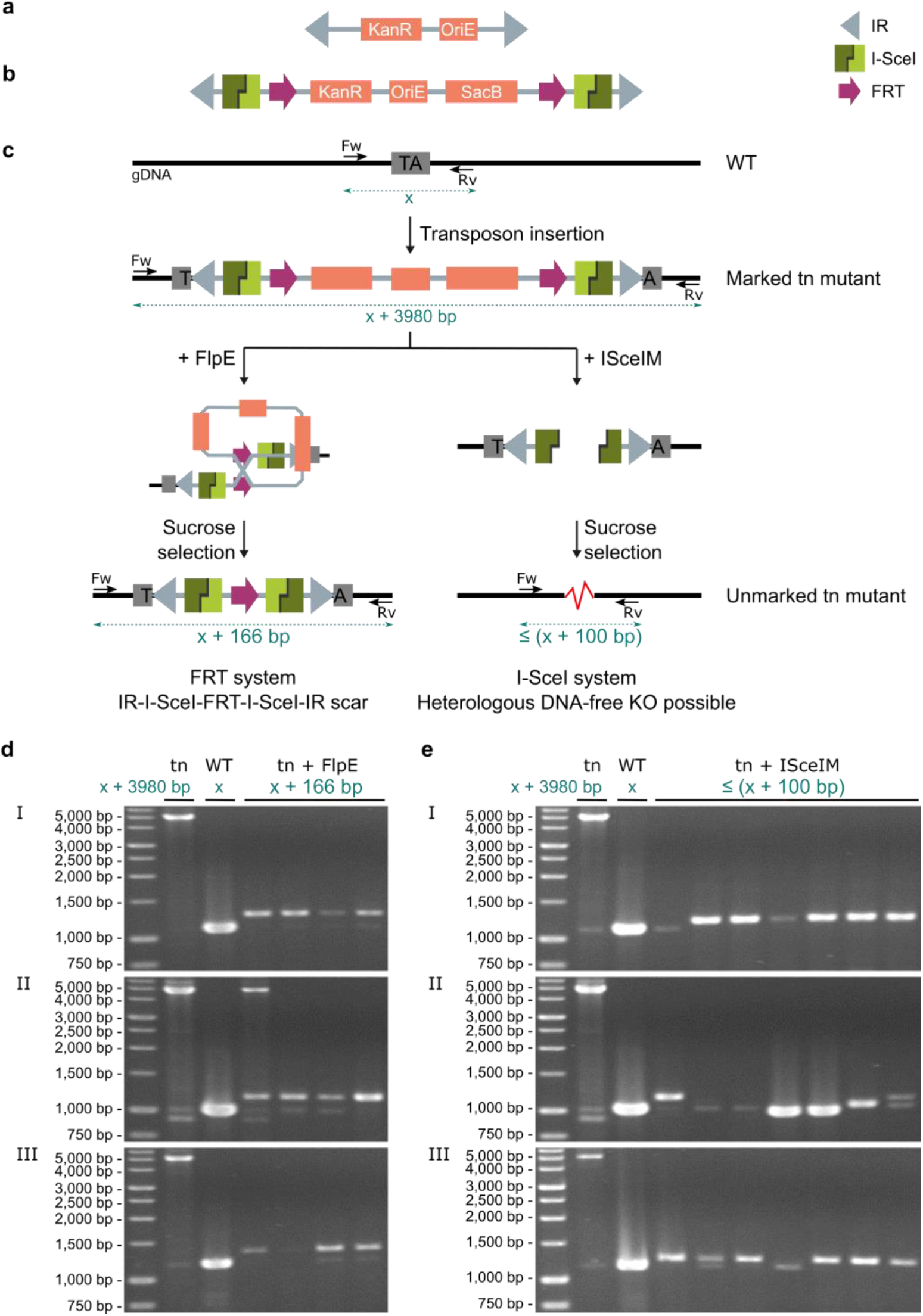
Creating clean KOs from transposon mutants using an optimized Himar1 transposon. **a)** Visual representation of the original Himar1 transposon (2,064 bp); consists of a kanamycin resistance marker (KanR) (795 bp) and an origin of replication for *E. coli* (OriE) (732 bp) flanked by two inverted repeats (IR) (27 bp). **b)** Optimized Himar1 transposon (3,980 bp); consists of a KanR (795 bp), an OriE (614 bp) and a SacB negative selection marker (1,419 bp) flanked by two *FRT* sites (34 bp), two *I-SceI* sites (18 bp) and two IR (27 bp). **c)** Illustration of the construction of a transposon mutant (marked) using the optimized transposon and subsequent unmarking strategies by bringing in FlpE or ISceIM into the cells and selecting the unmarked mutants. FlpE/*FRT*-mediated recombination inherently leaves an FRT scar (IR-*I-SceI*-*FRT*-*I-SceI*-IR, 166 bp) in the unmarked transposon mutant, whereas ISceIM-based transposon removal has the potential of creating KO mutants free of heterologous DNA. **d-e)** Proof-of-concept of the unmarking of three *M. bovis* BCG Danish transposon mutants (I, II and III). Transposon mutants were unmarked via FlpE/*FRT*-mediated recombination (**d**) or ISceIM-based transposon removal (**e)**. Therefore, replicable vectors expressing FlpE or ISceIM were introduced in the cells via electroporation, where after unmarked mutants (kanamycin sensitive and sucrose resistant) were analyzed via colony-PCR. Parental transposon mutants (I, II and II) and the original wild-type (WT) strain were taken along as controls. Length of expected amplicons are indicated in blue in subfigures c, d and e, x: length of WT amplicon, tn: transposon. Used primers are listed in **Table S1**.

Introduction of FlpE in the marked transposon mutants was designed to lead to site-specific recombination at the *FRT* sites, resulting in removal of most of the transposon-associated sequence, but leaving an IR-*I-SceI-FRT-I-SceI*-IR scar in the genome (**Fig. 1c**). Every FlpE-unmarked transposon mutant will contain the same scar in its genome. A proof-of-concept of this strategy was obtained by introducing FlpE in the marked transposon mutants using a standard replicable vector. Colony PCR performed on FlpE-unmarked transposon mutants indeed showed, for all clones, an amplicon of a size that was consistent with a 166 bp insertion as compared to the WT amplicon, both for fast-growing *M. smegmatis* (**Fig. S1**) as for slow-growing BCG Danish transposon mutants (**Fig. 1d**).

Introduction of ISceIM in the marked transposon mutants was designed to form a DSB that is repaired by NHEJ (**Fig. 1c**) (26). Hereby, more extensive exonuclease activity prior to DSB repair will lead to a different outcome. In a small percentage of cases, heterologous DNA-free KOs in which all of the transposon sequences were deleted, can be formed. A proof-of concept of this strategy using a standard replicable vector to introduce ISceIM in the marked transposon mutants, indeed showed that the amplicon length from ISceIM-unmarked clones differed, as shown for ISceIM-unmarked *M. smegmatis* (**Fig. S1**) and BCG Danish transposon mutants (**Fig. 1e**).

### Comparison of different genetic carriers for FlpE or ISceIM to unmark the transposon mutants

Standard replicable vectors proved to be sufficient to unmark transposon mutants using FlpE or ISceIM, but are known to be more difficult to cure from slow-growing bacteria. Therefore, we also produced several genetic carriers that can be easily cured from the cells or do not replicate in the cell. We compared three different vector systems (standard replicable vectors, replicable GalK vectors and temperature sensitive (ts) GalK vectors) and ts phages (**Fig. 2**). All genetic carriers were tested in both fast-growing (*M. smegmatis)* and slow-growing mycobacteria (BCG Danish) and evaluated for three characteristics: the transformation efficiency, the unmarking efficiency and the curing efficiency (if applicable). The results are summarized in **Fig. 2** and discussed below.

**Figure 2.**
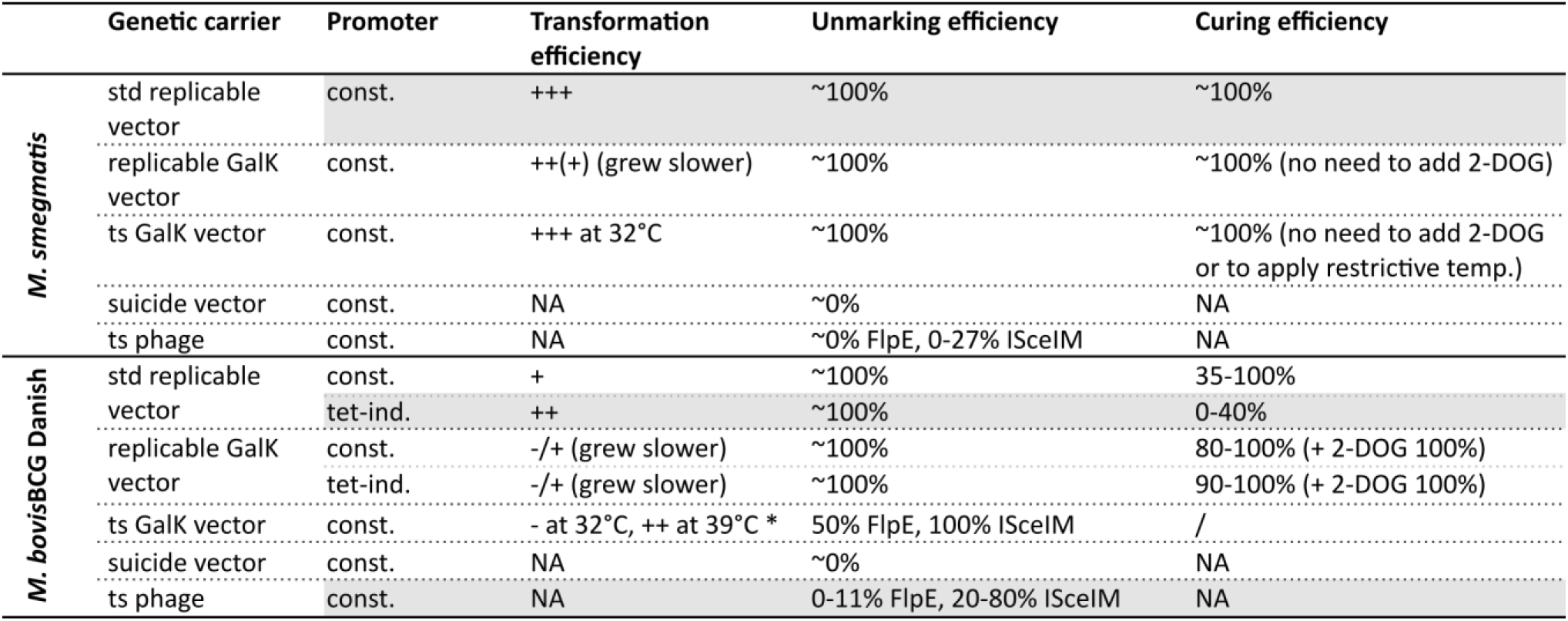
Unmarking *M. smegmatis* and *M. bovis* BCG Danish transposon mutants using different genetic carriers bringing FlpE or ISceIM to expression. 2-3 transposon mutants for *M. smegmatis* (upper panel) or *M. bovis* BCG Danish (lower panel) were unmarked by introducing FlpE or ISceIM by different genetic carriers, expressing FlpE or ISceIM from a constitutive or inducible promoter. The efficiency of transformation (amount of hygromycin-resistant clones) was given a score ranging from - to +++ based on the observed colony-forming units after incubation at 37°C (unless otherwise indicated, e.g. for the ts vectors). The efficiency of unmarking is indicated by the percentage of unmarked clones (kanamycin sensitive and sucrose resistant) among the sucrose-resistant clones (and hygromycin resistant for replicable vectors) obtained after unmarking. The efficiency of curing is indicated by the percentage of cured clones (hygromycin sensitive). All experiments were replicated once or more (e.g. replicable vectors), except the data for the replicable GalK vectors with the tet-inducible promoters, as no improvement was seen in the transformation efficiency compared to the replicable GalK vectors with the constitutive promoters in the first experiment. NA: not applicable (vector or phage can’t replicate in mycobacteria), /: not performed (would not give any added value), std: standard, GalK: negative selection marker galactokinase, ts: temperature sensitive, acet. ind.: acetamide inducible, tet-ind.: tetracyclin inducible. *: problem with the use of the ts vectors in BCG Danish: BCG Danish did not grow at 32°C, but grew at 39°C. In grey, we indicated the most preferred tools.

For *M. smegmatis*, we got similar results for a replicable vector with or without counter-selectable marker GalK and with or without a ts oriM: they all had an equal unmarking and curing efficiency.

The negative selection marker GalK, nor the ts oriM were needed to establish curing; subculturing the clones and plating them on non-selective plates was sufficient to cure the vectors at +/− 100% efficiency (**Fig. 2**). The standard replicable vector was the best option for unmarking *M. smegmatis* transposon mutants, as the GalK vectors showed a slightly lower transformation efficiency and a slower clone growth.

For BCG Danish, the ts GalK vectors were not feasible to unmark the transposon mutants, because transformed cells showed little to no growth at 32°C. Similar to the situation in *M. smegmatis,* the replicable GalK vector conferred more stress on the cells, resulting in a lower transformation efficiency and smaller colonies. On the other hand, due to apparent stress associated with the incorporation of the negative selection marker GalK, the replicable GalK vectors were more easily cured from the cells compared to the standard replicable vectors, with or without adding 2-DOG (2-deoxy-galactose) to the agar plates.

Suicide vectors and ts phages expressing FlpE or ISceIM do not need to be cured, as they cannot replicate (they lack an origin of replication for mycobacteria). This is advantageous, as unmarking can be done in a single round of transformation and selection, which is less time-consuming. However, they were not useful to unmark *M. smegmatis* transposon mutants, as their unmarking efficiency was very low. This was due to a high number of SacB escape mutants and the inability to select for uptake of the genetic constructs (the suicide vector or the ts phage). In BCG Danish, the same held true for the suicide vectors. However, the unmarking efficiency for the ts phage was satisfactory in BCG Danish, especially for ISceIM (20-80%), allowing their use to unmark transposon mutants in slow-growing mycobacteria. Nonetheless, the unmarking of our transposon mutants with these ts phages was still less efficient as compared to an earlier report on unmarking of specialized transduction mutants containing *res* sites with phAE280 (ts phage expressing γδ-resolvase), reportedly reaching up to 80-100% efficiency (Jain *et al.* 2014 (2), and our own unpublished observations).

As initial experiments (not shown) demonstrated that FlpE and ISceIM expression can be toxic in slow-growing mycobacteria, strongly reducing transformation efficiency, we also constructed tetracyclin-inducible vectors, next to the constitutive vectors. For the tetracyclin-inducible vector, we indeed observed a higher transformation efficiency compared to the respective constitutive vector, reliably obtaining ~100-200 clones for one transformation with the tet-inducible promoter, while in some experiments no unmarked clones were obtained with the constitutive promoters. Most likely because they cause less stress, the tetracyclin-inducible vectors were less efficiently cured. We tested the addition of the GalK counter-selectable marker to enhance the curing efficiency of these vectors. Unfortunately, when introducing GalK, we lost the positive effect of the tetracyclin-inducible promoters on the transformation efficiency. Probably, the toxic stress effect of GalK, which we also observed in *M. smegmatis*, overpowered the reduced stress imparted by low expression of FlpE/ISceIM allowed by tetracyclin-inducible promotors. Nevertheless, the curing efficiency of the few unmarked clones that were obtained was indeed enhanced under counter-selective conditions.

We conclude that for *M. smegmatis*, the replicable vectors are the best option to unmark transposon mutants. For *M. bovis* BCG Danish, it is difficult to compare the efficacy of the plasmid methods with the unmarking phages, due to the inherent differences between the methods. Some practical advantages of the ts phages are however quite clear: (i) transduction was easier to perform than electroporation and was more efficient, (ii) the carrier did not need to be cured afterwards. However, not all recovered colonies were unmarked, unlike with the replicable vectors, where ~100% of hygromycin- and sucrose-resistant clones were unmarked. Nonetheless, with ts phages we could easily obtain ~200 clones after one transduction, after which a straightforward screening method (streaking on plates with and without hygromycin) allowed the identification of unmarked mutants. Overall, we give preference to the methods that yield a robust number of clones upon transgenesis of the unmarking tool, together with high unmarking efficiency and easy curing of the unmarking vehicle. The tet-inducible replicable vectors and the ts phages struck the best compromise for BCG.

### Different outcomes of the *I-SceI* system

In contrast to the *FRT* system (that always yields the same scar), the *I-SceI* system creates a DSB that is repaired by NHEJ, yielding a range of scar sizes. We classified all obtained ISceIM-unmarked transposon mutants in three categories (**Fig. 3a**): (i) mutants with two remaining IR (leaving a transposable transposon in the genome), (ii) mutants with one remaining IR or (iii) mutants with no transposon sequences remaining in the genome. Several ISceIM-unmarked transposon mutants were analyzed in depth, to investigate the different outcomes of NHEJ in fast-growing *M. smegmatis* (test model) (**Fig. 3b**) and slow-growing BCG Danish (target organism) (**Fig. 3c**).

**Figure 3.**
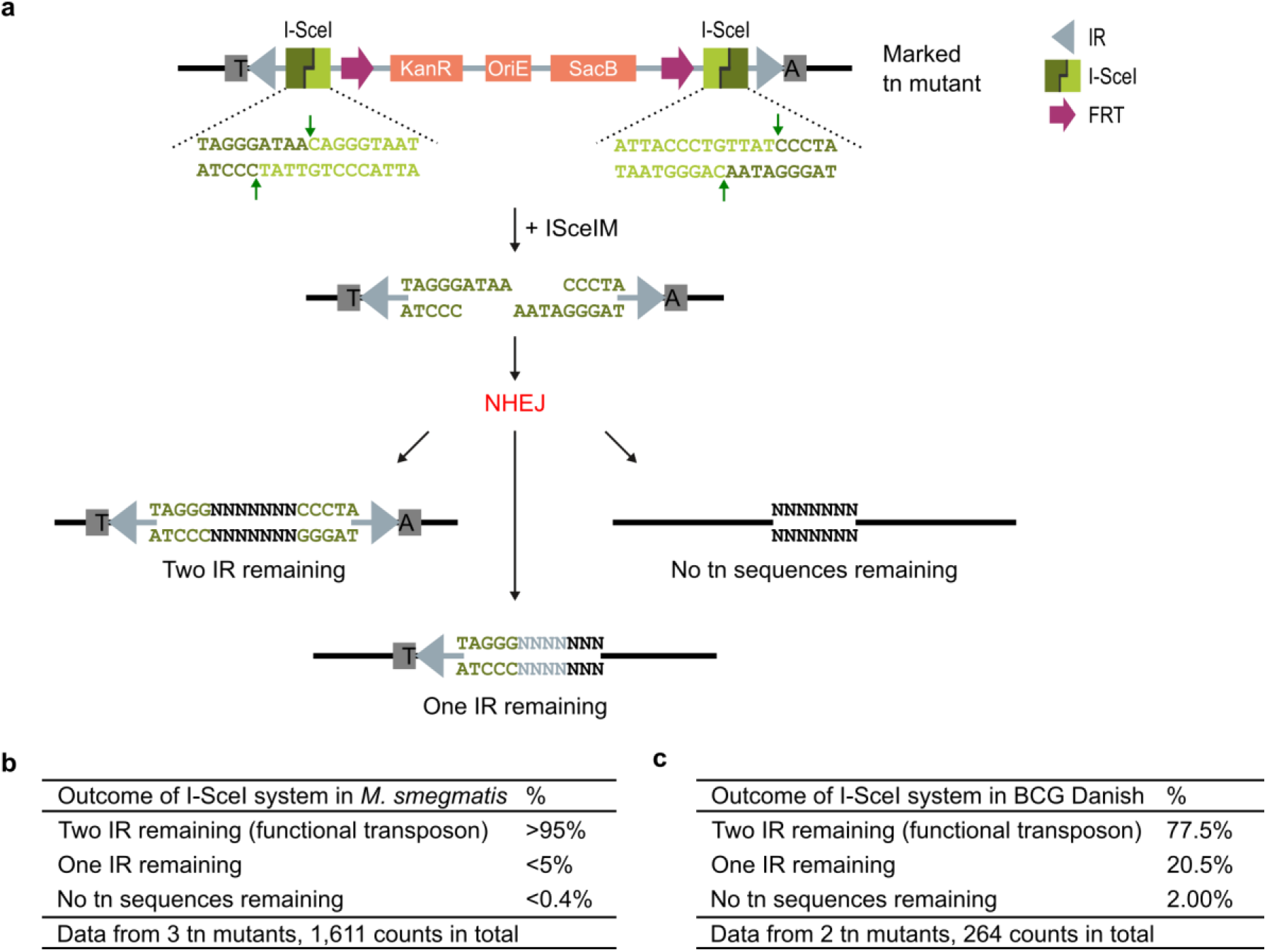
Outcome of unmarking transposon (tn) mutants via the I-SceI system. **a)** Graphic illustration of the outcome of unmarking via the *I-SceI* system. Upon introduction of ISceIM, the restriction enzyme will cut the *I-SceI* sites, creating a DSB that will be repaired by NHEJ, resulting in one of three possible outcomes. **b-c)** The outcome of the *I-SceI* system was analyzed for three *M. smegmatis* transposon mutants (**b**) and two BCG Danish transposon mutants (**c**) by performing PCR on kanamycin-sensitive and sucrose-resistant clones and Sanger sequencing the PCR amplicons (for *M. smegmatis* only a subset of the clones was Sanger sequenced). Outcome is given in percentages. Used primers are listed in **Table S1.**

For *M. smegmatis*, unmarking via the *I-SceI* system resulted mostly in small indels (>95%) in which two intact IR remain. Larger deletions occurred only for a small percentage, after which only one IR (<5%) or no transposon sequences remained (<0.4%). In depth analysis of the small indels was hard, as the repetitive nature of IR-indel-IR site (forms an almost perfect hairpin) completely obstructed Sanger sequencing of these indels.

Compared to *M. smegmatis*, the transformation efficiency in BCG Danish was lower, leading to fewer ISceIM-unmarked clones, but with a higher frequency of larger deletions after NHEJ than for *M. smegmatis*. In 21% of the unmarked clones only one intact IR remained in the genome, compared to <5% for *M. smegmatis*. In 2% of the cases, no transposon sequences remained in the genome, compared to <0.4% for *M. smegmatis*. Hence, the *I-SceI* system is a feasible way of generating heterologous DNA-free KOs from BCG Danish transposon mutants. If only inactivation of a transposable transposon is needed, destruction of one of the IRs is sufficient, which occurs at a ten-fold higher frequency (21% versus 2%).

### Development of a *Mycobacterium bovis* BCG Danish mutant resource

We have combined the use of this new transposon phage and CP-CSeq (8) in the BCG Danish 1331 vaccine strain (WHO reference 07/270), creating an ordered transposon insertion library of 2 sets of 96×96-well plates (**Data set S1**). Each transposon mutant in this library can be unmarked using the methods described above. CP-CSeq characterization of this library provided evidence for transposon insertions at 20,618 sites; 19,481 in unique genome regions and 1,137 in duplicated regions or highly repetitive genes (**Fig. 4a**). Theoretically, the library could only contain 18,432 mutants (i.e. the total number of wells in the library plates); the 2,186 mutants we have in plus can be explained in two ways. First, due to the clumping of BCG cells, many colonies on agar plates of the transposon library were likely not clonal. While we did the utmost in the clone picking to avoid mixed colonies, it was impossible to avoid this completely. This is also the reason why we stuck to manual picking by experienced scientists, rather than with robots, as these cannot yet achieve the same level of accuracy in distinguishing single colonies. Hence, 14% of the wells for set I and 11% for set II contained more than one clone. Second, due to a minimal aspecific amplification of the gDNA (higher background) during amplification of the transposon-flanking regions, a low level of spurious transposon insertion detection occurs. To limit this noise, we excluded transposon insertions that were detected with a total Cartesian sequencing read coverage below 60 (the average read depth was 4,709 ± 3,626 for the remaining non-noise transposon insertions).

**Figure 4.**
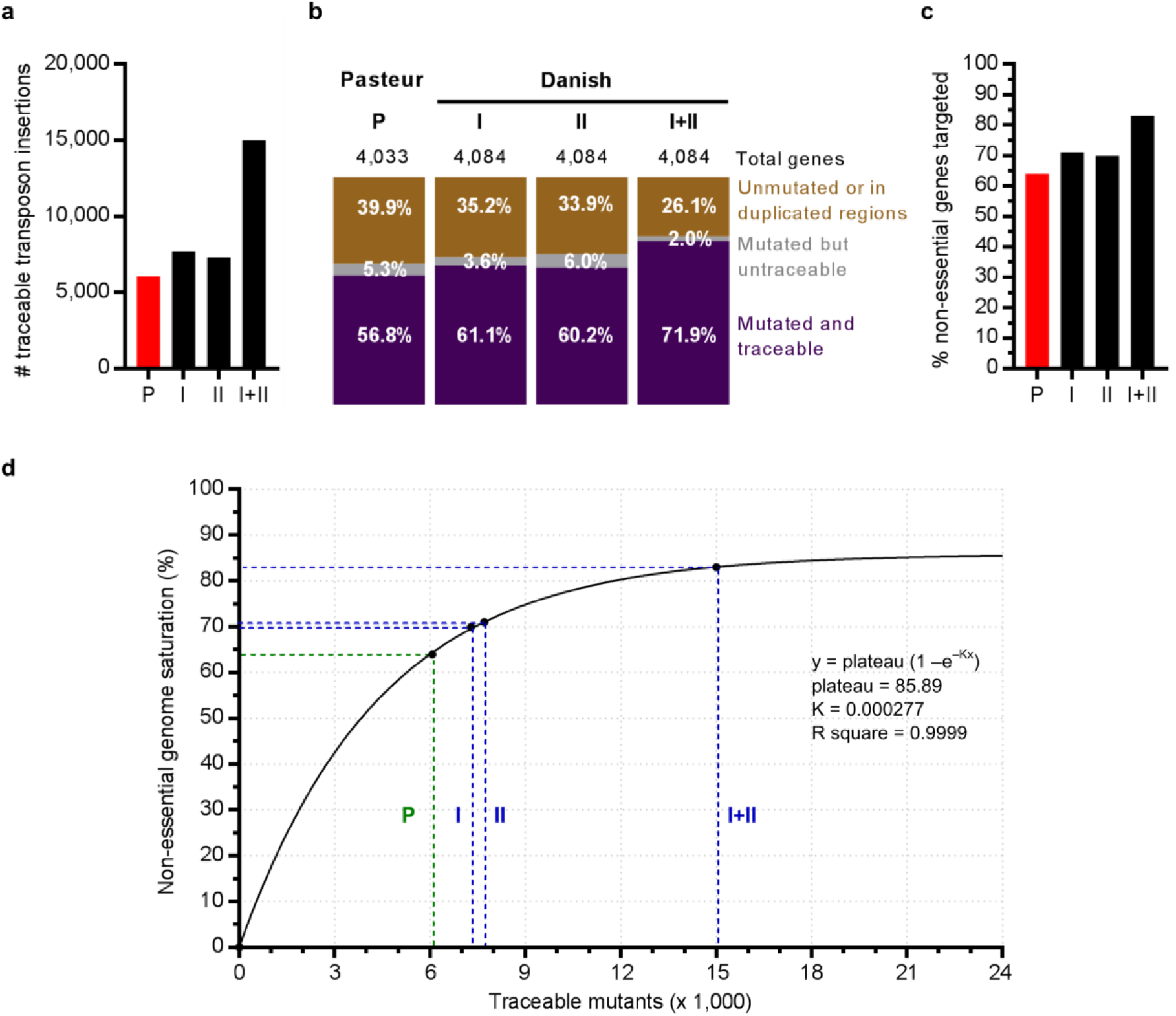
Results of CP-CSeq on a transposon insertion library of *M. bovis* BCG Danish (2 sets of 96×96-well plates, set I and set II, this study) compared to *M. bovis* BCG Pasteur (Vandewalle *et al.* 2015 (8)). **a)** Number of traceable transposon insertions for each library (set). **b)** Distribution between disrupted genes that are traceable (purple), disrupted genes for which the locations of their disruption mutants are unknown (‘untraceable’; grey) and untargeted genes or genes in duplicated regions (brown). **c)** Percentage of targeted non-essential genes for each library (set). **d)** Non-essential genome saturation in function of the amount of traceable mutants. In green, the observed values for the library in *M. bovis* BCG Pasteur (P). In blue, the observed values for the library in *M. bovis* BCG Danish (set I, set II and set I+II). To model what would be gained by further enlargement of CP-CSeq libraries, we fitted a Coupon’s collector’s equation (in black) to the available experimental data from our libraries, which should allow more accurate predictions in the future. The Coupon collector’s problem estimates the amount of traceable mutants needed to target a certain percentage of the non-essential genome. The equation for the curve is given in the graph. This curve goes not through 100%, probably due to inherent characteristics of transposon mutagenesis and clone-picking of transposon mutants grown on agar.

Using these settings for CP-CSeq, for 15,320 insertion events in unique regions of the BCG Danish genome, the well position of the mutant in the plate library could be identified; 7860 for set I and 7460 for set II (**Fig. 4a**). In the combined resource (set I + II), about 85% of these insertions mapped against genes, targeting 83% of all non-essential *M. tb* orthologs at least once (**Fig. 4b**). The current library in BCG Danish outperformed the previous library in BCG Pasteur (8), as the separate sets of the new library targeted 71% and 70% of all non-essential genes, while this was only 64% for the old library (**Fig. 4c**). This improvement was most likely due to experimental improvements in the construction of the libraries (**Method S7**).

As Himar1 transposon insertion is a random event that can occur at a finite number of TA sites in the genome, the relation between the ‘number of attempts’ and the chance of hitting increasing numbers of TA sites is akin to the ‘Coupon Collector’s problem’ in statistics. To model what would be gained by further enlargement of CP-CSeq libraries, we fitted a Coupon’s collector’s equation to the available experimental data from our libraries (**Fig. 4d**). As our combined library is very near to the plateau of this empirical model, it is clear that further enlargement of the library would not be effective. Indeed, as expected, the non-essential genes that remained untargeted (371 in nonduplicated regions) are short genes and/or contain few TA’s (**Table S2**), thus having much less chance to be mutagenized during transposon mutagenesis.

To validate the CP-CSeq assignments of transposon insertions to particular wells in the plate library, the position of 90 transposon mutants from 4 different plates was verified via PCR and all were correct (**Fig. 5**). Even though CP-CSeq had identified more than 96 transposon mutants in plate I-65, all 32 selected mutants of which we checked the position for this plate, were confirmed via PCR, indicating that the robustness of our approach is high in the face of unavoidable picking of clonally impure colonies. Additionally, we confirmed that we could easily recover mutants in a clonally pure form in all cases by streaking the well contents for single colonies on agar and picking a few colonies for PCR-testing (**Fig. S2**). This does not add much supplementary burden in practice, as in any case, it is good microbiological practice to streak for single colonies from culture collection resources prior to commencing further experimentation.

**Figure 5.**
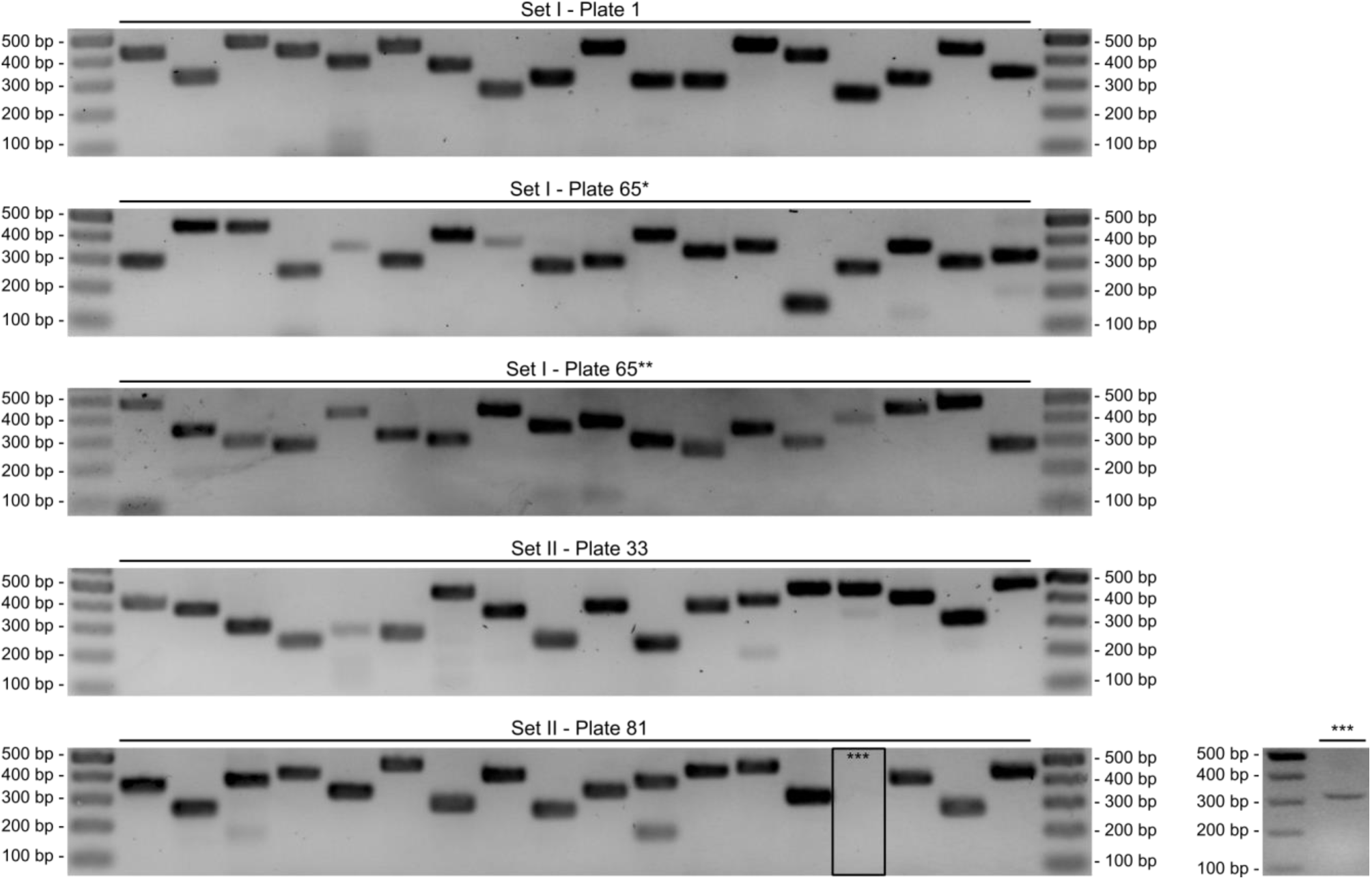
Positional validation of a subset of transposon insertion mutants by PCR. 90 CP-CSeq assignments for transposon insertion mutants coming from 4 different 96-well plates were verified by PCR with a primer hybridizing to the transposon IR and a gene specific primer. For plate I-1, II-33 and II-81, containing < 96 mutants, we selected 18 transposon insertion mutants with a coverage > 300. For plate I-65, containing >96 mutants, we selected 18 transposon insertion mutants with a coverage >1200 (I-65*) and 18 transposon insertion mutants with a coverage > 300 and/or in wells with > 1 mutant (I-65**). All CP-CSeq assignments were confirmed, except for 1 from plate II-81 (***). This PCR was successfully redone on a fraction of the glycerol stock (small gel panel on the right), so that all CP-CSeq assignments were confirmed. The list of mutants and their corresponding primers can be found in **Table S3**.

## Discussion

Transposon mutagenesis has been used throughout *Mycobacterium* research to establish genotype-phenotype relationships and to investigate the genetic requirements of *in vivo* persistence of pathogenic bacteria such as *M. tb*. It is by far the most effective way of generating genome saturating mutant libraries. However, there are two disadvantages: (i) specific mutants cannot be easily retrieved from the mixed libraries and (ii) the transposon-carried selection genes can trigger polar effects on neighboring genes. We solved the first problem previously by introducing CP-CSeq to characterize large clone-picked ordered 96-well plate collections obtained from transposon mutant libraries (8). Here, by construction of an optimized Himar1 transposon and optimized efficient unmarking tools, we enable the creation of clean, markerless mutants from marked transposon mutants to resolve the second problem. Combining this new transposon with the CP-CSeq protocol (creating archived mutant libraries) (8) now permits the creation of genomewide mutant resources, from which clean KOs can subsequently be efficiently generated (**Fig. 6**).

**Figure 6.**
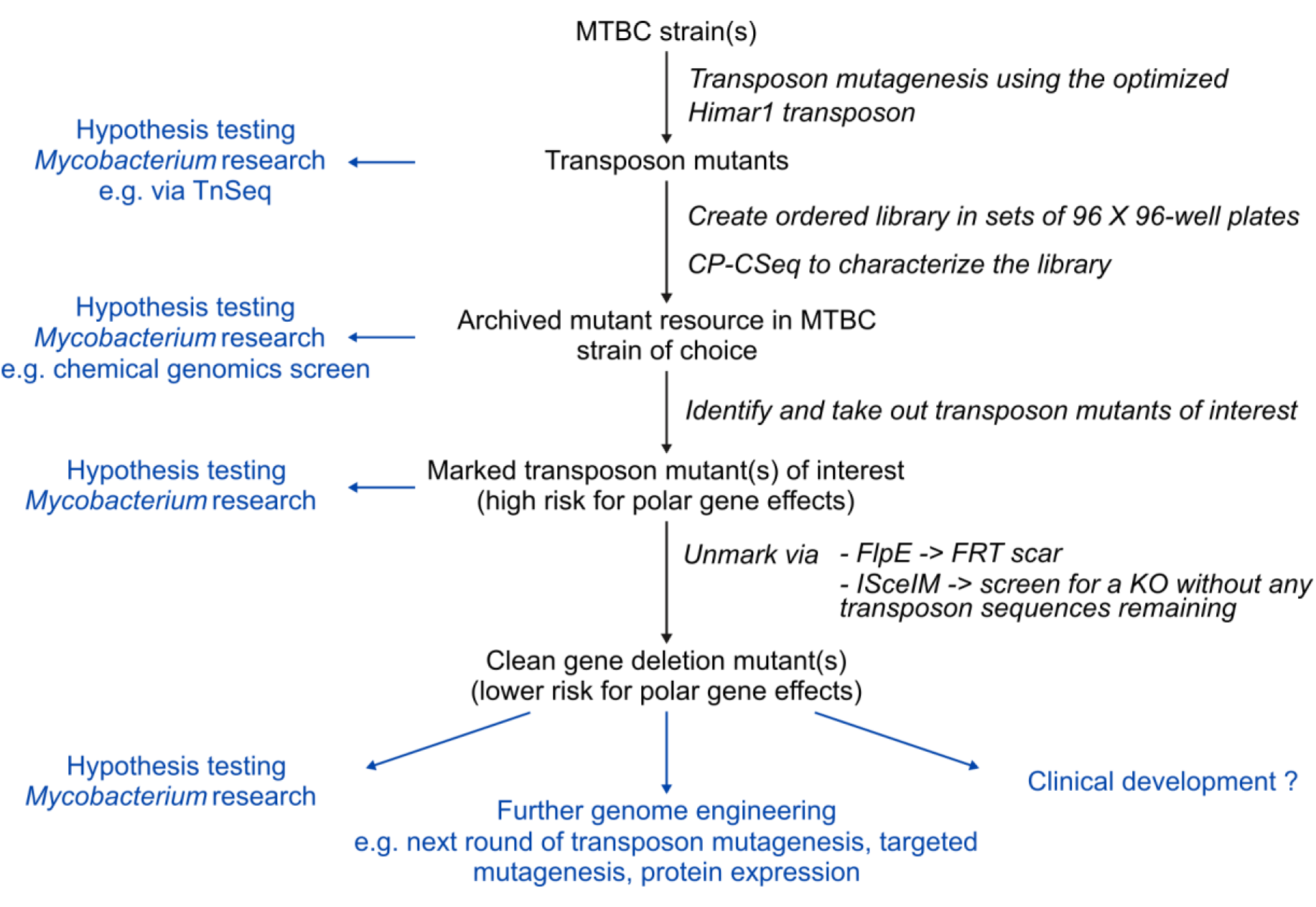
Applications of transposon mutagenesis using the optimized transposon. Overview of how an archived transposon library can be generated and how transposon mutants of interest can be isolated and unmarked. The scheme includes the different applications (in blue) of the generated products. MTBC: *Mycobacterium tuberculosis* complex.

By incorporating *FRT* sites for FlpE/*FRT*-mediated recombination and *I-SceI* sites for ISceIM-based transposon removal, we enable two possibilities to create unmarked mutants from marked transposon mutants. The first approach is highly efficient and inherently leaves an *FRT* scar, whereas the *I-SceI* system has the potential of creating clean, markerless mutants but requires clone screening. Once the transposon insertion mutants are created, generating the KO counterparts no longer requires complicated cloning or transduction procedures, but only a straightforward introduction of FlpE or ISceIM (via electroporation or transduction). As such, this research contributes to the global goal of creating genome-wide KO mutants in multiple strains of the *M. tb* complex.

The *I-SceI* system, described here, provides a new way to remove the antibiotic resistance cassette in mycobacteria. Although the *I-SceI* system requires screening to identify the clones of interest (in contrast to the site-specific recombination systems, like the *FRT* system), it is the only available strategy that leaves no target recognition sites in the genome. As the *I-SceI* sites are cut and cannot be reformed, the unmarked organisms have no intact *I-SceI* sites remaining in the genome, excluding the risk for unwanted genomic rearrangements during iterative construction of multiple mutant strains (31). The *I-SceI* system can now also be implemented in other mutagenesis methods (two-step allelic exchange, specialized transduction (2, 11) or ORBIT (4), for which unmarking of mutants generated with these methods still leave scars in the genome (32)), to create unmarked mutants.

We constructed an archived library of 2 sets of 96×96-well plates of optimized transposon mutants in the WHO reference strain *M. bovis* BCG Danish 1331 (NIBSC 07/270). Each of these transposon mutants can be unmarked to create markerless KOs for genes of interest. The created resource in BCG Danish (15,320 traceable transposon mutants) approaches genome saturation, as it targets 83% of all non-essential genes (excluding the duplicated genes). Applying (enhanced) specialized transduction (2, 3) or ORBIT (4) to target the remaining non-essential genes would be the most convenient option to complete the mutant resource, as the remaining genes are hard to target via transposon mutagenesis. For this biosafe BCG model, which is also the current TB vaccine and chassis for several of the improved BCG candidates, this goal now comes tantalizingly close. Only 371 non-essential genes remain to be targeted with targeted mutagenesis methods.

As a service to the TB research community, several research groups have done (or are doing) the effort to create large mutant resources and make them publicly available in order to catalyze mycobacterial research all over the world. Indeed, transposon insertions resources in *M. tb* (33, 34) and other mutants were integrated by the TARGET program (Tuberculosis Animal Research and Gene Evaluation Taskforce, http://webhost.nts.jhu.edu/target/), now offering over 4,300 defined transposon insertion mutants in *M. tb* (35). Furthermore, the lab of W. R. Jacobs Jr. (in collaboration with others) took on the effort to construct a comprehensive, ordered library of mutants, each mutated for a non-essential gene, based on specialized transduction (2, 3, 36). Supporting these efforts, we have deposited both of our BCG mutant resources (BCG Pasteur library of 6,072 traceable transposon mutants and this study’s BCG Danish library of 15,320 traceable, unmarkable transposon mutants) at the public BCCM-ITM Mycobacteria Collection (http://bccm.belspo.be/about-us/bccm-itm), making these mutants easy accessible to the scientific community. These resources are currently the largest resources of mutants in any strain of the *M. tb* complex that are publicly available. Replicating the effort for other *M. tb* complex strains is feasible and we hope that it will be pursued.

## MATERIALS AND METHODS

### Bacterial strains, media, plasmids and phages

The *M. bovis* BCG Danish 1331 sub-strain (first WHO Reference Reagent, 07/270, NIBSC, Hertfordshire) and the streptomycin-resistant *M. smegmatis* strain (a gift of Dr. P. Sander, Institute for Medical Microbiology, Zürich) were grown in Middlebrook 7H9 broth (Difco) supplemented with 10% Middlebrook OADC (Becton Dickinson), 0.2% glycerol and 0.05% Tween 80 (Sigma). For electroporation of *M. bovis* BCG and *M. smegmatis*, cultures were grown in 7H9 broth supplemented with 0.05% Tween 80 and 10% home-made ADS (5 g/L of BSA, 2 g/L of dextrose and 0.85 g/L of NaCl), no glycerol. Difco Middlebrook 7H10 agar supplemented with 10% OADC and 0.5% glycerol was used for growth of *M. bovis* BCG on solid culture, LB agar (10 g/l of BD Bacto Tryptone, 5 g/l of BD Bacto Yeast, 5 g/l of NaCl and 20 g/l of BD Bacto Agar) was used for growth of *M. smegmatis* on solid culture. A list of all plasmids, shuttle phasmids and phages used can be found in **Table S4**, detailed procedures for the construction of the plasmids, phasmids and phages generated in this study can be found in **Method S1**. Liquid or agar media were supplemented with 50 μg/ml of ampicillin, 50 μg/ml of kanamycin, 75 μg/ml of hygromycin, 2% or 5% sucrose, 0.2% or 0.5% 2-DOG (2-deoxy-galactose), 0.2% acetamide or 100 ng/ml of tetracyclin.

### Transposon mutagenesis

Bacteria were grown to an OD600 of 0.8-1.0 in supplemented 7H9 broth. Cells were washed 3 times with MP buffer and pre-warmed to 39°C. Next, warm phage solution (39°C), or MP buffer as negative control, was added to the warm bacilli. The transduction mixture was incubated for 4 hours at 37°C. After incubation, the transduction mix was plated on pre-warmed (37°C) selective plates containing 50 μg/ml kanamycin for incubation at 37°C. Additional details can be found in **Method S2**.

### Unmarking of transposon mutants via electroporation of unmarking plasmids

Bacteria were grown to mid-log phase (OD_600_ of 0.7-0.8) in 7H9 medium supplemented with Tween 80 and ADS. For *M. bovis* BCG cultures, 1.5% glycine was added to the cultures the day before electroporation. On the day of electroporation, cells were centrifuged and washed 3 times with 0.05% Tween 80 (in water). The cells were resuspended in the same buffer, after which the cells were mixed with the plasmid/phasmid (1 μg) in a pre-warmed 2 mm electroporation cuvette and electroporated (2.5 kV, 25 μF and 800 Ω). After recovery of the cells without antibiotics, the transformation mixture was plated on selective LB agar plates and incubated at 37°C (or at 32°C or 39°C when working with ts plasmids). The number of hygromycin-resistant clones was calculated as a measure for the efficiency of transformation. Details can be found in **Method S3**.

### Unmarking of transposon mutants via transduction of unmarking phages

Bacteria were grown to an OD_600_ of 0.8-1.0 in supplemented 7H9 broth. Cells were centrifuged, washed 3 times with MP buffer and resuspended in the same buffer. Next, 500 μl of ~10^10^ plaque-forming units of warm phage suspension, or MP buffer as negative control, was added to the bacilli. After incubation, the transduction mix was plated on selective plates for incubation at 37°C. Details can be found in **Method S4**.

### Analysis of unmarked transposon mutants

The colonies obtained after unmarking were analyzed for the presence or absence of resistance markers by streaking the clones on agar plates containing different supplements. The percentage of unmarked clones (kanamycin sensitive and sucrose resistant) among the obtained sucrose-resistant colonies was calculated as a measure of the efficiency of unmarking. Unmarked transposon mutants were subsequently analyzed via colony PCR and/or Sanger sequencing. The used primers are listed in **Table S1**, the primer sequences in **Table S5**, additional details in **Method S5**.

### Curing mycobacteria of unmarking plasmids

Unmarked mycobacteria carrying an unmarking plasmid, containing a hygromycin positive selection marker and/or a GalK negative selection marker, were inoculated in 7H9 broth and subcultured before making serial dilutions and plating them on agar plates with or without 2-DOG. After incubation, single colonies (at least 20) were streaked on agar plates with or without hygromycin to check if the unmarking plasmid was cured. The percentage of cured clones (hygromycin sensitive) was calculated as a measure for the efficiency of curing. Details can be found in **Method S6**.

### Construction and validation of the archived mutant library

The construction of the archived mutant library of two sets of 96×96-well plates in the *M. bovis* BCG Danish 1331 strain is based on the protocol by Vandewalle and others (8). For creating the transposon library, the medium was extra supplemented with 1% MD-VS vitamin supplement (ATCC), 0.2% Casamino Acids (Bacto) and 50 μg/ml of L-Trp, to allow the inclusion of auxotrophic mutants in the library. The 192×96-well plates containing the transposon mutants were processed in two sets of 96-96-well plates. We recently established a reference genome sequence for *M. bovis* BCG Danish 1331 (NIBSC 07/270, https://doi.org/10.6084/m9.figshare.c.4489496 (37, 38)), which was used to map the sequencing reads. A detailed procedure for the construction of the archived mutant library can be found in **Method S7**. The library size calculations are based on calculations described by Vandewalle and colleagues (8).

To verify the CP-CSeq assignment of the transposon insertion sites, frozen aliquots (from set I plate 1 and 65, set II plate 33 and 81) were heated to 98°C for 45 min to release gDNA in the medium. After centrifugation, the supernatant was used in a PCR reaction with a primer hybridizing to the transposon IR (Rv primer) and gene-specific primers (Fw primer) (**Table S3**). To analyze the clonal purity for a subset of transposon mutants, mutants were streaked on 7H10 agar plates supplemented with kanamycin, after which single clones were picked and grown in liquid medium. A PCR on supernatant of the BCG cultures and/or on gDNA of these single clones was performed with a unique primer set for the mutant (**Table S6**). PCR was performed using Phusion High Fidelity polymerase (NEB) after which PCR products were run on a 1.2% or 2% agarose gel, stained with ethidium bromide and visualized under ultraviolet light.

## ACKNOWLEDGEMENTS

Research was funded through a PhD fellowship to K.B. from the Flanders Innovation & Entrepreneurship agency (VLAIO), an ERC Consolidator grant ‘GlycoTarget’ to N.C. and VIB and UGhent institutional funding to N.C. The funders had no role in study design, data collection and interpretation, or the decision to submit the work for publication. We declare no conflicts of interest.

We thank the VIB Nucleomics core (http://www.nucleomics.be/, Leuven) for the Illumina NextSeq sequencing services, the VIB Genetics Service Facility (https://www.neuromicssupportfacility.be/, Antwerp) for the Sanger sequencing services and the Applied Bioinformatics and Biostatistics group (https://www.psb.ugent.be/applied-bioinformatics-biostatistics, Ghent) for the use of the Galaxy web platform (https://usegalaxy.org/) (39) via the Galaxy server of PSB. We thank Peter Sander for the streptomycin-resistant *M. smegmatis* strain. We thank William R. Jacobs Jr. for the improved shuttle phAE159 and for useful discussions and motivation. We thank Charlot Depoorter (Master 2 student Ghent University) for her help in the construction of some plasmids, phasmids and specialized transducing phages and for her assistance in optimizing and performing the library preparation.

K.B. designed and performed the experiments, analyzed the data and co-wrote the manuscript. K.V. assisted in experimental design and interpretation, and constructed some vectors. G.M. constructed some plasmids and performed some unmarking experiments and curing experiments. A.V.H. aided in the transduction, growing and pooling of the transposon insertion mutant library. E.P. aided in the plating after transduction. A.V.H., E.P., L.v.S., N.F. and S.V. assisted in picking the colonies. The Illumina library preparation and sequencing was performed by the VIB Nucleomics Core Facility. L.S. refined the BioPerl algorithm for automated location assignment. N.C. and N.F. assisted in experimental design and interpretation, and co-wrote the manuscript. N.C. conceived and supervised the project.

## DATA/CODE AVAILABILITY STATEMENT

Supplementary Material accompanies this paper at the Figshare repository with DOI https://doi.org/10.6084/m9.figshare.c.4507472, also including the Galaxy workflows for the bioinformatics data analysis and the updated BioPerl Script for the automated mutant location assignment. The BCG Danish 1331 genome assembly as used during the data analysis can be found on Figshare with DOI https://doi.org/10.6084/m9.figshare.c.4489496 (37). The novel sequencing data (raw Illumina reads) generated in this study have been deposited at NCBI’s Sequence Read Archive under project code PRJNA528037. The plasmids and phasmids generated in this study, have been deposited at BCCM/GeneCorner Plasmid Collection (http://bccm.belspo.be/about-us/bccm-genecorner), their LMBP accession numbers are listed in **Table S5**. The remaining data that support the findings of this study are available on request from the corresponding author N.C.

