## Supplementary figures, tables and methods for "Use of a counterselectable transposon to create markerless knockouts from a 18,432-clone ordered *M. bovis* BCG mutant resource"

### Supplementary Material

**Located at Figshare repository with DOI https://doi.org/10.6084/m9.figshare.c.4507472**

**In this pdf document:**

Figure S1. Validation of the unmarking of three *M. smegmatis* transposon mutants (I, II and III) via colony-PCR.
Figure S2. Recovering clonally pure cultures for a subset of transposon insertion mutants.
Figure S3. Practical implementation of Cartesian pooling using a 96-channel pipette.

Table S1. List of PCR and Sanger sequencing primers to analyze the unmarking of 3 transposon mutants for *M. smegmatis* and *M. bovis* BCG Danish.
Table S2. Non-mutated or untraceable non-essential genes are shorter and contain few TA's than mutated and traceable non-essential genes.
Table S3. List of primers for positional validation (library subset).
Table S4. List of plasmids, shuttle phasmids and phages used in this study.
Table S5. List of additional primer sequences
Table S6. List of primers to check clonal purity.
Table S7. List of gBlock sequences (and one PCR amplicon g00x).
Table S8. List of custom adaptors
Table S9. List of primers for library preparation

Method S1. Detailed protocols for the construction of plasmids, phasmids and phages generated in this study
Method S2. Detailed protocol for transposon mutagenesis
Method S3. Detailed protocol for the unmarking of transposon mutants via electroporation of unmarking vectors
Method S4. Detailed protocol for the unmarking of transposon mutants via electroporation of unmarking phages
Method S5. Detailed protocol for the analysis of unmarked transposon mutants
Method S6. Detailed protocol for curing mycobacteria of unmarking plasmids
Method S7. Detailed protocol for the CP-CSeq approach of *Mycobacterium* transposon-tagged mutant libraries

**As separate files on Figshare:**

Data set S1. List of BCG Danish transposon insertion mutants (Excel file: .xls)

Zip file S1. Galaxy workflows and BioPerl algorithm that were used to deconvolute the mutant positions

### Supplementary Figures

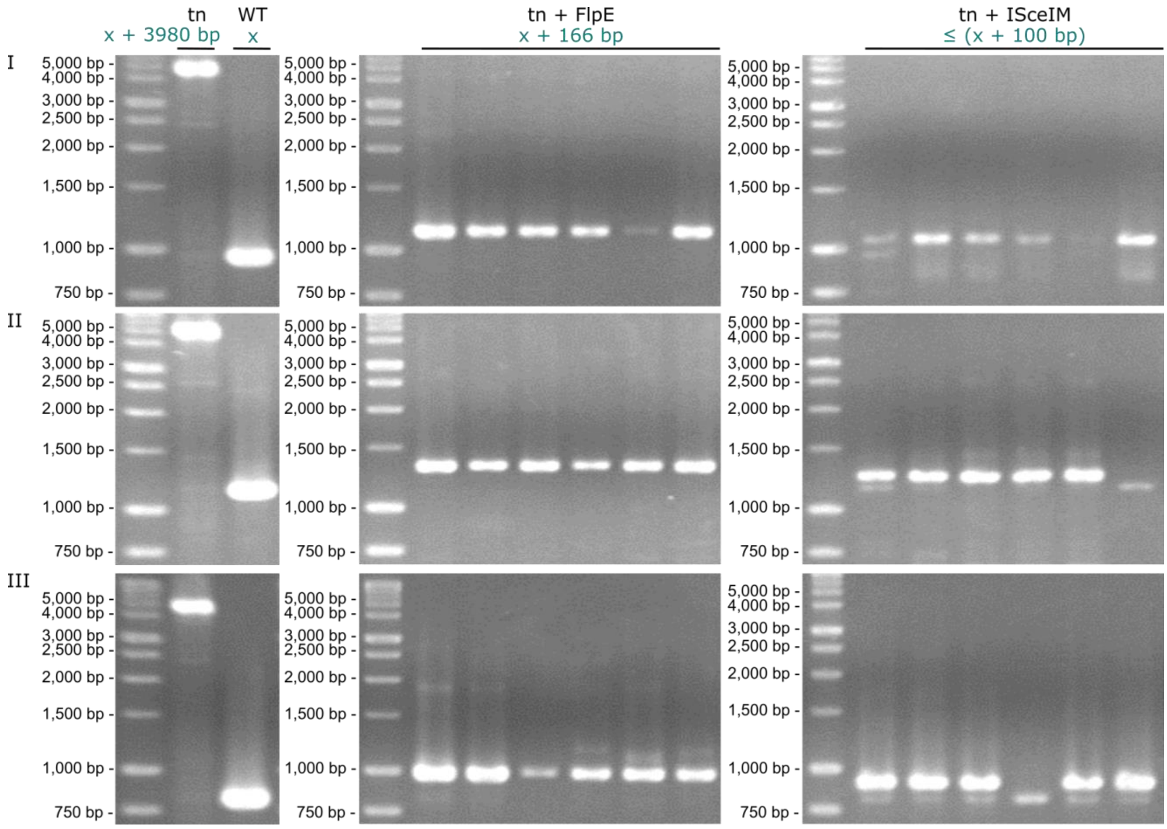

**Figure S1 | Validation of the unmarking of three *M. smegmatis* transposon mutants (I, II and III) via colony-PCR.** Parental transposon mutants (I, II and II) and the original WT strain were taken along as controls. Transposon mutants were unmarked via FlpE/*FRT*-mediated recombination or ISceIM-based transposon removal. Length of expected amplicons are indicated in blue, x: length of WT amplicon, tn: transposon. Used primers are listed in **Table S1.**

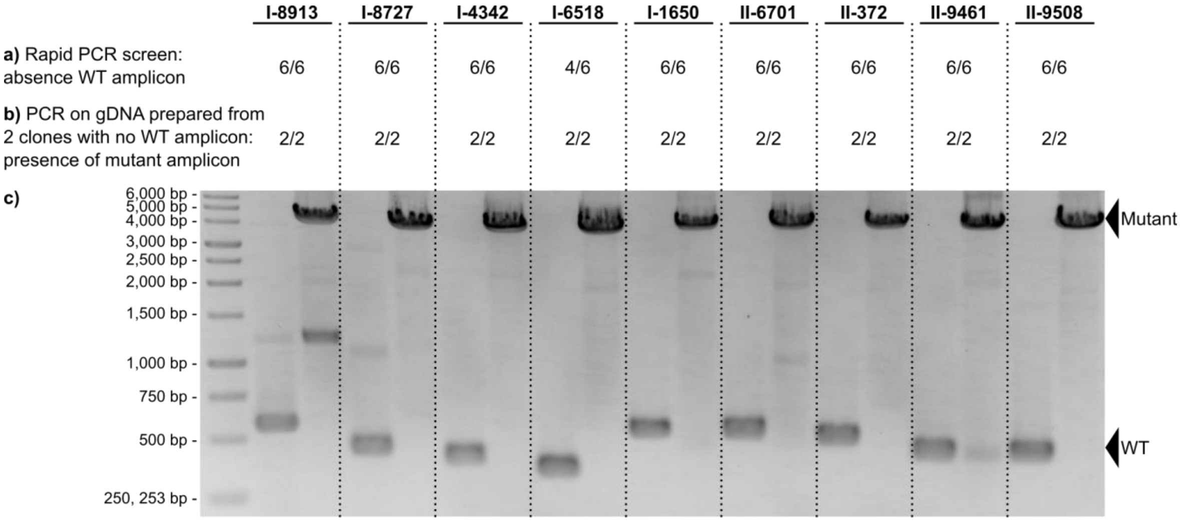

**Figure S2 | Recovering clonally pure cultures for a subset of transposon insertion mutants.** 9 mutants were streaked on agar plates with kanamycin, after which 6 single clones were picked and grown in liquid medium. A rapid PCR screen was performed on supernatant prepared from 50 µl culture (heated to 98°C for 45 minutes and centrifuged for 10 minutes at 3,800 rpm) and WT gDNA as control, to check for absence of a WT amplicon (the large mutant amplicon, which is 3,980 bp larger than the WT amplicon, could not be formed in such a crude PCR). In the figure, the success rate for not detecting a WT amplicon is indicated **(a)**. Subsequently, we prepared, for each mutant, gDNA of two single clones for which no WT amplicon was detected, after which the gDNA was amplified with the same primer set as in the rapid PCR screen (as a control WT gDNA was taken along). For each clone (two per mutant), we could successfully detect the mutant amplicon, while the WT amplicon was absent, indicating that the clones were clonally pure **(b)**. The gel shows for each transposon insertion mutant, the WT amplicon (performed on WT gDNA) and the mutant amplicon (performed on the transposon mutant gDNA) **(c)**. The used primer sets can be found in **Table S6.** The primers hybridize up- and downstream of the investigated transposon insertion event. The unique mutant ID is indicated. By performing the rapid PCR screen we could reduce the necessary gDNA preparations from 54 to 18.

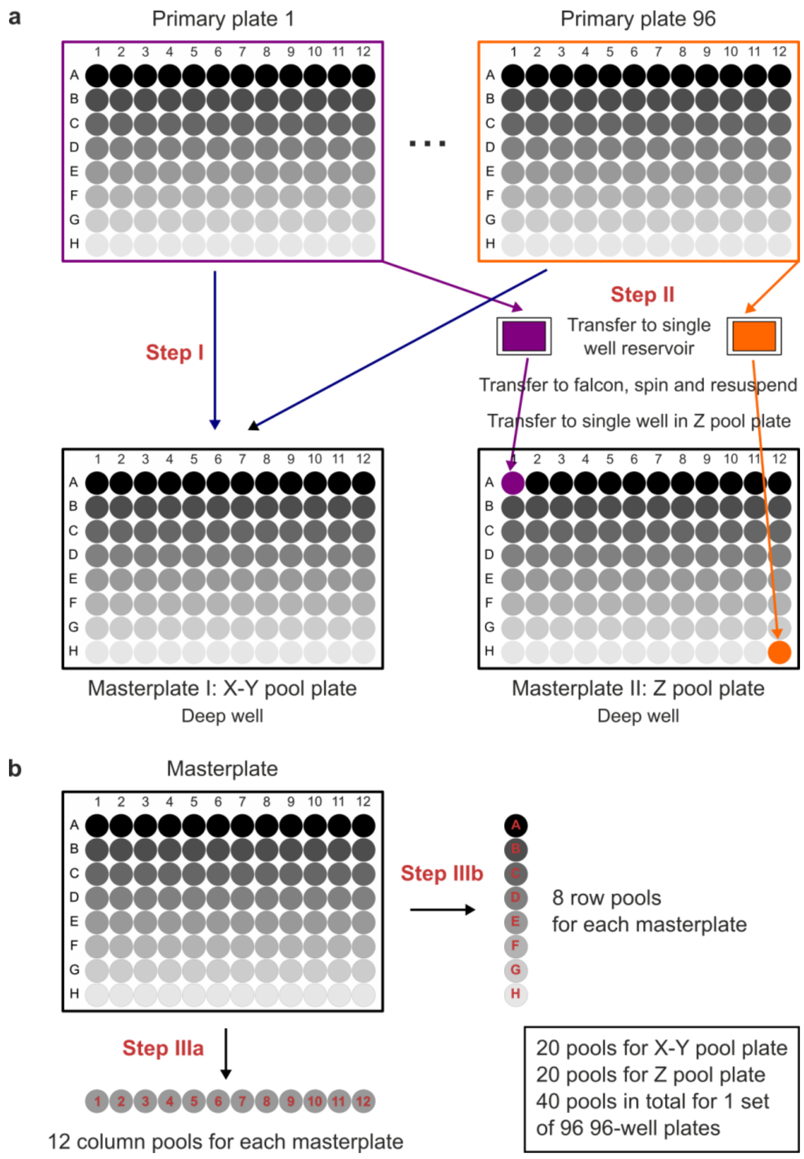

**Figure S3 | Practical implementation of Cartesian pooling using a 96-channel pipette. a)** X-Y coordinate pooling (step I) and Z coordinate pooling (step II) **b)** Masterplate pooling along the rows (step IIIa) and along the columns (step IIIb) creating 20 pools for the X-Y pool plate and 20 pools for the Z pool plate, so 40 pools in total. Figure adapted from Vandewalle *et al*. 2015 (1).

### Supplementary Tables

**Table S1 | List of PCR and Sanger sequencing primers to analyze the unmarking of 3 transposon mutants for *M. smegmatis* and *M. bovis* BCG Danish.** Primer sequences are listed in **Table S5**. The length of transposon (Tn) amplicon is the length of WT amplicon + 3980 bp, the length of FRT amplicon is the length of WT amplicon + 166 bp.

| Tn mutants | | Targeted region | PCR primers | | Length PCR amplicon | | | Sequencing primers | |
|---|---|---|---|---|---|---|---|---|---|
| | | | Fw | Rv | Tn | WT | FRT | Fw | Rv |
| *M. smegmatis* | I | MSMEI_3493 | KB176 | KB177 | 4933 | 953 | 1119 | KB176 | KB177 |
| | II | echA12 | KB179 | KB182 | 5120 | 1140 | 1306 | KB180 | KB181 |
| | III | MSMEI_3898 | KB184 | KB185 | 4787 | 807 | 973 | KB184 | KB185 |
| *M. bovis* BCG Danish | I | BCG_3955c | KB211 | KB212 | 5077 | 1097 | 1263 | KB211 | KB212 |
| | II | lipD | KB215 | KB216 | 4990 | 1010 | 1176 | KB215 | KB216 |
| | III | BCG_0186c-PE1 | KB218 | KB221 | 5149 | 1169 | 1335 | KB219 | KB220 |

**Table S2 | Non-mutated or untraceable non-essential genes are shorter and contain few TA's than mutated and traceable non-essential genes.** Gene length and number (#) of TA for mutated and traceable non-essential genes or non-mutated or untraceable non-essential genes. Non-essential genes that are duplicated in the genome where excluded from the analysis. For each parameter the median, minimum and maximum value is indicated.

| **Non-essential genes** (excluding duplicated genes) (3189 genes) | | median | minimum | maximum |
|---|---|---|---|---|
| Mutated and traceable (2,818 genes) | gene length (bp) | 862.5 | 105 | 12,456 |
| | # TA | 14 | 1 | 187 |
| Non-mutated or untraceable (371 genes) | gene length (bp) | 450 | 71 | 3,696 |
| | # TA | 6 | 0 | 46 |

**Table S3 | List of primers for positional validation (library subset).** Fw primers are listed. Rv primer (IR primer), KB_LIB_000 (5' -> 3': CGGGGACTTATCAGCCAACCTGT).

| Position | Tn insertion | Locus | Name | Location assignment | TA coverage | Fw primer | Fw primer sequence (5' -> 3') | Amplicon |
|---|---|---|---|---|---|---|---|---|
| I-1-H1 | I-7829 | BCGDan_3143 | adhD | Unique | >1200 | KB_LIB_001 | GCGACACCGCCGTCATCTAC | 428 |
| I-1-B2 | I-5526 | . | . | Unique | >1200 | KB_LIB_002 | GATCCGCGCGCTCAAATAGTG | 318 |
| I-1-C2 | I-8913 | BCGDan_3590 | BCGDan_3590 | Unique | >1200 | KB_LIB_003 | GCAGCGCATCGAATCTACCG | 473 |
| I-1-D2 | I-8727 | BCGDan_3510 | BCGDan_3510 | Unique | >1200 | KB_LIB_004 | GCACCTGGGTTCGGATTTGC | 424 |
| I-1-E2 | I-7046 | BCGDan_2868 | BCGDan_2868 | Unique | >1200 | KB_LIB_005 | GGAGGCTGGCGAGGTAGGTG | 369 |
| I-1-F3 | I-2547 | BCGDan_1068 | lpqU | Unique | >1200 | KB_LIB_006 | GTCGCCGAGGTCGAGAATCC | 444 |
| I-1-G3 | I-4342 | BCGDan_1785 | BCGDan_1785 | Unique | >1200 | KB_LIB_007 | GACGGGAATGTCGCGAGGAG | 353 |
| I-1-A5 | I-6518 | BCGDan_2620 | BCGDan_2620 | Unique | 300-1200 | KB_LIB_008 | ACGGCCATCACGTGCAGAAC | 259 |
| I-1-B5 | I-9251 | . | . | Zok_unique | >1200 | KB_LIB_009 | ATCACCCGGAAACGCTGCTG | 301 |
| I-1-C6 | I-1650 | BCGDan_0682 | BCGDan_0682 | Unique | >1200 | KB_LIB_010 | CCTCGACGGGCGTTATCTGC | 436 |
| I-1-E6 | I-4675 | BCGDan_1875 | BCGDan_1875 | Heuristic | >1200 | KB_LIB_011 | GCCGACTTCCAGCGGTTCTC | 294 |
| I-1-C8 | I-5082 | BCGDan_2001 | BCGDan_2001 | Unique | >1200 | KB_LIB_012 | TTCTTGCCACCTGCGCATTC | 293 |

| | | | | | | | | |
|---|---|---|---|---|---|---|---|---|
| I-1-D8 | I-1426 | BCGDan_0585 | BCGDan_0585 | Unique | >1200 | KB_LIB_013 | CGGCGTGAAGTGGGTTTCG | 451 |
| I-1-D9 | I-4296 | . | . | Unique | >1200 | KB_LIB_014 | GATCAGCCACGGTCCGAAGG | 405 |
| I-1-H10 | I-2306 | BCGDan_0968 | pstA1 | Heuristic | >1200 | KB_LIB_015 | GCGTGCTGCCAGAGCAATTC | 258 |
| I-1-B11 | I-4302 | BCGDan_1774 | narX | Zok_unique | >1200 | KB_LIB_016 | GCCGGTAGATCAGCAGGGTAACG | 320 |
| I-1-A12 | I-3549 | BCGDan_1466 | fmu | Unique | >1200 | KB_LIB_017 | GGTCGGTATTCGCCGTTTGC | 457 |
| I-1-D12 | I-3153 | BCGDan_1292 | corA | Unique | >1200 | KB_LIB_018 | TCCAGCTCGGGCATGAAGTG | 354 |
| I-65-B1 | I-7133 | BCGDan_2926 | BCGDan_2926 | Unique | >1200 | KB_LIB_019 | GTCAGGCCCGCGACATAAGC | 294 |
| I-65-C1 | I-4520 | BCGDan_1831 | PE19 | Heuristic | >1200 | KB_LIB_020 | TCCCGACGGGGCTTTATCTG | 444 |
| I-65-D1 | I-8709 | BCGDan_3502 | BCGDan_3502 | Unique | >1200 | KB_LIB_021 | CGCAATCAGCGTGCTTACCG | 445 |
| I-65-E1 | I-1108 | BCGDan_0450 | pssA | Unique | >1200 | KB_LIB_022 | CGTTGCTTGGGCTTGTCGTC | 257 |
| I-65-C4 | I-1771 | BCGDan_0740 | atsA | Unique | >1200 | KB_LIB_023 | CCGGCAAGACCACCCAGTTC | 347 |
| I-65-D4 | I-6935 | BCGDan_2819 | BCGDan_2819 | Unique | >1200 | KB_LIB_024 | ACTCCGCGAATCCACAGCAG | 294 |
| I-65-E4 | I-880 | . | . | Unique | >1200 | KB_LIB_025 | ACGCGTCATTCCGGACTTGG | 401 |
| I-65-F4 | I-5503 | BCGDan_2148 | metH | Zok_unique | >1200 | KB_LIB_026 | TGCGGGATTGAGACCAAACG | 373 |
| I-65-G4 | I-7471 | BCGDan_2995 | pks1 | Unique | >1200 | KB_LIB_027 | CGGTCAACCCGTACCCTTGC | 275 |
| I-65-B7 | I-3079 | BCGDan_1264 | BCGDan_1264 | Unique | >1200 | KB_LIB_028 | TTGGCAGTTCAATGTCGGGTATG | 294 |
| I-65-C7 | I-1296 | BCGDan_0511 | BCGDan_0511 | Unique | >1200 | KB_LIB_029 | CATCCTCGACGGCGTCTTCC | 409 |
| I-65-D7 | I-6264 | BCGDan_2505 | pepN | Unique | >1200 | KB_LIB_030 | CGGTGGACGGCGAGACCTAC | 338 |
| I-65-F8 | I-4073 | BCGDan_1671 | BCGDan_1671 | Unique | >1200 | KB_LIB_031 | CGATGGTGACGACGCAAACC | 367 |
| I-65-G8 | I-6385 | . | . | Unique | >1200 | KB_LIB_032 | CTGGCTGCGTTTAGGTCTCACG | 162 |
| I-65-H8 | I-3429 | . | . | Unique | >1200 | KB_LIB_033 | CTGCCGAACGGTATGCATGG | 286 |
| I-65-D11 | I-2480 | BCGDan_1041 | BCGDan_1041 | Unique | >1200 | KB_LIB_034 | TTGGTAGCGCGACTCGTTCG | 374 |
| I-65-E11 | I-6580 | BCGDan_2641 | BCGDan_2641 | Unique | >1200 | KB_LIB_035 | CGGTGATCGCACCAGACACC | 314 |
| I-65-G11 | I-7374 | BCGDan_2987 | drrC | Zok_unique | >1200 | KB_LIB_036 | CGACGTTGACGTGGTTGTTCG | 334 |
| I-65-B3 | I-2272 | BCGDan_0954 | betP | Unique | 300-1200 | KB_LIB_073 | CGAGCCGCTCAGCCACTACC | 483 |
| I-65-B3 | I-4914 | BCGDan_1949 | aao | Unique | >1200 | KB_LIB_074 | CCGCTGTATGCAATGGTCGAG | 350 |
| I-65-B3 | I-7610 | . | . | Unique | >1200 | KB_LIB_075 | TAGTCCATGAGCGCCCAGACC | 294 |
| I-65-H5 | I-3346 | BCGDan_0382 | BCGDan_0382 | Unique | 300-1200 | KB_LIB_076 | AGCATGCCACTGCCGAATTG | 273 |
| I-65-H5 | I-958 | BCGDan_1387 | glgP | Unique | 300-1200 | KB_LIB_077 | TCCTTCTTGCGCCCGATACC | 415 |
| I-65-D6 | I-5860 | . | . | Unique | 300-1200 | KB_LIB_078 | GTGCACGTCGCCTCCTTGAC | 303 |
| I-65-D6 | I-7331 | BCGDan_2983 | ppsD | Unique | 300-1200 | KB_LIB_079 | GCACCCAACATCGTGATCAACC | 276 |
| I-65-D6 | I-8749 | BCGDan_3521 | BCGDan_3521 | Unique | >1200 | KB_LIB_080 | CCAGCCTTGGAGCGATCCTG | 416 |
| I-65-G7 | I-2472 | . | . | Unique | >1200 | KB_LIB_081 | GCGTTGGGTCTCGACCACTG | 340 |
| I-65-G7 | I-4545 | . | . | Zok_unique | >1200 | KB_LIB_082 | GCAGGTGGCTAACCGCAACC | 368 |
| I-65-E9 | I-9833 | . | . | Unique | 300-1200 | KB_LIB_083 | CGGCGAGCTCCGAGTCTAGTG | 286 |
| I-65-F10 | I-2764 | . | . | Heuristic | 300-1200 | KB_LIB_084 | GGCTTGGAGGCCTGCTGTTC | 261 |
| I-65-F10 | I-6639 | BCGDan_2677 | BCGDan_2677 | Zok_unique | 300-1200 | KB_LIB_085 | GGTGGCCCAAGTGCCCTAAG | 351 |
| I-65-C11 | I-2803 | BCGDan_1165 | BCGDan_1165 | Unique | >1200 | KB_LIB_086 | TGTATCTGCCGATGGATGTGGAC | 297 |
| I-65-C11 | I-4910 | BCGDan_1948 | BCGDan_1948 | Unique | >1200 | KB_LIB_087 | ACCTCCCGCCCTCCAATACG | 397 |

| | | | | | | | | |
|---|---|---|---|---|---|---|---|---|
| I-65-B12 | I-813 | BCGDan_0320 | BCGDan_0320 | Unique | 300-1200 | KB_LIB_088 | CCGTCATACCCGTTCGTCAGC | 452 |
| I-65-E12 | I-2693 | BCGDan_1118 | echA8 | Unique | >1200 | KB_LIB_089 | CGCCTTAGCCTTGCCGATAGC | 481 |
| I-65-E12 | I-5394 | BCGDan_2119 | helY | Unique | 300-1200 | KB_LIB_090 | TTCGCGACCAGCGGTAGATG | 276 |
| II-33-G1 | II-10396 | BCGDan_4004 | ethR | Unique | >1200 | KB_LIB_037 | TGCTGCTGACCCTGCTGGAC | 415 |
| II-33-H1 | II-6701 | BCGDan_2522 | lipQ | Heuristic | >1200 | KB_LIB_038 | CCGTCGCCCGTAGATCACAC | 388 |
| II-33-B3 | II-520 | BCGDan_0187 | BCGDan_0187 | Zok_unique | >1200 | KB_LIB_039 | CGCATCCGCTGTGGTTATGG | 314 |
| II-33-C3 | II-4392 | BCGDan_1698 | pks7 | Unique | >1200 | KB_LIB_040 | CGGGTTCTTGACGGATGTCG | 256 |
| II-33-D4 | II-2846 | BCGDan_1123 | BCGDan_1123 | Unique | >1200 | KB_LIB_041 | TCATGATTACCGCCGCATCC | 301 |
| II-33-F5 | II-2649 | . | . | Unique | >1200 | KB_LIB_042 | GCAGCGGCGCAAGACTTCTC | 290 |
| II-33-G5 | II-5660 | BCGDan_2103 | BCGDan_2103 | Unique | >1200 | KB_LIB_043 | TAGTCGAGCGGCCTGCACTG | 475 |
| II-33-A6 | II-1922 | . | . | Unique | >1200 | KB_LIB_044 | TCGGCTATCTGGGGCATGTG | 380 |
| II-33-B6 | II-9985 | BCGDan_3862 | BCGDan_3862 | Unique | >1200 | KB_LIB_045 | CTATGGCGCTCGCCTGGAAC | 262 |
| II-33-C6 | II-372 | BCGDan_0154 | BCGDan_0154 | Unique | >1200 | KB_LIB_046 | CCAGCGGCATCCAGAATTGAG | 405 |
| II-33-D6 | II-9461 | BCGDan_3590 | BCGDan_3590 | Unique | >1200 | KB_LIB_047 | CAGTCGTTGATCCGCCATTGTC | 252 |
| II-33-H7 | II-9508 | BCGDan_3619 | BCGDan_3619 | Unique | >1200 | KB_LIB_048 | TCAACGCACCAACGTGCATC | 401 |
| II-33-H7 | II-10305 | BCGDan_3971 | pks2 | Unique | 300-1200 | KB_LIB_049 | CGACTTCGGCCAGCATTGTG | 428 |
| II-33-E9 | II-4149 | BCGDan_1606 | mmpL6 | Unique | >1200 | KB_LIB_050 | CGACCCTGCCGTGGTCATAC | 484 |
| II-33-F9 | II-5662 | . | . | Unique | >1200 | KB_LIB_051 | AAGGCCCTGCGATTGACTGG | 478 |
| II-33-B10 | II-6344 | BCGDan_2385 | PPE71 | Unique | >1200 | KB_LIB_052 | GCCGTCGACCTAGGCTTCTTCC | 431 |
| II-33-C10 | II-8204 | BCGDan_3104 | BCGDan_3104 | Unique | >1200 | KB_LIB_053 | CAGCCGCCACTGTTCCACAC | 333 |
| II-33-A12 | II-3170 | BCGDan_1233 | BCGDan_1233 | Unique | >1200 | KB_LIB_054 | CGCCGTCGGGTAGTGGAATC | 483 |
| II-81-C2 | II-10529 | BCGDan_4043 | BCGDan_4043 | Unique | >1200 | KB_LIB_055 | GTGTCAACTGGCCCGTGTGTG | 355 |
| II-81-D2 | II-8287 | BCGDan_3134 | BCGDan_3134 | Unique | >1200 | KB_LIB_056 | AGGGTGCCGTTCGTCATTGC | 255 |
| II-81-E2 | II-5713 | BCGDan_2124 | BCGDan_2124 | Unique | >1200 | KB_LIB_057 | TTGAGCATGGTGGTGGTCTCG | 372 |
| II-81-F2 | II-1730 | . | . | Unique | >1200 | KB_LIB_058 | AATTCCCGCGACAGCGACTC | 410 |
| II-81-G2 | II-7433 | BCGDan_2854 | BCGDan_2854 | Unique | >1200 | KB_LIB_059 | GCGACGGACCCGTTGGATAC | 327 |
| II-81-H2 | II-2654 | BCGDan_1058 | pks16 | Unique | >1200 | KB_LIB_060 | CCAAGGCCGTGATCGTCTCC | 467 |
| II-81-A4 | II-731 | BCGDan_0263 | nirB | Unique | >1200 | KB_LIB_061 | TGCCTACGGCTGGGAGTTCG | 282 |
| II-81-B4 | II-3766 | BCGDan_1469 | BCGDan_1469 | Zok_unique | >1200 | KB_LIB_062 | CGATCGCGGTGATGGTGAATATC | 414 |
| II-81-D6 | II-3714 | . | . | Zok_unique | >1200 | KB_LIB_063 | CCCGATCTCAGCGCCTTCAC | 263 |
| II-81-E6 | II-1462 | . | . | Unique | >1200 | KB_LIB_064 | GTCGATCTCGCCTCGGATGG | 336 |
| II-81-G7 | II-8279 | BCGDan_3134 | BCGDan_3134 | Unique | >1200 | KB_LIB_065 | TTCGACGAGCACGGGATCAG | 384 |
| II-81-G7 | II-8297 | BCGDan_3136 | BCGDan_3136 | Unique | >1200 | KB_LIB_066 | GCGGAGGATAGGGCTTTGTCAC | 441 |
| II-81-A9 | II-8091 | BCGDan_3041 | BCGDan_3041 | Unique | >1200 | KB_LIB_067 | TATATCGCCGCGAGGGAACG | 463 |
| II-81-F10 | II-2884 | . | . | Unique | >1200 | KB_LIB_068 | GCTGGAGCGGATGGACCAAG | 318 |
| II-81-G11 | II-3837 | BCGDan_1500 | PE_PGRS26 | Unique | >1200 | KB_LIB_069 | GTTGGCACCGCTGTTAGGACTG | 307 |
| II-81-H11 | II-1406 | BCGDan_0511 | BCGDan_0511 | Unique | >1200 | KB_LIB_070 | GCGTTAGGCCCGTTGATTGG | 408 |
| II-81-C12 | II-598 | BCGDan_0208 | BCGDan_0208 | Unique | >1200 | KB_LIB_071 | CGGTCAGCCCGTCGATAACC | 281 |
| II-81-D12 | II-10290 | BCGDan_3970 | papA1 | Zok_unique | >1200 | KB_LIB_072 | TATCGGCGCCGGTCAACTC | 438 |

**Table S4 | List of plasmids, shuttle phasmids and phages used in this study**. The genotype, reference and source for each plasmid, shuttle phasmid and phage are indicated. oriE: origin of replication for *E. coli*, oriM: origin of replication for mycobacteria, ts: temperature sensitive, P: promoter, MCS: multiple cloning site, GalK: galactokinase, FlpE: FlpE recombinase, ISceIM: I-SceI meganuclease, MOP: Mycobacterial Optimized Promoter. Addgene or BCCM/GeneCorner (LMBP) accession numbers are indicated were possible.

| Type | Name | Genotype | Reference | Source (and accession number) |
|---|---|---|---|---|
| Plasmid | pMV261Kan | oriE, oriM, KanR, PgroEL, MCS | (2) | Prof. Dr. W.R. Jacobs Jr. |
| | pMyC | oriE, oriM, HygR, Pacetamidase, MCS | Dr. A. Geerlof (unpublished) | Dr. A. Parret & Prof. Dr. M. Wilmanns (Addgene #42192) |
| | pWM18 | oriE, ts oriM, HygR, GmR, SacB, MCS | (3) | Prof. Dr. C. Guilhot |
| | pWM18- | oriE, ts oriM, HygR, MCS | This study | Prof. Dr. N. Callewaert (LMBP 11290) |
| | pDB88 | oriE, HygR, MOP-GalK, SacB, 3’ ligD - 5’ ligD | (4) | Prof. Dr. M.S. Glickman (Addgene #26958) |
| | pML597 | oriE, oriM, KanR, Pimyc-FlpE | (5) | Prof. Dr. M. Niederweis (Addgene #32235) |
| | pGOAL19 | oriE, oriM, AmpR, marker gene cassette [PacI (HygR, PAg85-LacZ, Phsp60-SacB)] | (6) | Prof. Dr. T. Parish (Addgene #20190) |
| | pAmp_Tn[IR-I-SceI-FRT(KanR-OriE)]_PgroEL-Transposase | oriE, AmpR, λcos, PacI, transposon cassette [IR-I-SceI-FRT(KanR-OriE)], PgroEL-Transposase | This study | Prof. Dr. N. Callewaert (LMBP 11288) |
| | pAmp_Tn[IR-I-SceI-FRT(KanR-OriE-SacB)]_PgroEL-Transposase | oriE, AmpR, λcos, PacI, transposon cassette [IR-I-SceI-FRT(KanR-OriE-SacB)], PgroEL-Transposase | This study | Prof. Dr. N. Callewaert (LMBP 11289) |
| | pMV261Hyg (pMV261Hyg_empty) | oriE, oriM, HygR, PgroEL, MCS | (2) | Prof. Dr. E. Rubin |
| | pMV261Hyg_Pimyc-FlpE | oriE, oriM, HygR, Pimyc-FlpE | This study | Prof. Dr. N. Callewaert (LMBP 11291) |
| | pRGM1 (pMV261Hyg_MOP-ISceIM) | oriE, oriM, HygR, MOP-ISceIM | (7) | Prof. Dr. M.S. Glickman (Addgene #48789) |
| | pMyc_ISceIM (pMyc_Pacetamidase-ISceIM) | oriE, oriM, HygR, Pacetamidase-ISceIM | This study | Prof. Dr. N. Callewaert (LMBP 11292) |
| | pMV261Hyg_TetRTetO-FlpE | oriE, oriM, HygR, FlpE under inducible promoter TetRTetO (based on pWK08-Lx)(8) | This study | Prof. Dr. N. Callewaert (LMBP 11299) |
| | pMV261Hyg_TetRTetO-ISceIM | oriE, oriM, HygR, ISceIM under inducible promoter TetRTetO (based on pWK08-Lx)(8) | This study | Prof. Dr. N. Callewaert (LMBP 11300) |
| | pMV261Hyg_PimycTetR-Pmyc1-FlpE | oriE, oriM, HygR, FlpE under inducible promoter PimycTetR-Pmyc1TetO (based on vector pST-KT)(9) | This study | Prof. Dr. N. Callewaert (LMBP 11301) |
| | pMV261Hyg_PimycTetR-Pmyc1-ISceIM | oriE, oriM, HygR, ISceIM under inducible promoter PimycTetR-Pmyc1TetO (based on vector pST-KT)(9) | This study | Prof. Dr. N. Callewaert (LMBP 11302) |
| | pMV261Hyg_GalK_empty | oriE, oriM, HygR, MOP-GalK, Phsp60, MCS | This study | Prof. Dr. N. Callewaert (LMBP 11283) |
| | pMV261Hyg_GalK_Pimyc-FlpE | oriE, oriM, HygR, MOP-GalK, Pimyc-FlpE | This study | Prof. Dr. N. Callewaert (LMBP 11284) |
| | pMV261Hyg_GalK_MOP-ISceIM | oriE, oriM, HygR, MOP-GalK, MOP-ISceIM | This study | Prof. Dr. N. Callewaert (LMBP 11297) |
| | pMV261Hyg_GalK_TetRTetO-FlpE | oriE, oriM, HygR, MOP-GalK, FlpE under inducible promoter TetRTetO (based on pWK08-Lx)(8) | This study | Prof. Dr. N. Callewaert (LMBP 11303) |
| | pMV261Hyg_GalK_TetRTetO-ISceIM | oriE, oriM, HygR, MOP-GalK, ISceI-M under inducible promoter TetRTetO (based on pWK08-Lx)(8) | This study | Prof. Dr. N. Callewaert (LMBP 11304) |
| | pMV261Hyg_GalK_PimycTetR_PmyctetO-FlpE | oriE, oriM, HygR, MOP-GalK, FlpE under inducible promoter PimycTetR-Pmyc1TetO (based on vector pST-KT)(9) | This study | Prof. Dr. N. Callewaert (LMBP 11305) |

| | | | | |
|---|---|---|---|---|
| | pMV261Hyg_GalK_PimycTetR_PmyctetO-ISceIM | oriE, oriM, HygR, MOP-GalK, ISceI-M under inducible promoter PimycTetR-Pmyc1TetO (based on vector pST-KT)(9) | This study | Prof. Dr. N. Callewaert (LMBP 11306) |
| | ts vector empty | oriE, ts oriM, HygR, MOP-GalK | This study | Prof. Dr. N. Callewaert (LMBP 11307) |
| | ts vector FlpE | oriE, ts oriM, HygR, MOP-GalK, Pimyc-FlpE | This study | Prof. Dr. N. Callewaert (LMBP 11295) |
| | ts vector ISceIM | oriE, ts oriM, HygR, MOP-GalK, MOP-ISceIM | This study | Prof. Dr. N. Callewaert (LMBP 11296) |
| | suicide vector empty | oriE, HygR, λcos, PacI | This study | Prof. Dr. N. Callewaert (LMBP 11298) |
| | suicide vector FlpE | oriE, HygR, λcos, PacI, Pimyc-FlpE | This study | Prof. Dr. N. Callewaert (LMBP 11293) |
| | suicide vector ISceIM | oriE, HygR, λcos, PacI, MOP-ISceIM | This study | Prof. Dr. N. Callewaert (LMBP 11294) |
| Shuttle phasmids | ϕMycoMarT7 phasmid | ts phasmid derived from pMycoMar, C9 Himar1 Transposase, λcos site, transposon cassette [IR, PT7 (oriE, KanR)] | (10) | Prof. Dr. E. Rubin |
| | phAE159 | ts phasmid derived from PH101(11) with Δgp48-gp64 | (12) | Prof. Dr. W.R. Jacobs Jr. |
| | optimized transposon phasmid | phAE159::pAmp_Tn[IR-ISceI-FRT(KanR-OriE-SacB)]_PgroEL-Transposase | This study | Prof. Dr. N. Callewaert (LMBP 11287) |
| | phasmid FlpE | phAE159::suicide vector FlpE (Pimyc-FlpE) | This study | Prof. Dr. N. Callewaert (LMBP 11285) |
| | phasmid ISceIM | phAE159::suicide vector ISceIM (MOP-ISceIM) | This study | Prof. Dr. N. Callewaert (LMBP 11286) |
| Phages | optimized transposon phage | ts phage derived from the optimized transposon phasmid carrying PgroEL-Transposase and the transposon cassette [IR-ISceI-FRT(KanR-OriE-SacB)] | This study | Prof. Dr. N. Callewaert |
| | phage empty (phAE159) | ts phage derived from phAE159 | (12) | Prof. Dr. W.R. Jacobs Jr. |
| | phage FlpE | ts phage derived from phasmid FlpE carrying Pimyc-FlpE | This study | Prof. Dr. N. Callewaert |
| | phage ISceIM | ts phage derived from phasmid ISceIM carrying MOP-ISceIM | This study | Prof. Dr. N. Callewaert |
| | phage resolvase (phAE280) | ts phage derived from phAE159::pYB1672 carrying γδ-resolvase (tnpR) | (12) | Prof. Dr. W.R. Jacobs Jr. |

**Table S5 | List of additional primer sequences.** 5' overhangs are underlined, introduce a Gibson assembly overhang or 1-2 restriction site(s).

| Primer nb | Primer sequence (5' -> 3') | Modification |
|---|---|---|
| 379 | ATCGGCATTTTCTTTTGCGTTTTTATTTGTTAACTGTTAATTGTCC | 5' phosphorylation |
| 382 | TAACTAGCGTACGATCGACTGCCAGG | - |
| 383b | TGCGAAGTGATTCCTCCGGATC | - |
| 387 | AGGAGGGCTGGAAGAAGCAG | - |
| 436 | CCACCATGATATTCGGCAAG | - |
| 450 | GGCAGTGCATGCTTATTCAACATAGTTC | 5' phosphorylated |
| 457 | TACCAGATCTTTAAATCTAGAGGTGACCACAACG | - |
| 451 | CATTTACGGGTCTTGTTGTCGTTGG | 5' phosphorylation |
| 460 | TCAGTGGAACGAAAACTCACGTTAAGG | - |
| 466 | CAAGGGCTGCTAAAGGAAGC | - |
| 471 | ACGTATTACGATCCTGATCAGG | - |
| 472 | GCTCTCGCGAATGCATCTTG | - |

| | | |
|---|---|---|
| 473 | GATGGTCGACCTCGACTCGG | - |
| 474 | GCTATGTCTATGCGCCTAGATGTATTCAGC | - |
| 477 | GTCGCGTTTCAGGTCAAGCTG | - |
| 478 | ACGAGAATCGTTAGCGGCACTATTG | - |
| 498 | TCAGATGCGGCGGTTGATGTAC | 5’ phosphorylated |
| 499 | TGTTTAAACTCTAGAAATATTGGATCGTCGCAC | 5’ phosphorylated |
| 558 | CAATTCGCATGCAGAAAGGAGG | - |
| 559 | GAATTCGGATCCTTATTATTTCAGGAAAGTTTCGGAGGAGATAGTG | - |
| KB080 (NdeI site) | CTTTCTAGAGAATAGGAACTTCCCATATGCCAGCTCCAATCTGGTGTGAATG | - |
| KB081 | GCGACGTGCCAACTAGGTCTCCT | 5’ phosphorylated |
| KB103 | GATCTCTCCGGCTTCACCGAT | - |
| KB142 | ACGTGTTCCGCTTCCTTTAGCAG | - |
| KB143 | GCTATCGCCATGTAAGCCCACTG | - |
| KB144 | AGATGCATTCGCGAGAGCTCG | - |
| KB145 | GCCAAGCTTCCTGCTGAACATC | - |
| KB146 | AGTTCGGTGTAGGTCGTTCGCTC | - |
| KB147 | TCGACGCTCAAGTCAGAGGTGG | - |
| KB149 | ACGATAGTTACCGGATAAGGCGCA | - |
| KB150 | GTTACCAGTGGCTGCTGCCAGT | - |
| KB176 | CAGCCACATCGATGTGCTGTG | - |
| KB177 | CATCTCAGTTGCCGCAGCAG | - |
| KB179 | CTTGAGGACCTGGTGCAGAAGATC | - |
| KB180 | GATAGCGTGCTTGTCCGTGACTG | - |
| KB181 | CTCGATCAACAGCAGCGTGC | - |
| KB182 | GATGCGCATCGTGGTGGTCTTAC | - |
| KB184 | GACACCCTCGCACTTGAGCAC | - |
| KB185 | GTGACTCAGCACACCGACATCTCC | - |
| KB211 | GCCGGATTCTGATCTCCGGTC | - |
| KB212 | CGGCTTACGGCTCTGATCTACG | - |
| KB215 | ACAAGGACGAGGTCATGGACCAC | - |
| KB216 | ACCCAGCTGCGAGAGTGGTTC | - |
| KB218 | CTGCTGGTGCGCAATATCAGC | - |
| KB219 | GCGAGTATCTCAACGCGGTTCC | - |
| KB220 | ACGTTGGCCGACATTACTCAGC | - |
| KB221 | CGGCATGATTGAATTCATCGC | - |
| KB222Fw | GCTCACATGTTCTTTCCTGCGTT | - |
| KB222Rv | AACGCAGGAAAGAACATGTGAGC | - |
| KB223Fw | CGCAGTTCGACATCCTGTGC | - |
| KB223Rv | GCACAGGATGTCGAACTGCG | - |

| | | |
|---|---|---|
| KB224Fw | GCCATTTCCGCTGAATATCGTG | - |
| KB224Rv | CACGATATTCAGCGGAAATGGC | - |
| KB225Fw | GGATGCCACCACAGGCACTAC | - |
| KB227Rv | GTGGACCTCGACGACCTGCAG | - |
| KB228 | CAAGACGTTTCCCGTTGAATATGGC | - |
| KB234 | CACGATATTCAGCGGAAATGGCCATCGAGCCGAGAACGTTATCG | - |
| KB235 | GCTCACATGTTCTTTCCTGCGTTCAAGACGTTTCCCGTTGAATATGGC | - |
| KB236 | GTAGTGCCTGTGGTGGCATCCGTGGACCTCGACGACCTGCAG | - |
| KB237 (NheI-PmeI site) | GTTTAAACGCTAGCGGCTCAGTCGAAAGACTGGGC | 5' phosphorylated |
| KB238 (SpeI site) | ACTAGTATAAAGCCAGGTGAGCCCACCA | 5' phosphorylated |
| KB247 | CCTGACAGGATCCGGAGGAAATGTTATGCCGCAGTTCGACATCCT | - |
| KB248 | CCTGACAGGATCCGGAGGAAATGTTATGCAGAAAGGAGGCCATATGG | - |
| KB249*** | AGAGGTTTAAACAGTATTAAACGCACGATATTCAGCGGAAATGGCTTG | - |
| KB250*** | ATACCCATGCGAAGGAGATATACATATGCCGCAGTTCGACATCCT | - |
| KB251*** | ATACCCATGCGAAGGAGATATACATATGCAGAAAGGAGGCCATATGG | - |
| KB252*** | GGTGCGACGATCCAATATTTACGTACGATATTCAGCGGAAATGGCTTG | - |
| KB261 (SpeI site) | ACTAGTCCGATTCTGGCGGTGAGCTC | 5' phosphorylated |
| KB263 | ATCGGTGAAGCCGGAGAGATC | - |

**Table S6 | List of primers to check clonal purity.** Fw and Rv primers are listed. The length of the transposon amplicon is the length of the WT amplicon plus the length of transposon (3980 bp).

| Position | Tn insertion | Locus | Name | Location-assignment | TA coverage | Fw primer | Fw primer sequence (5' -> 3') | Rv primer | Rv primer sequence (5' -> 3') | WT amplicon |
|---|---|---|---|---|---|---|---|---|---|---|
| I-1-C2 | I-8913 | BCGDan_3590 | eccB4 | Unique | >1200 | KB_LIB_003 | GCAGCGCATCGAATCTACCG | KB_LIB_103 | CCGGTGACACTGGCACAAGC | 571 |
| I-1-D2 | I-8727 | BCGDan_3510 | - | Unique | >1200 | KB_LIB_004 | GCACCTGGGTTCGGATTTGC | KB_LIB_104 | GTTCACGATCGCGCTTCTGC | 443 |
| I-1-G3 | I-4342 | BCGDan_1785 | - | Unique | >1200 | KB_LIB_007 | GACGGGAATGTCGCGAGGAG | KB_LIB_107 | CCGTCGGGGTTTCCGATATG | 395 |
| I-1-A5 | I-6518 | BCGDan_2620 | - | Unique | 300-1200 | KB_LIB_008 | ACGGCCATCACGTGCAGAAC | KB_LIB_108 | TGGGGTCGCCAGTCTGGTATC | 342 |
| I-1-C6 | I-1650 | BCGDan_0682 | - | Unique | >1200 | KB_LIB_010 | CCTCGACGGGCGTTATCTGC | KB_LIB_110 | TCGGTCAGGGAGAAGCTGTGC | 534 |
| II-33-H1 | II-6701 | BCGDan_2522 | lipQ | Heuristic | >1200 | KB_LIB_038 | CCGTCGCCCGTAGATCACAC | KB_LIB_138 | GTGGGTTCGCGAGAACATCG | 535 |
| II-33-C6 | II-372 | BCGDan_0154 | - | Unique | >1200 | KB_LIB_046 | CCAGCGGCATCCAGAATTGAG | KB_LIB_146 | CGTCACGGATCAGCGGATTG | 507 |
| II-33-D6 | II-9461 | BCGDan_3590 | eccB4 | Unique | >1200 | KB_LIB_047 | CAGTCGTTGATCCGCCATTGTC | KB_LIB_147 | TCAGCGGGTACCGATTTCTGC | 450 |
| II-33-H7 | II-9508 | BCGDan_3619 | - | Unique | >1200 | KB_LIB_048 | TCAACGCACCAACGTGCATC | KB_LIB_148 | CCCGAGGTGCTGATCATTTGC | 457 |

**Table S7 | List of gBlock sequences (and one PCR amplicon g00x).** Sequence overlaps between the sequence fragments are indicated by underlining them in the same style.

| gBlock nb | gBlock organization | gBlock sequence (5’ -> 3’) |
|---|---|---|
| g00x | Primer 471_PacI_λcos_AmpR_PgroEL-Transposase_Primer 450 | ACGTATTACGATCCTGATCAGGCGCCTTAATTAAGATCCCTATAGTGAGTCGTATTATGCGGCCGCGAATTCTCGATCGTTTGCGGAAAATTTTCATAAATAGCGAAAAC<br>CCGCGAGGTCGCCGCCCCGTAACAAGGCGGATCGCCGGAAAGGACCCGCAAATGATAATAATTATCCATCCGCTTACAGACAAGCTGTGACCGTCTCCGGGAGCTGCA<br>TGTGTCAGAGGTTTTCACCGTCATCACCGAAACGCGCGAGACGAAAGGGCCTCGTGATACGCCTATTTTTATAGGTTAATGTCATGATAATAATGGTTTCTTAGACGTCA<br>GGTGGCACTTTTCGGGGAAATGTGCGCGGAACCCCTATTTGTTTATTTTTCTAAATACATTCAAATATGTATCCGCTCATGAGACAATAACCCTGATAAATGCTTCAATAA<br>TATTGAAAAAGGAAGAGTATGAGTATTCAACATTTCCGTGTCGCCCTTATTCCCTTTTTTGCGGCATTTTGCCTTCCTGTTTTTGCTCACCCAGAAACGCTGGTGAAAGTA<br>AAAGATGCTGAAGATCAGTTGGGTGCACGAGTGGGTTACATCGAACTGGATCTCAACAGCGGTAAGATCCTTGAGAGTTTTCGCCCCGAAGAACGTTTTCCAATGATG<br>AGCACTTTTAAAGTTCTGCTATGTGGCGCGGTATTATCCCGTATTGACGCCGGGCAAGAGCAACTCGGTCGCCGCATACACTATTCTCAGAATGACTTGGTTGAGTACTC<br>ACCAGTCACAGAAAAGCATCTTACGGATGGCATGACAGTAAGAGAATTATGCAGTGCTGCCATAACCATGAGTGATAACACTGCGGCCAACTTACTTCTGACAACGATC<br>GGAGGACCGAAGGAGCTAACCGCTTTTTTGCACAACATGGGGGATCATGTAACTCGCCTTGATCGTTGGGAACCGGAGCTGAATGAAGCCATACCAAACGACGAGCGT<br>GACACCACGATGCCTGTAGCAATGGCAACAACGTTGCGCAAACTATTAACTGGCGAACTACTTACTCTAGCTTCCCGGCAACAATTAATAGACTGGATGGAGGCGGATA<br>AAGTTGCAGGACCACTTCTGCGCTCGGCCCTTCCGGCTGGCTGGTTTATTGCTGATAAATCTGGAGCCGGTGAGCGTGGGTCTCGCGGTATCATTGCAGCACTGGGGCC<br>AGATGGTAAGCCCTCCCGTATCGTAGTTATCTACACGACGGGGAGTCAGGCAACTATGGATGAACGAAATAGACAGATCGCTGAGATAGGTGCCTCACTGATTAAGCA<br>TTGGTAACTGTCAGACCAAGTTTACTCATATATACTTTAGATTGATTTAAAACTTCATTTTTAATTTAAAAGGATCTAGGTGAAGATCCTTTTTGATAATCTCATGACCAAA<br>ATCCCTTAACGTGAGTTTTCGTTCCACTGAATGTCGACGTAGTTTACCAGATCTTTAAATCTAGAGGTGACCACAACGCGCCCGCTTTGATCGGGGACGTCTGCGGCCGA<br>CCATTTACGGGTCTTGTTGTCGTTGGCGGTCATGGGCCGAACATACTCACCCGGATCGGAGGGCCGAGGACAAGGTCGAACGAGGGGCATGACCCGGTGCGGGGCTT<br>CTTGCACTCGGCATAGGCGAGTGCTAAGAATAACGTTGGCACTCGCGACCGGTGAGTGCTAGGTCGGGACGGTGAGGCCAGGCCCGTCGTCGCAGCGAGTGGCAGCG<br>AGGACAACTTGAGCCGTCCGTCGCGGGCACTGCGCCCGGCCAGCGTAAGTAGCGGGGTTGCCGTCACCCGGTGACCCCCGTTTCATCCCCGATCCGGAGGAATCACTT<br>CGCAATGGAAAAAAAGGAATTTCGTGTTTTGATAAAATACTGTTTTCTGAAGGGAAAAAATACAGTGGAAGCAAAAACTTGGCTTGATAATGAGTTTCCGGACTCTGCC<br>CCAGGGAAATCAACAATAATTGATTGGTATGCAAAATTCAAGCGTGGTGAAATGAGCACGGAGGACGGTGAACGCAGTGGACGCCCGAAAGAGGTGGTTACCGACG<br>AAAACATCAAAAAAATCCACAAAATGATTTTGAATGACCGTAAAATGAAGTTGATCGAGATAGCAGAGGCCTTAAAGATATCAAAGGAACGTGTTGGTCATATCATTCA<br>TCAATATTTGGATATGCGGAAGCTCTGTGCGAAATGGGTGCCGCGCGAGCTCACATTTGACCAAAAACAACGACGTGTTGATGATTCTAAGCGGTGTTTGCAGCTGTTA<br>ACTCGTAATACACCCGAGTTTTTCCGTCGATATGTGACAATGGATGAAACATGGCTCCATCACTACACTCCTGAGTCCAATCGACAGTCGGCTGAGTGGACAGCGACCG<br>GTGAACCGTCTCCGAAGCGTGGAAAGACTCAAAAGTCCGCTGGCAAAGTAATGGCCTCTGTTTTTTGGGATGCGCATGGAATAATTTTTATCGATTATCTTGAGAAGGG<br>AAAAACCATCAACAGTGACTATTATATGGCGTTATTGGAGCGTTTGAAGGTCGAAATCGCGGCAAAACGGCCCCACATGAAGAAGAAAAAAGTGTTGTTCCACCAAGA<br>CAACGCACCGTGCCACAAGTCATTGAGAACGATGGCAAAAATTCATGAATTGGGCTTCGAATTGCTTCCCCACCCGCCGTATTCTCCAGATCTGGCCCCCAGCGACTTTT<br>TCTTGTTCTCAGACCTCAAAAGGATGCTCGCAGGGAAAAAATTTGGCTGCAATGAAGAGGTGATCGCCGAAACTGAGGCCTATTTTGAGGCAAAACCGAAGGAGTACT<br>ACCAAAATGGTATCAAAAAATTGGAAGGTCGTTATAATCGTTGTATCGCTCTTGAAGGGAACTATGTTGAATAAGCATGCACTGCC |
| g007 | IR_I-SceI_FRT_KanR (part 1) | GAATAAGCATGCACTGCCAACTAGCGTACGATTAAGTTGTGTGGAATTGTGAGCGGATAACAAACCTGCAGAATTCGGTACTTAAGGCCTTGACTAGAGTAACAGGTT<br>GGCTGATAAGTCCCCGGTCTCTAGACAGTTACGCTAGGGATAACAGGGTAATATAGCATATGGAAGTTCCTATACTTTCTAGAGAATAGGAACTTCCCATATGTAGCTTC<br>ACGCTGCCGCAAGCACTCAGGGCGCAAGGGCTGCTAAAGGAAGCGGAACACGTAGAAAGCCAGTCCGCAGAAACGGTGCTGACCCCGGATGAATGTCAGCTACTGG<br>GCTATCTGGACAAGGGAAAACGCAAGCGCAAAGAGAAAGCAGGTAGCTTGCAGTGGGCTTACATGGCGATAGCTAGACTGGGCGGTTTTATGGACAGCAAGCGAACC<br>GGAATTGCCAGCTGGGGCGCCCTCTGGTAAGGTTGGGAAGCCCTGCAAAGTAAACTGGATGGCTTTCTTGCCGCCAAGGATCTGATGGCGCAGGGGATCAAGATCTG<br>ATCAAGAGACAGGATGAGGATCGTTTCGCATGATTGAACAAGATGGATTGCACGCAGGTTCTCCGGCCGCTTGGGTGGAGAGGCTATTCGGCTATGACTGGGCACAAC<br>AGACAATCGGCTGCTCTGATGCCGCCGTGTTCCGGCTGTCAGCGCAGGGGCGCCCGGTTCTTTTTGTCAAGACCGACCTGTCCGGTGCCCTGAATGAACTCCAAGACGA<br>GGCAGCGCGGCTATCGTGGCTGGCCACGACGGGCGTTCCTTGCGCAGCTGTGCTCGACGTTGTCACTGAAGCGGGAAGGGACTGGCTGCTATTGGGCGAAGTGCCGG<br>GGCAGGATCTCCTGTCATCTCACCTTGCTCCTGCCGAGAAAGTATCCATCATGGCTGATGCAATGCGGCGGCTGCATACGCTTGATCCGGCTACCTGCCCATTCGACCAC<br>CAAGCGAAACATCGCATCGAGCGAGCACGTACTCGGATGGAAGCCGGTCTTGTCGATCAGGATGATCTGGACGAAGAGCATCAGGGGCTCGCGCCAGCCGAACTGTT |

| | | |
|---|---|---|
| | | CGCCAGGCTCAAGGCGCGGATGCCCGACGGCGAGGATCTCGTCGTGACCCATGGCGATGCCTGCTTGCCGAATATCATGGTGGAAAATGGC |
| g008 | KanR (part 2)_OriE_FRT_I-SceI_IR_PacI | GATGCCCGACGGCGAGGATCTCGTCGTGACCCATGGCGATGCCTGCTTGCCGAATATCATGGTGGAAAATGGCCGCTTTTCTGGATTCATCGACTGTGGCCGGCTGGG<br>TGTGGCGGACCGCTATCAGGACATAGCGTTGGCTACCCGTGATATTGCTGAAGAGCTTGGCGGCGAATGGGCTGACCGCTTCCTCGTGCTTTACGGTATCGCCGCTCCC<br>GATTCGCAGCGCATCGCCTTCTATCGCCTTCTTGACGAGTTCTTCTGAGCGGGACTCTGGGGTACGCGTAATACGACTCACGGCTAGCTGATCACCGCGGCCATGATGG<br>CCGGAAAAGGCCGCGTTGCTGGCGTTTTTCCATAGGCTCCGCCCCCCTGACGAGCATCACAAAAATCGACGCTCAAGTCAGAGGTGGCGAAACCCGACAGGACTATAA<br>AGATACCAGGCGTTTCCCCCTGGAAGCTCCCTCGTGCGCTCTCCTGTTCCGACCCTGCCGCTTACCGGATACCTGTCCGCCTTTCTCCCTTCGGGAAGCGTGGCGCTTTCT<br>CATAGCTCACGCTGTAGGTATCTCAGTTCGGTGTAGGTCGTTCGCTCCAAGCTGGGCTGTGTGCACGAACCCCCCGTTCAGCCCGACCGCTGCGCCTTATCCGGTAACTA<br>TCGTCTTGAGTCCAACCCGGTAAGACACGACTTATCGCCACTGGCAGCAGCCACTGGTAACAGGATTAGCAGAGCGAGGTATGTAGGCGGTGCTACAGAGTTCTTGAA<br>GTGGTGGCCTAACTACGGCTACACTAGAAGAACAGTATTTGGTATCTGCGCTCTGCTGAAGCCAGTTACCTTCGGAAAAAGAGTTGGTAGCTCTTGATCCGGCAAACAA<br>ACCACCGCTGGTAGCGGTGGTTTTTTTGTTTGCAAGCAGCAGATTACGCGCAGAAAAAAAGGATCTCAAGAAGATCCTTTGATCTTTTCTACCAAACAACGGATCCGTTT<br>AAACTTGGCCAAGATGCATTCGCGAGAGCTCGATCAGAAGTTCCTATACTTTCTAGAGAATAGGAACTTCCTATATTACCCTGTTATCCCTAGCGTAACTGTCTAGAGAC<br>CGGGGACTTATCAGCCAACCTGTTATCTAGATACGTATTACGATCCTGATCAGGCGCCTTAATTAAGATCCCTATAGTGAGTCGTATTATGCGGCCGCGAATTCTCGA |
| KB003 | Primer KB222_λcos_PacI_Primer KB224 | GCTCACATGTTCTTTCCTGCGTTCCTTGTTACGGGGCGGCGACCTCGCGGGTTTTCGCTATTTATGAAAATTTTCCGGTTTAAGGCGTTTCCGTTCTTCTTCGTCATAACTT<br>AATGTTTTTATTTAAAATACCCTCTGAAAAGAAAGGAAACGACAGGTGCTGAAAGCGAGCTTTTTGGCCTCTGTCGTTTCCTTTCTCTGTTTTTGTCCGTGGAATGAACAA<br>TGGAAGTCAACAAAAAGCAGAGCTTATCGATGATAAGCGGTCAAACATGAGAATTCGCGGCCGCATAATACGACTCACTATAGGGATCTTAATTAAGGCGGCCATTTCC<br>GCTGAATATCGTG |
| KB004 | Primer KB224_Pimyc inactivated PacI_Primer KB223 | GCCATTTCCGCTGAATATCGTGGAGCTCACCGCCAGAATCGGTGGTTGTGGTGATGTACGTGGCGAACTCCGTTGTAGTGCTTGTGGTGGCATCCGTGGCGCGGCCGC<br>GGTACCAGATCTTTAAATCTAGAAATATTGGATCGTCGCACCGGGTTAAGCCTGGAGTGCGGTGGTGCCTGGTCGGCATTTTCGCAGTCGAGGGCTCTCGTGTAGCCTG<br>GGCGAGTTGCCGACGCAGGCGACCCTCCTGCCACGGATCCTTATATAACAGAAAGGAGGTTAATAATGCCGCAGTTCGACATCCTGTGC |
| KB005 | TetRTetO from pWK08-Lx | TGCGTTTAATACTGTTTAAACCTCTAGATCACGATTCGCTCGAGGTCAATGACGCTGTGCGGGAGTGAGAGAATCCGTCGATCAGTGCTGTCAGGCCGAAATCGAATTC<br>CCGGTCGCGTTCAGGAACGCTCGCATTGCCGACCTCTGCGGTTTGCTCCTCCAGTACGACACCGACGGTGTATCGGCTGACCGCGATGAGCATGCTGATGGCTGTCTGT<br>TCATCGAATCCCTGAGACACGAGAAGCCCGACTTGCTGACCGGAAGCCTCAGCTCCTTCGTCGGTGTCGTCGCCGGATGGTCTGCGGTGGTGCTCAGCGTGGAGACGC<br>GCACCGTCACGGACCGCGTGTAGTGCGTGGCGGAAACTTCTTGCGTTTCGCAAGAGAAATGCGTCCCATTTCTCGTCGGACTCGGGGAAGGAAGCGTGATGCTCTCGG<br>TCAAGCACGTCGCTCGCCAGCGCTGCGAGGAGTTCGGCCTTCGTGCGGAAGTGCCAGTAGAGGCCGGGCTGCTGTACCTGTAAGTGAGCCGCCAGCGCGCGAGTGGT<br>GAAGCCATCGAGCCCAGTCTCGTCGAGCACCTGCCGGGCCCCGAGCAACACGGACGTGCGGTCGAGACGCTTCCGGTGGTGAGTCATAGTTGCACTTTATCATCGATA<br>ACTTTATCTTAGATAAAGTGACTGCTCGCTACTCTCATCTGACTGCTCGCTACTCTCATCGTGGAATCCTGACAGGATCCGGAGGAAATGTT |
| KB006 | PimycTetR-Pmyc1TetO from pST-KT | TACGTAAATATTGGATCGTCGCACCGGGTTAAGCCTGGAGTGCGGTGGTGCCTGGTCGGCATTTTCGCAGTCGAGGGCTCTCGTGTAGCCTGGGCGAGTTGCCGACGC<br>AGGCGACCCTCCTGCCACGGATCGGAGGAATCACTTCGCAATGTCAAGATTAGATAAAAGTAAAGTGATTAACAGCGCATTAGAGCTGCTTAATGAGGTCGGAATCGA<br>AGGTTTAACAACCCGTAAACTCGCCCAGAAACTTGGTGTAGAGCAGCCTACATTGTATTGGCATGTAAAAAATAAGCGAGCCCTGCTCGACGCCTTAGCCATTGAGATG<br>TTAGATAGGCACCATACTCACTTTTGCCCTTTAGAAGGGGAAAGCTGGCAAGATTTTTTGCGTAATAACGCTAAAAGTTTTAGATGTGCTTTACTAAGTCATCGCGATGG<br>AGCAAAAGTACATTTAGGTACACGGCCTACAGAAAAACAGTATGAAACTCTCGAAAATCAATTAGCCTTTTTATGCCAACAAGGTTTTTCACTAGAGAACGCGTTATATG<br>CACTCAGCGCTGTGGGGCATTTTACCTTAGGTTGCGTATTGGAAGATCAAGAGCATCAAGTCGCTAAAGAAGAAAGGGAAACACCTACTACTGATAGTATGCCGCCATT<br>ATTACGACAAGCTATCGAATTATTTGATCACCAAGGTGCAGAGCCAGCCTTCTTATTCGGCCTTGAATTGATCATCTGCGGATTAGAAAAACAACTTAAATGTGAAAGTG<br>GGTCTTAATGACCATGGGCTACGTAGAGCCCACCAGATCGTAGAACGTCTGCACCGTCACGGCCGTAGAAGGCCATCCTGACGGATGGCCTTTTTGCGTTTAATACTGT<br>TTAAACCTCTAGAAATAGCACCGTCACCGTGGGAGGCGGCACGTTGCCCGACGTGATGATCGGCCGCATCCCCACGGTGCTGCGCAGTGAGCTCTACGCCATCCCGGC<br>GTTGATCTGTGCGTTCGCACGCACAGGCCCGGTGTGAGAAGGGTCTCTGACGAGCGGGAGAACTCCCTATCAGTGATAGAGTTTGTCCTCCCTATCAGTGATAGATAG<br>GCTCTGGGAGTACCCGTGTGTACGACCAGCACGGCATACATCATTTCGACGCCGAGAGATTCGCCGCCCGAAATGAGCACGATACCCATGCGAAGGAGATATACAT |

**Table S8 | List of custom adaptors** (1). Adaptors for the library were made by combining A000d with A00x in equimolar amounts, heating to 95°C for 5 min and slowly decreasing the temperature to 4°C to anneal both fragments (underlined sequences of A000d and A00x anneal). The adaptors contain the standard Illumina P7 sequence, an 8 bp specific barcode that differs between each Cartesian pool sample, a region to which the standard Illumina barcode sequencing primer hybridizes, a 5' phosphate and a 3' ddC (to block 3' polymerase extension and self-ligation) on the short DNA arm, and a 3' T overhang on the long DNA arm.

| Oligo | Pool assignment | Illumina P7 sequence (5' -> 3') | 8bp barcode (5' -> 3') | Adaptor (5' -> 3') | Modification |
|---|---|---|---|---|---|
| A000d | NA | | | GATCGGAAGAGCACAC | /5Phos/, /3ddC/ |
| A001 | XY_A | CAAGCAGAAGACGGCATACGAGAT | ATCACGTT | GTGACTGGAGTTCAGACGTGTGCTCTTCCGATC**T** | none |
| A002 | XY_B | CAAGCAGAAGACGGCATACGAGAT | CGATGTTT | GTGACTGGAGTTCAGACGTGTGCTCTTCCGATC**T** | none |
| A003 | XY_C | CAAGCAGAAGACGGCATACGAGAT | TTAGGCAT | GTGACTGGAGTTCAGACGTGTGCTCTTCCGATC**T** | none |
| A004 | XY_D | CAAGCAGAAGACGGCATACGAGAT | TGACCACT | GTGACTGGAGTTCAGACGTGTGCTCTTCCGATC**T** | none |
| A005 | XY_E | CAAGCAGAAGACGGCATACGAGAT | ACAGTGGT | GTGACTGGAGTTCAGACGTGTGCTCTTCCGATC**T** | none |
| A006 | XY_F | CAAGCAGAAGACGGCATACGAGAT | GCCAATGT | GTGACTGGAGTTCAGACGTGTGCTCTTCCGATC**T** | none |
| A007 | XY_G | CAAGCAGAAGACGGCATACGAGAT | CAGATCTG | GTGACTGGAGTTCAGACGTGTGCTCTTCCGATC**T** | none |
| A008 | XY_H | CAAGCAGAAGACGGCATACGAGAT | ACTTGATG | GTGACTGGAGTTCAGACGTGTGCTCTTCCGATC**T** | none |
| A009 | XY_1 | CAAGCAGAAGACGGCATACGAGAT | GATCAGCG | GTGACTGGAGTTCAGACGTGTGCTCTTCCGATC**T** | none |
| A010 | XY_2 | CAAGCAGAAGACGGCATACGAGAT | TAGCTTGT | GTGACTGGAGTTCAGACGTGTGCTCTTCCGATC**T** | none |
| A011 | XY_3 | CAAGCAGAAGACGGCATACGAGAT | GGCTACAG | GTGACTGGAGTTCAGACGTGTGCTCTTCCGATC**T** | none |
| A012 | XY_4 | CAAGCAGAAGACGGCATACGAGAT | CTTGTACT | GTGACTGGAGTTCAGACGTGTGCTCTTCCGATC**T** | none |
| A013 | XY_5 | CAAGCAGAAGACGGCATACGAGAT | TGGTTGTT | GTGACTGGAGTTCAGACGTGTGCTCTTCCGATC**T** | none |
| A014 | XY_6 | CAAGCAGAAGACGGCATACGAGAT | TCTCGGTT | GTGACTGGAGTTCAGACGTGTGCTCTTCCGATC**T** | none |
| A015 | XY_7 | CAAGCAGAAGACGGCATACGAGAT | TAAGCGTT | GTGACTGGAGTTCAGACGTGTGCTCTTCCGATC**T** | none |
| A016 | XY_8 | CAAGCAGAAGACGGCATACGAGAT | TCCGTCTT | GTGACTGGAGTTCAGACGTGTGCTCTTCCGATC**T** | none |
| A017 | XY_9 | CAAGCAGAAGACGGCATACGAGAT | TGTACCTT | GTGACTGGAGTTCAGACGTGTGCTCTTCCGATC**T** | none |
| A018 | XY_10 | CAAGCAGAAGACGGCATACGAGAT | TTCTGTGT | GTGACTGGAGTTCAGACGTGTGCTCTTCCGATC**T** | none |
| A019 | XY_11 | CAAGCAGAAGACGGCATACGAGAT | TCTGCTGT | GTGACTGGAGTTCAGACGTGTGCTCTTCCGATC**T** | none |
| A020 | XY_12 | CAAGCAGAAGACGGCATACGAGAT | TTGGAGGT | GTGACTGGAGTTCAGACGTGTGCTCTTCCGATC**T** | none |
| A021 | Z_A | CAAGCAGAAGACGGCATACGAGAT | TCGAGCGT | GTGACTGGAGTTCAGACGTGTGCTCTTCCGATC**T** | none |
| A022 | Z_B | CAAGCAGAAGACGGCATACGAGAT | TGATACGT | GTGACTGGAGTTCAGACGTGTGCTCTTCCGATC**T** | none |
| A023 | Z_C | CAAGCAGAAGACGGCATACGAGAT | TGCATAGT | GTGACTGGAGTTCAGACGTGTGCTCTTCCGATC**T** | none |
| A024 | Z_D | CAAGCAGAAGACGGCATACGAGAT | TTGACTCT | GTGACTGGAGTTCAGACGTGTGCTCTTCCGATC**T** | none |
| A025 | Z_E | CAAGCAGAAGACGGCATACGAGAT | TGCGATCT | GTGACTGGAGTTCAGACGTGTGCTCTTCCGATC**T** | none |
| A026 | Z_F | CAAGCAGAAGACGGCATACGAGAT | TTCCTGCT | GTGACTGGAGTTCAGACGTGTGCTCTTCCGATC**T** | none |
| A027 | Z_G | CAAGCAGAAGACGGCATACGAGAT | TAGTGACT | GTGACTGGAGTTCAGACGTGTGCTCTTCCGATC**T** | none |
| A028 | Z_H | CAAGCAGAAGACGGCATACGAGAT | TACAGGAT | GTGACTGGAGTTCAGACGTGTGCTCTTCCGATC**T** | none |
| A029 | Z_1 | CAAGCAGAAGACGGCATACGAGAT | TCCTCAAT | GTGACTGGAGTTCAGACGTGTGCTCTTCCGATC**T** | none |

| | | | | | |
|---|---|---|---|---|---|
| A030 | Z_2 | CAAGCAGAAGACGGCATACGAGAT | TGTGGTTG | GTGACTGGAGTTCAGACGTGTGCTCTTCCGATC**T** | none |
| A031 | Z_3 | CAAGCAGAAGACGGCATACGAGAT | TAGTCTTG | GTGACTGGAGTTCAGACGTGTGCTCTTCCGATC**T** | none |
| A032 | Z_4 | CAAGCAGAAGACGGCATACGAGAT | TTCCATTG | GTGACTGGAGTTCAGACGTGTGCTCTTCCGATC**T** | none |
| A033 | Z_5 | CAAGCAGAAGACGGCATACGAGAT | TCGAAGTG | GTGACTGGAGTTCAGACGTGTGCTCTTCCGATC**T** | none |
| A034 | Z_6 | CAAGCAGAAGACGGCATACGAGAT | TAACGCTG | GTGACTGGAGTTCAGACGTGTGCTCTTCCGATC**T** | none |
| A035 | Z_7 | CAAGCAGAAGACGGCATACGAGAT | TTGGTATG | GTGACTGGAGTTCAGACGTGTGCTCTTCCGATC**T** | none |
| A036 | Z_8 | CAAGCAGAAGACGGCATACGAGAT | TGAACTGG | GTGACTGGAGTTCAGACGTGTGCTCTTCCGATC**T** | none |
| A037 | Z_9 | CAAGCAGAAGACGGCATACGAGAT | TACTTCGG | GTGACTGGAGTTCAGACGTGTGCTCTTCCGATC**T** | none |
| A038 | Z_10 | CAAGCAGAAGACGGCATACGAGAT | TCTCACGG | GTGACTGGAGTTCAGACGTGTGCTCTTCCGATC**T** | none |
| A039 | Z_11 | CAAGCAGAAGACGGCATACGAGAT | TCAGGAGG | GTGACTGGAGTTCAGACGTGTGCTCTTCCGATC**T** | none |
| A040 | Z_12 | CAAGCAGAAGACGGCATACGAGAT | TAAGTTCG | GTGACTGGAGTTCAGACGTGTGCTCTTCCGATC**T** | none |

**Table S9 | List of primers for library preparation** (1). A mixture of primer P5-IR A-D (underlined bases differ between the different primers) was used for the tn junction amplification to optimize the base read diversity during the 1st cycles of Illumina sequencing.

| Primer | 5' overhang (5' -> 3') | Primer specific sequence (5' -> 3') |
|---|---|---|
| P7 | | CAAGCAGAAGACGGCATACG |
| P5-IR-A (TL028-A) | AATGATACGGCGACCACCGAGATCTACACTCTTTCCCTACACGACGCTCTTCCGATCT | GTCTAGAGACCGGGGACTTATCAGC |
| P5-IR-B (TL028-B) | AATGATACGGCGACCACCGAGATCTACACTCTTTCCCTACACGACGCTCTTCCGATCT | AGTCTAGAGACCGGGGACTTATCAGC |
| P5-IR-C (TL028-C) | AATGATACGGCGACCACCGAGATCTACACTCTTTCCCTACACGACGCTCTTCCGATCT | CAGTCTAGAGACCGGGGACTTATCAGC |
| P5-IR-D (TL028-D) | AATGATACGGCGACCACCGAGATCTACACTCTTTCCCTACACGACGCTCTTCCGATCT | TCAAGTCTAGAGACCGGGGACTTATCAGC |

### Supplementary Methods

#### Method S1 | Detailed protocols for the construction of plasmids, phasmids and phages generated in this study

All PCR reactions were performed with Phusion High-Fidelity polymerase (Bioké) and the primers (IDT DNA) are listed in **Table S5**. The primer melting temperature was calculated using an online tool (http://dharmacon.gelifesciences.com/resources/tools-and-calculators/tm-calculator/), and a PCR annealing temperature was used 2-3°C above the lowest melting temperature. In general, the vectors were constructed via classical restriction enzyme cloning or Gibson assembly (NEB). Ordered gBlocks (IDT) are listed in **Table S7**. DNA fragments were dephosphorylated with FastAP (Thermo Fisher Scientific) or phoshorylated with T4 polynucleotide kinase (Promega), where indicated. Ligations were performed with FastLigase (Enzymatics) or NEBLigase (NEB). Assembled vectors were transformed to *E. coli* MC1061 or DH5α by heat shock and plated on LB with appropriate antibiotics. Single clones were then checked by colony PCR using GoTaqGreen (Promega). Positive clones were prepped (Promega PureYield Mini-prep or Machery-Nagel Nucleobond Xtra Midi-prep) and analyzed by restriction enzyme control digest and Sanger sequencing (VIB Genetics Service Facility (http://www.vibgeneticservicefacility.be/)). Purification of PCR or restriction enzyme fragments was done by magnetic bead purification (AMPure XP beads or CleanNA PCR beads).

The **pAmp_Tn[IR-*I-SceI-FRT*(KanR-OriE)]_PgroEL-Transposase** vector was assembled by Gibson assembly using the following fragments: a PCR amplicon g00x (containing *PacI* site, *λcos* site, ampicillin resistance marker and PgroEL transposase gene, PCR was performed with primers 471 and 450 on pTnFRT_groELTransposase, a previously designed vector), gBlock g007 (containing IR, *I-SceI* site, *FRT* site and part 1 of the kanamycin selection marker) and gBlock g008 (containing part 2 of the kanamycin selection marker, OriE, *I-SceI* site, *FRT* site, IR and *PacI* site).

The **pAmp_Tn[IR-*ISceI-FRT*(KanR-OriE-SacB)]_PgroEL-Transposase** vector was constructed by ligation of the PmeI digested and dephosphorylated pAmp_Tn[IR-*I-SceI-FRT*(KanR-OriE)]_PgroEL-Transposase vector with a purified PCR amplicon of the SacB gene (amplified from pGOAL19 with 5' phosphorylated primers 379 and 451).

We performed an MlsI digest on pWM18 to cut out the SacB and GmR gene and ligated the cut vector to form **pWM18-**.

The **pMV261Hyg_Pimyc-FlpE** vector was created by classical restriction enzyme cloning. The recombinase gene was amplified from the pML597 vector (5' phosphorylated primers 498 and 499), digested with XbaI and cloned into the pMV261Hyg backbone (digested with XbaI, HpaI and 5' dephosphorylated).

The **pMyc_ISceIM** plasmid (with ISceIM under the acetamidase promoter) was constructed by classical restriction enzyme cloning. The I-SceI meganuclease ORF was amplified by PCR from the pRGM1 vector with primers 558 (contains SphI site) and 559 (contains *BamHI*), digested with SphI and BamHI and cloned in the pMyc backbone (digested with NcoI and BamHI) to form the pMyc_I-SceI meganuclease plasmid.

To create the **pMV261Hyg_TetRTetO-FlpE** vector a Gibson Assembly (NEB) was performed using the following fragments: PCR amplicon containing FlpE (no promoter), hygromycin selection marker, oriE and oriM (amplified from the pMV261Hyg_FlpE, primers KB247 and KB249***) and gBlock KB005 containing TetRTetO (based on vector pWK08-Lx(8)).

To create the **pMV261Hyg_TetRTetO-ISceIM** vector a Gibson Assembly (NEB) was performed using the following fragments: PCR amplicon 1 (amplified from the pMV261Hyg_MOP-ISceIM (pRGM1), primers KB248 and KB103) and PCR amplicon 2 (amplified from the pMV261Hyg_MOP-ISceIM (pRGM1), primers KB263 and KB249***) these two amplicons together contain ISceIM (no promoter), hygromycin selection marker, oriE and oriM and are assembled together with gBlock KB005 containing TetRTetO (based on vector pWK08-Lx(8)).

To create the **pMV261Hyg_PimycTetR-Pmyc1-FlpE** vector a Gibson Assembly (NEB) was performed using the following fragments: PCR amplicon containing FlpE (no promoter), hygromycin selection marker, oriE and oriM (amplified from the pMV261Hyg_FlpE, primers KB250*** and KB252***) and gBlock KB006 containing PimycTetR-Pmyc1TetO (based on vector pST-KT(9)).

To create the **pMV261Hyg_PimycTetR-Pmyc1-ISceIM** vector a Gibson Assembly (NEB) was performed using the following fragments: PCR amplicon 1 (amplified from the pMV261Hyg_ISceIM (pRGM1), primers KB251*** and KB103) and PCR amplicon 2 (amplified from the pMV261Hyg_ISceIM (pRGM1), primers KB263 and KB252***) these two amplicons together contain ISceIM (no promoter), hygromycin selection marker, oriE and oriM and are assembled together with gBlock KB006 containing PimycTetR-Pmyc1TetO (based on vector pST-KT(9)).

The **pMV261Hyg_GalK_empty**, **pMV261Hyg_GalK_Pimyc-FlpE** and **pMV261Hyg_GalK_MOP-ISceIM** plasmids were created by classical restriction enzyme cloning. The GalK gene was amplified from the pDB88 vector (KB227Rv and KB228), phosphorylated and cloned into the pMV261Hyg_empty, pMV261Hyg_Pimyc-FlpE or pRGM1 (pMV261Hyg_MOP-ISceIM) backbone (digested with PciI, blunted and 5' dephosphorylated).

The **pMV261Hyg_GalK_TetRTetO-FlpE** and **pMV261Hyg_GalK_TetRTetO-ISceIM** plasmids were created by classical restriction enzyme cloning. The GalK gene was amplified from the pDB88 vector (KB227Rv and KB228), phosphorylated and cloned into the pMV261Hyg_TetRTetO-FlpE or pMV261Hyg_TetRTetO-ISceIM backbone (digested with PciI, blunted and 5' dephosphorylated).

The **pMV261Hyg_GalK_PimycTetR_PmyctetO-FlpE** and **pMV261Hyg_GalK_PimycTetR_PmyctetO-ISceIM** plasmid was created by classical restriction enzyme cloning. The GalK gene was amplified from the pDB88 vector (KB227Rv and KB228), phosphorylated and cloned into the pMV261Hyg_PimycTetR_PmyctetO-FlpE or pMV261Hyg_PimycTetR_PmyctetO-ISceIM backbone (digested with PciI, blunted and 5' dephosphorylated).

To create the **ts vector FlpE**, a Gibson Assembly was performed using the following fragments: PCR amplicon containing FlpE, hygromycin selection marker and oriE (amplified from the pMV261Hyg_FlpE vector, primers KB224Fw and KB222Rv), PCR amplicon containing GalK (amplified from the pDB88 vector, primers KB235 and KB236) and PCR amplicon containing tsOriM (amplified from the pWM18- vector, primers KB225Fw and KB234).

To create the **ts vector ISceIM**, a Gibson Assembly was performed using the following fragments: PCR amplicon containing ISceIM, hygromycin selection marker and oriE (amplified from the pMV261Hyg_ISceIM vector (pRGM1), primers KB224Fw and KB222Rv), PCR amplicon containing GalK (amplified from the pDB88 vector, primers KB235 and KB236) and PCR amplicon containing tsOriM (amplified from the pWM18- vector, primers KB225Fw and KB234).

The **ts vector empty** was created by ligating the PCR amplicon made from the ts vector FlpE using phosphorylated primers KB237 and KB238.

To create the **suicide vector FlpE**, a Gibson Assembly was performed using the following fragments: gBlock KB_004 containing the Pimyc promoter with an inactive *PacI* site (primers KB_224Fw and KB_223Rv), PCR amplicon containing FlpE (without promoter), hygromycin selection marker and oriE (amplified from the pMV261Hyg_FlpE vector, primers KB223Fw and KB222Rv) and gBlock KB003 containing the *λcos* site and the *PacI* site (primers KB222Fw and KB224Rv).

To create the **suicide vector ISceIM**, a Gibson Assembly was performed using the following fragments: PCR amplicon containing MOP-ISceIM, hygromycin selection marker and oriE (amplified from the pMV261Hyg_ISceIM vector (pRGM1), primers KB224Fw and KB222Rv) and gBlock KB003 containing the *λcos* site and the *PacI* site (primers KB222Fw and KB224Rv).

The **suicide vector empty** was created by ligating the PCR amplicon made from the suicide vector FlpE using phosphorylated primers KB237 and KB261.

For the production of the **optimized transposon phasmid** or the **phasmids bringing FlpE or ISceIM** to expression; the shuttle phasmid phAE159 and the insert plasmid (pAmp_Tn[IR-ISceI-FRT(OriE-KanR-SacB)]_PgroEL-Transposase or suicide vectors FlpE or ISceIM) were digested with PacI and ligated together in a 1:1 molar ratio. With 10 µl of the ligation mix, a Lambda Packaging reaction was performed, using *E. coli* TOP10 or MC1061 cells and MaxPlaxTM Lambda Packaging Extracts (Epicentre, according to the manufacturer's instructions), after which we plated out the cells on agar plates with kanamycin and proceeded as for normal plasmids.

The **ts phages** were created from the corresponding phasmids, following the protocols for mycobacteriophage generation and propagation from Jain *et al.* 2014 (12).

#### Method S2 | Detailed protocol for transposon mutagenesis

*M. smegmatis* and *M. bovis* BCG were grown to an $OD_{600}$ of 0.8-1.0 in 7H9 broth supplemented with OADC, glycerol and Tween 80. Cells (10 ml per transduction) were centrifuged (at room temperature), washed 3 times with pre-warmed (37°C) MP buffer and resuspended in the same buffer (1/10$^{th}$ or 1/100$^{th}$ of the initial volume). A 200 µl aliquot was removed to serve as a negative control. The washed bacteria and phage stock were pre-warmed to 39°C before mixing them. Next, 0.12 or 1.2 ml of 4-5 x $10^{10}$ PFU (plaque-forming units) warm phage solution, or MP buffer as negative control, was added to the warm bacilli. The transduction mixture was incubated for 4 hours at 37°C. After incubation, the transduction mix was plated on pre-warmed (37°C) selective plates containing 50 µg/ml kanamycin. For *M. smegmatis*, LB plates were used, which were incubated at 37°C for 3 days. For *M. bovis* BCG, 7H10 plates were used, which were incubated at 37°C for 2-4 weeks. The titer of the transduced bacteria was calculated as PFU/ml with the following formula: [(1000 µl)(# colonies)]/[amount plated in µL)(dilution)]

#### Method S3 | Detailed protocol for the unmarking of transposon mutants via electroporation of unmarking plasmids

*M. smegmatis* or *M. bovis* BCG were grown to mid-log phase ($OD_{600}$ of 0.7-0.8) in 7H9 medium supplemented with Tween 80 and ADS. The day before electroporation the cells were subcultured to reach an $OD_{600}$ of 0.7-0.8 the next day, 1.5% glycine was added to the *M. bovis* BCG cultures. On the day of electroporation, cells were centrifuged (at room temperature) and washed 3 times with pre-warmed (37°C) 0.05% Tween 80 (in water). The cells were resuspended in the same buffer (210 µl of buffer per 10 ml of culture and per electroporation), after which the cells were mixed with the plasmid/phasmid (1 µg) in a pre-warmed 2 mm electroporation cuvette and electroporated (2.5 kV, 25 µF and 800 Ω). 1,200 µl of pre-warmed 7H9-ADS-Tween 80 medium was added immediately after electroporation. For *M. smegmatis*, the transformation mix was recovered without antibiotics for 24 hours, plated on selective LB agar plates and incubated for 2-4 days at 37°C. For *M. bovis* BCG, the transformation mix was recovered without antibiotics for 3 days, plated on selective 7H10 agar plates and incubated for 2-4 weeks at 37°C. When working with ts plasmids the agar plates were incubated at 32°C or 39°C. The number of hygromycin-resistant clones was calculated as a measure for the efficiency of transformation. No absolute numbers were reported to compare the different vectors, as the electroporation efficiency differed between the different experiments. Nonetheless, differences in transformation efficiency between the different vectors remained similar, therefore the efficiency of transformation was given a +, ++ or +++.

#### Method S4 | Detailed protocol for the unmarking of transposon mutants via transduction of unmarking phages

*M. smegmatis* and *M. bovis* BCG were grown to an OD600 of 0.8-1.0 in 7H9 broth supplemented with OADC, glycerol and Tween 80. Cells (5 ml per transduction) were centrifuged (at room temperature), washed 3 times with pre-warmed (37°C) MP buffer and resuspended in the same buffer (1/20th of the initial volume or 250 µl). The washed bacteria and phage stock were pre-warmed to 39°C (or not) before mixing them. Next, 500 µl of ~1010 plaque-forming units of warm phage, or MP buffer as negative control, was added to the warm bacilli. The transduction mixture was incubated overnight (*M. smegmatis*) or 3 days (*M. bovis* BCG) at 37°C. After incubation, the transduction mix was plated on pre-warmed (37°C or room temperature) selective plates: for *M. smegmatis*, LB plates were used, which were incubated at 37°C for 3 days, for *M. bovis* BCG, 7H10 plates were used, which were incubated at 37°C for 2-4 weeks.

The transduction protocol described by Jain *et al.* 2014 (12) used to unmark specialized transduction mutants using PhAE280 is similar, but contains differences compared to our protocol. They use 10% sucrose plates instead of 2% sucrose plates. It is possible to use 10% sucrose plates to unmark specialized transduction mutants, but to unmark transposon mutants the use of 2% sucrose plates is highly recommended. In addition, the protocol of Jain *et al.* does not instruct to pre-warm the buffer, cultures, phages and plates. Not pre-heating everything to 37-39°C also proved to be essential to get a higher percentage of unmarked mutants with the unmarking phages. This was of greater importance for the FlpE and ISceIM unmarking phages as the percentage of unmarked mutants for the resolvase unmarking phage (phAE280) is already quite high regardless of pre-heating the samples. To get the highest possible percentage of unmarked mutants, a balance needs to be found between not overheating the samples (not enough lysis of SacB escape mutants) and not cooling down the samples too much (too much lysis of cells).

#### Method S5 | Detailed protocol for the analysis of unmarked transposon mutants

The colonies obtained after unmarking (by electroporating unmarking plasmids or transducing unmarking phages) were analyzed for the presence or absence of resistance markers by streaking the clones on agar plates containing different supplements. Marked transposon mutants are kanamycin resistant and sucrose sensitive, unmarked transposon mutants are kanamycin sensitive and sucrose resistant and SacB escape mutants are kanamycin and sucrose resistant. The percentage of unmarked clones (kanamycin sensitive and sucrose resistant) among the obtained sucrose-resistant colonies was calculated as a measure of the efficiency of unmarking.

Unmarked transposon mutants were subsequently analyzed via colony PCR and/or Sanger sequencing. Therefore, scraped cells from streaked colonies were boiled for 30-45 min at 98°C in 30 µl water. The debris was centrifuged and the supernatant (2-5 µl) was used as a template in a PCR reaction performed with Phusion High Fidelity polymerase. The PCR amplicons were analyzed by running the samples on the fragment analyzer (DNF-935 kit, AATI) or on an (1% or 2%) agarose gel, stained with ethidium bromide or Midori Green and visualized under UV light. For Sanger sequencing, PCR products were purified with DNA binding magnetic beads (CleanPCR beads, CleanNA) and Sanger sequenced with the indicated primers by the VIB Genetics Service Facility (http://www.vibgeneticservicefacility.be/) or Eurofins Genomics (https://www.eurofinsgenomics.eu/en/custom-dna-sequencing/). The primers used to analyze the unmarking of 3 transposon mutants for *M. smegmatis* and *M. bovis* BCG Danish are listed in **Table S1**, the primer sequences in **Table S5**.

#### Method S6 | Detailed protocol for curing mycobacteria of unmarking plasmids

Unmarked mycobacteria carrying an unmarking plasmid, containing a hygromycin positive selection marker and/or a GalK negative selection marker, were inoculated in 7H9 broth and subcultured (once for *M. smegmatis*, twice for *M. bovis* BCG) before making serial dilutions and plating them on agar plates with or without 2-DOG (0.2% for *M. bovis* BCG and 0.5% for *M. smegmatis*). The agar plates were incubated at 37°C for 3 days for *M. smegmatis* or for 2-3 weeks for *M. bovis* BCG. Subsequently, single colonies (at least 20) were streaked on agar plates with or without hygromycin, incubated at 37°C for 2 days (*M. smegmatis*) or ~1 week (*M. bovis* BCG) to check if the unmarking plasmid was cured. The percentage of cured clones (hygromycin sensitive) was calculated as a measure for the efficiency of curing.

### Method S7 | Detailed protocol for the CP-CSeq approach for *Mycobacterium* transposon-tagged mutant libraries

This protocol outlines the detailed CP-CSeq procedure used to generate the transposon library in *M. bovis* BCG Danish 1331, with an emphasis on the Cartesian pooling steps and the subsequent custom Illumina library preparation and sequencing data analysis. This protocol is based on the protocol described by Vandewalle *et al.* 2015 (1), but has been adapted or extended at several points.

#### 1. Special equipment and materials

##### 1.1. Archived mutant library construction

###### a. Special equipment

- Laminar flow cabinets
- BSL1-3 facility (depending on the organism of interest)
- 37°C incubators (to grow the library in 96-well plates, preferably an incubator with humidity control option)
- 96-well multichannel pipette, e.g. Platemaster P220 (Gilson) or Viaflo 96 (Integra)
- Optional Univo Manual Capper CM480 (Micronic), highly recommended when using Matrix Storage Tubes in combination with Sepraseal Mats

###### b. Materials

- Mycobacterial strain of interest
- Phage stock (optimized transposon phage) ($10^{10}$ p.f.u.)
- MP buffer (50 mM of Tris, pH 7.5, 150 mM of NaCl and 2 mM of $CaCl_2$)
- Sterile fine tips (e.g. 10 µl) for picking the clones
- Sterile filter tips (compatible with 96-well multichannel pipette) (e.g. Diamond Tips for Platemaster, DF300ST, blister refill 960, Gilson catalog #F172703)
- Large plates, e.g. BioAssay dishes (square, standard height) (Nunc, VWR catalog 734-2179)
- Flat bottom sterile 96-well tissue culture plates with low evaporation lid (Falcon, VWR catalog #734-0023)
- 96-well plates for storage of *Mycobacterium* mutant library in -80°C, e.g. Matrix Blank and Alphanumeric Storage Tubes racked in Latch Racks (Fisher Brand, Fischer Scientific catalog #10496163) and X960 Sepraseal Mats, Caps, Sterile (Fisher Brand, Fischer Scientific catalog #10233393)
- Medium components
    - Middlebrook mycobacterial growth medium 7H9 (BD catalog #271310), 7H10 (BD catalog #262710), OADC (BD catalog #212351)
    - Antibiotics (kanamycin (Sigma-Aldrich catalog #K4378))
    - Glycerol
    - Tween-80
    - Optional: mycobacterial growth supplements (MD-VS vitamin supplement (ATCC), Casamino Acids (Bacto) and L-Trp (Sigma)

##### 1.2. Cartesian pooling

###### a. Special equipment

- Laminar flow cabinets
- BSL1-3 facility (depending on the organism of interest)
- 37°C incubator (preferably with humidity control option)
- 96-well multichannel pipette, e.g. Platemaster P220 (Gilson) or Viaflo 96 (Integra)

#### b. Materials

- Sterile filter tips (compatible with 96-well multichannel pipette) (e.g. Diamond Tips for Platemaster, DF300ST, blister refill 960, Gilson catalog #F172703)
- 96-well deep well plates (Corning, Sigma-Aldrich catalog #Z717266)
- Single-well plate disposable reservoirs (Viaflo, Elscolab catalog #6312)

### 1.3. Library preparation Coordinate-Sequencing

#### a. Special equipment

- Bioruptor (Diagenode catalog # B01010001) or Covaris DNA shearing instrument
- Eppendorf magnetic stand or 96-well plate magnet (e.g. Thermo Fisher Scientific catalog #12321D or #12331D)
- Nanodrop spectrophotometer
- Qubit 2.0 fluorometer
- Fragment analyzer (AATI, Agilent)

#### b. Materials

- Genomic DNA preparation materials
    - Lysozyme (Sigma catalog #L6876). Prepare 5 mg/ml lysozyme solution in 20 mM Tris/HCl pH 9; Store in small aliquots at -20°C (do not repeatedly freeze and thaw).
    - SDS solution (10%; store at room temperature)
    - Proteïnase K (2 mg/ml; store aliquots at -20°C)
    - Tris/HCl (20 mM, pH 9)
    - NaCl (5 M)
    - CTAB/NaCl (10% CTAB in 0.7M NaCl). Dissolve 4.1 g of NaCl in 80 ml of distilled water. While stirring, add 10 g of CTAB. If necessary, heat solution to 65°C. Adjust volume to 100 ml with distilled water. Store at room temperature for no longer than 6 months.
    - Phenol/Chloroform/Isoamyl alcohol (Thermo Fisher Scientific catalog #15593-031; store at 4°C or at -20°C)
    - Isopropanol
    - 75% ethanol
    - 1x TE (Tris-EDTA) buffer
- TPX eppendorfs for DNA shearing (Diagenode catalog #C30010010-50)
- NEBNext UltraII end repair/dA-tailing module (NEB, Bioké catalog #E7546L)
- T4 DNA ligase (Enzymatics, Westburg catalog #L-6030-HC-L) or NEB T4 DNA ligase (NEB, catalog #M0202)
- GoTaq Green Master Mix (Promega catalog #M7122 or #M7123)
- DNA binding magnetic beads e.g. CleanPCR beads (CleanNA, GC biotech catalog #CPCR-0050)
- 1x TE (Tris-EDTA) buffer
- 1x duplex buffer (100 mM KAc, 30 mM HEPES, pH7.5)
- Custom adaptors (IDT, **Table S8**)
- Qubit dsDNA HS (High Sensitivity) assay kit, 10 pg/µl - 100 ng/µl (Invitrogen, ThermoFisher Scientific catalog #Q32851)
- High Sensitivity NGS Fragment Analysis Kit (AATI, Agilent, catalog #DNF-474)

### 2. Protocol

#### 2.1. Archived mutant library construction

##### a. Transduction and plating out

- Grow 250 ml (10 x 25 ml) of *M. bovis* BCG to $OD_{600}$ = 1.5 (cultures grown to a different $OD_{600}$ can also be used) in 7H9 broth supplemented with 10% OADC, 0.2% glycerol and 0.05% Tween80.
- Wash the cells three times (in 5 aliquots of 50 ml) with MP buffer (pre-warmed to 37°C) and resuspend in 17.5 ml (5 x 3.5 ml) of MP buffer.
- Infect the washed cells with 12.5 ml (5 x 2.5 ml) $10^{10}$ p.f.u. of the optimized transposon phage for 4 h at 37°C.
    - ! Pre-warm the bacteria and phage to 39°C before performing the transduction.
- After transposon mutagenesis, plate out the *Mycobacterium* strain of interest on suitable medium (e.g. 7H10 agar, supplemented with 10% OADC, 0.5% glycerol and 100 µg/ml kanamycin or other appropriate antibiotics, depending on the transposon mutagenesis phage used) using glass beads.
    - ! In addition, we supplemented with 1% MD-VS vitamin supplement (ATCC), 0.2% Casamino Acids (Bacto) and 50 µg/ml L-Trp to allow incorporation of auxotrophic mutants in the library.
    - ! The plating was performed at a 1:2 dilution (in pre-warmed MP buffer) on 60 large agar plates to obtain well-separated single colonies. (800 µl 1:2 dilution on each plate).
    - ! Pre-warm the agar plates at 37°C before, as these big agar plates warm up slowly.
    - ! We wrapped stacks of 4 plates in thin plastic bags. Making bigger stacks or using thick plastic bags is discouraged as this slows the warming up of the plates. Avoid overfilling the incubator.
- Incubate 2-3 weeks at 37°C.
    - ! **We recommend optimizing the transduction protocol before attempting to make a transposon library, since many parameters (like a different BCG strain, different phage stock, personal transduction manner, different medium, big versus small agar plates, …) influence the success rate of transposon mutagenesis and it is the goal to attain enough well-separated single colonies on the agar plates allowing precise picking of the transposon library.**

##### b. Single colony picking into archived mutant library

- Once the colonies are well grown on the agar plates, inoculate single colonies into flat bottom 96-well tissue culture plates with low evaporation lid, prefilled with 250 µl of 7H9 broth (supplemented with 10% OADC, 0.2% glycerol, 0.05% Tween-80, 100 µg/ml kanamycin or other appropriate antibiotics and optionally 1% MD-VS vitamin supplement (ATCC), 0.2% Casamino Acids (Bacto) and 50 µg/ml L-Trp)
    - ! Picking was done manually (1 week with 4 people) using sterile tips to avoid clustered colonies. With appropriate optimization, robotic colony pickers would likely also work.
    - ! We picked a library of two sets of 96 x 96-well plates (18,432 clones), but this can be scaled up or down, with minor modifications to the subsequent steps.
    - ! In our experience, mycobacterial growth is faster in smaller volumes (200 µl or less) in these TC plates compared to higher volumes (1 ml) in deep wells. Moreover, a larger number of the mutants could be successfully cultivated in the lower-volume wells.
    - ! One can visually inspect under the microscope if one is really putting cells in each well. To have adequate growth, a small clump of cells needs to be visible under the microscope (40x magnification)
- Shake plates shortly (450 rpm) to assure the bacteria are well-dispersed in the medium.
- For slow-growing mycobacteria, incubate statically for 2 – 4 weeks at 37°C.

- ! *M. bovis* BCG Danish cultures were incubated 3 weeks at 37°C
- ! We incubated the cultures in a humidity controlled incubator to minimize evaporation.
- ! We made stacks of 6 plates with a water plate (300 µl sterile water) on the top and on the bottom of the stack (making a stack of 8 plates). To avoid evaporation in the top plates.
- ! We have shaken the plates every four days (max 600 rpm) to avoid severe sticking of the cells to the bottom of the plates and to stimulate their growth

##### c. Replicating and storing the library

- Shake plates (max 750 rpm for one plate) until completely homogenous cultures
- Transfer a small volume (50 µl) of each library plate to a fresh plate filled with supplemented medium (200 µl) (1:5 dilution) to replicate the library in triplicate
- Incubate the replicated plates 1-2 weeks at 37°C
    - ! We made stacks of 8 plates with a water plate (300 µl sterile water) on the top and on the bottom of the stack (making a stack of 10 plates). To avoid evaporation in the top plates.
    - ! We incubated the plates in a humidity controlled incubator to minimize evaporation. Due to space-issues we also incubated the plates in a non-humidity controlled incubator, these stacks were wrapped in plastic bags to minimize evaporation.
    - ! No shaking of the plates during this incubation
    - ! We incubated the plates 1 week at 37°C to an $OD_{600}$ of 0.9, but we recommend growing the strains longer. To enable the preparation of more genomic DNA.
- Perform the pooling steps (**Section 2.2**) using one replicate
- Use the other two replicates to make two stocks of the library, which are preferably frozen in two separate -80°C freezers. Transfer 160-180 µl to -80°C compatible storage plates (prefilled with 40-45 µl glycerol; final concentration 20%) for long-term storage.

#### 2.2. Cartesian pooling

The Cartesian pooling concept can be practically implemented in different ways, depending on available liquid handling equipment and manpower. The option that is described here uses a 96-channel pipette (manual or robotic), with which 96 plates can be processed in a day by two persons. One could perform the pooling with one person, but we recommend doing it with two persons to avoid mix-ups between the plates and pools.

The Cartesian pooling concept is developed for one set of 96 96-well plates. Pooling less plates together using this strategy is possible. When one wants to pool more than 96 96-well plates together, one needs to perform the Cartesian pooling separately for the different sets of 96 96-well plates. So 192 96-well plates are seen as two separate sets of 96 96-well plates that are pooled separate from each other, creating 2 x 40 pools.

An illustration of the practical implementation of the Cartesian pooling strategy is depicted in **Fig. S3**.

##### a. X-Y coordinate pooling (step I)

- X-Y coordinate pooling, creating masterplate type I or X-Y pool plate.
- Transfer 20 µl culture volume from each well of primary plate 1 into the corresponding well of a 96-deep well masterplate using a 96-channel pipette. The positions of the wells (X and Y coordinate) remain the same.
- Repeat for each of the 96 primary plates, transferring to one and the same deep well plate.

##### b. Z coordinate pooling (step II)

- Z coordinate pooling, creating masterplate type II or Z pool plate

- Transfer 20 µl of each well of primary plate 1 into a single-well reservoir (filled with 10 ml of 7H9 medium) using a 96-channel pipette. Then, manually pipette or pour this pool into a falcon tube, spin down the cells (4000 rpm 15 min), resuspend in 1.5 ml of 7H9 medium and transfer to well A1 of the Z Pool Plate.
- Repeat for each of the 96 primary plates, transferring each plate to a single well of the Z pool plate

##### c. Masterplate pooling (step III)

- Create column and row pools for the two masterplates: X-Y and Z pool plates. You will obtain 12 column pools and 8 row pools for each master plate. So for a complete library of 96 96-well plate you will end up with 20 pools for each master plate or 40 pools in total.
    - ! Change tips in between changing rows to avoid cross-contaminations.
    - ! Be careful while pipetting. Put in your tip carefully/slowly to avoid overflow
    - ! Pipet solutions up and down before taking a fraction out
    - ! Take fresh 1 ml tip boxes, to be able to follow what you have been doing
- Column pools **(step IIIa)**: transfer 3/5 (with a ≥5% margin) of the volume of every well of a certain column in a 15 ml falcon corresponding to the appropriate column pool
- Row pools **(step IIIb)**: transfer 2/5 (with a ≥5% margin) of the volume of every well of a certain row in a 15 ml falcon corresponding to the appropriate row pool
- Optionally: mix pools well (vortex) and divide in two fractions to have a spare fraction
- Centrifuge (4000 rpm 15 min)
- Remove supernatant and either continue with genomic DNA preparation (**Section 2.3**) or store the bacterial pellet in the freezer.

#### 2.3. Library preparation Coordinate-Sequencing

The 40 pooled samples obtained from Cartesian pooling are subsequently processed for Illumina sequencing. Treat these samples independently during the entire library preparation protocol up until the final step. Use an aliquot of each pooled sample for library preparation and store the rest in the freezer as a back-up.

The enzymes, kits and devices mentioned below are the ones we used. Nevertheless, with minor modifications, kits from other suppliers may function equally well.

##### a. Genomic DNA preparation

Prepare genomic DNA starting from the 40 pooled bacterial pellets.

- Add 100 µl of 5 mg/ml lysozyme to the mycobacterial pellet of 2 ml and mix gently.
- Incubate O/N at 37°C (shaking).
- Shortspin and add pre-warmed 10% SDS to a final concentration of 2% (30 µl) and Proteïnase K to a final concentration of 33 µg/ml (2.5 µl). Adjust the volume to 150 µl with 20 mM Tris/HCl pH 9 (+17.5µl).
- Incubate for 3 h at 37°C (shaking).
- Add 30 µl of 5 M NaCl.
- Add 30 µl of CTAB/NaCl which is prewarmed at 65°C.
- Vortex and incubate 15 minutes at 65°C.
- Shortspin and add 200 µl of Phenol/Chloroform and mix gently by inverting the tube a few times.
- Centrifuge at room temperature for 15 minutes at 11,000 x g.
- Transfer aqueous supernatant to a new tube

- Add 200 µl of Phenol/Chloroform, mix gently by inverting the tube a few times.
- Centrifuge at room temperature for 10 minutes at 13,000 x g.
- Transfer aqueous supernatant to a new tube.
- Add 0.6 volumes of isopropanol to precipitate the nucleic acids.
- Place at least 30 minutes at -20°C (or longer, e.g. overnight).
- Centrifuge at room temperature for 10 minutes at 13,000 x g, discard the supernatant.
- Add 1 ml of cold 75% ethanol (-20°C) and invert the tube a few times.
- Centrifuge at room temperature for 10 minutes at 13,000 x g, remove supernatant carefully.
- Dry the pellet (10 minutes).
- Resuspend the pellet in 50 µl of 1x TE buffer, incubate 10-30 minutes until the DNA is completely dissolved.
- Keep at -4°C for short-term storage.
- Measure the DNA concentration spectrophotometrically (e.g. on a Nanodrop instrument) and fluorometrically (e.g. on a Qubit instrument with the Qubit dsDNA HS assay kit, 10 pg/µl - 100 ng/µl)
    - ! We need ~1 µg of DNA for the following steps.
    - ! Depending on the degree of saturation of the cultures, ± 2 µg of DNA can be obtained from a 2 ml culture.
    - ! An average $OD_{600}$ of 0.9 is insufficient to get 1 µg of DNA from 2 ml for all the pools. We suggest to grow the cultures to a higher OD (e.g. 1.5) or start from a bigger volume (e.g. 4 ml).

##### b. Genomic DNA fragmentation

- Transfer ± 1 µg of gDNA (20 ng/µl) into 1.5 ml TPX eppendorfs.
- Vortex samples 10-15 s, centrifuge 10 s, store samples 10-15’ on ice.
- Shear DNA with Bioruptor (setting High, 5 cycles, 30” on – 30” off, for size range of 200 – 1000 bp).
- Vortex samples 10-15 s, centrifuge 10 s.
- Control the fragmentation of the DNA by gel electrophoresis, e.g. by putting a 2 µl aliquot on the fragment analyzer (AATI) using the High Sensitivity NGS Fragment Analysis kit (DNF-474). Pools for which the fragmentation is not sufficient should have extra cycles until the size range falls between 200 – 1,000 bp.
    - ! Recommended to optimize the settings for your particular shearing device and samples

##### c. Blunt-end repair/dA-tailing of the fragmented DNA (NEBNext Ultra II End Repair/dA-tailing module)

- Reaction (assemble on ice)

| | |
|---|---|
| Fragmented gDNA sample | 48 µl |
| 8,5x end repair reaction buffer II | 7 µl |
| end prep enzyme mix | 3 µl |
| 1x TE buffer | 2 µl |
| Total | 60 µl |

- Mix well by pipetting, quick spin and run following program in a thermocycler

| | °C | time |
|---|---|---|
| 1x | 20°C | 30 min |
| 1x | 65°C | 30 min |
| 1x | 4°C | until further processing |

- DNA clean-up step with DNA binding magnetic beads. Elute in 20 µl of ultra-pure water.

- Measure concentration of the DNA on the Nanodrop and/or Qubit system.

  ! The original protocol used two different kits for performing the blunt-end repair and dA-tailing steps (1), by using a combined blunt-end repair/dA-tailing kit time is saved and an intermediated DNA clean-up step is avoided.

  ! If necessary samples can be stored at -20°C, however a slight loss in yield (20%) may be observed. It is recommended to continue with adaptor ligation before stopping.

#### d. Adaptor ligation

- Prepare the custom adaptors (**Table S8**) by mixing equimolar amounts of primer A000d (2.5 µl) and primer A00x (2.5 µl) with 45 µl duplex buffer (final concentration 10 µM) and running the following program in a thermocycler. These adaptors remain stable for at least several weeks when stored at 4°C.

| | °C | time |
|---|---|---|
| 1x | 95°C | 5 min |
| 1x | -0.1°C/s to 4°C | |
| 1x | 4°C | until further processing |

- Dilute dA-tailed fragmented DNA samples- to 6 ng/µl with ultra-pure water
- Ligase reaction using the T4 DNA ligase (Enzymatics) (assemble on ice). An equal result is achieved when using the T4 DNA ligase (NEB) following the manufacturer's instructions.

| | |
|---|---|
| 120 ng dA-tailed gDNA sample (6 ng/µl) | 20 µl |
| Pool-specific adaptor (10 µM) | 2 µl |
| MQ | 1.4 µl |
| 2x Rapid-ligation buffer (keep buffer on ice) | 25 µl |
| T4 DNA ligase (Enzymatics) | 1.6 µl (1000 U) |
| Total | 50 µl |

  ! Add excess of adaptor to the fragmented DNA (e.g. 20 – 100-fold molar excess, e.g. using 240 ng of 200 – 1,000 bp fragments (2 – 0.4 pmol) and 4 µl of a 10 µM adaptor mix (40 pmol)).

- Mix well by pipetting, quick spin
- Incubate at 25°C for 10 min when using the rapid T4 DNA ligase (Enzymatics)
- DNA clean-up step with DNA binding magnetic beads. Elute in 21 µl of ultra-pure water.

  ! A single clean-up step is insufficient to completely remove the excess of unligated adaptor, although the level of removal obtained by this means is sufficient for allowing the subsequent PCR reaction. Presuming > 95% recovery after DNA clean-up, we estimate the ligated DNA concentration at ± 10 ng/µl. For exact quantification at this step, consider multiple clean-up steps.

#### e. Tn gDNA-junction enrichment

- Prepare a mix of the P5-IR primers A, B, C, D2 in a 1:1:2:2 ratio (**Table S9**)
- Reaction

| | |
|---|---|
| 2x GoTaq Green mastermix | 25 µl |
| P7 primer (10 µM) | 2.5 µl |
| P5-IR primer mix (10 µM) | 2.5 µl |
| 20 µl template (~5-6 ng/µl) | 20 µl |
| Total | 50 µl |

- Mix well by pipetting, quick spin and run following program in a thermocycler

| | °C | time |
|---|---|---|
| 1x | 98°C | 3 min |
| | 98°C | 20 sec |
| 4x | 58°C | 20 sec |
| | 72°C | 3 min |
| | 98°C | 20 sec |
| 16x | 52°C | 20 sec |
| | 72°C | 3 min |
| 1x | 72°C | 10 min |
| 1x | 12°C | until further processing |

- Check the amplification by gel electrophoresis, e.g. by putting a 2 µl aliquot on the fragment analyzer (AATI) using the High Sensitivity NGS Fragment Analysis kit (DNF-474).

  ! Originally the protocol was performed with Phusion High Fidelity polymerase (NEB) (1), however this was unsuccessful for creating the library described in this study. The Phusion High Fidelity polymerase did not work to amplify the sample, in contrast amplification was achieved with KAPA Hifi hotstart polymerase, Q5 polymerase and GoTaq Green polymerase (Promega). We chose to use the GoTaq Green polymerase as this is the cheapest option.

  ! By using the High Sensitivity NGS Fragment Analysis kit (DNF-474) to analyze the amplification on the fragment analyzer, accurate concentration information can be acquired. This information can be used to ensure equal mixing of the DNA pools.

  ! First PCR cycles at higher annealing temperature to enrich the complementary strand using the P5-IR primer mix (normally no hybridization of P7 primer at this temperature). Then the annealing temperature is lowered to amplify the P5 – P7 fragments.

##### f. DNA pools mixing and size selection

- Combine the 40 PCR reactions for the 40 pools of one set of 96 96-well plates

  ! When mixing the PCR reactions, take into account that each A-H pool is derived from ± 768 mutants (96 x 8), while every 1-12 pool is derived from ± 1,152 (96 x 12) mutants. To equalize the sequencing read depth per mutant, mix 3 units of every row pool with 2 units of every column pool

  ! Using the concentration information from the fragment analyzer one can mix equal amounts of DNA instead of mixing equal volumes. We pooled 900 ng for each row pool and 600 ng for each column pool. This strategy improves the variation in sequencing coverage per pool. To further reduce this, one could consider to apply bead-based normalization (13) (included in some commercial kits) or enzyme-based normalization (e.g. Swift Biosciences Normalase kit).

- Run the mixture on a 2% preparative agarose gel
- Cut out the 250 – 750 bp region and extract the DNA from the agarose gel (e.g. Machery Nagel gel clean-up kit) and elute in 30 µl elution buffer
- Perform an extra DNA clean-up using DNA binding magnetic beads and elute in 0.1x TE buffer. This DNA clean-up step is added to make sure that the DNA is pure enough for subsequent quality control steps and finally Illumina sequencing.
- Confirm DNA fragment sizing on gel or the fragment analyzer using the High Sensitivity NGS Fragment Analysis kit (DNF-474)
- Measure exact DNA concentration (Nanodrop spectrophotometer, Qubit fluorometer, BioAnalyzer, Fragment analyzer, e.g. using the High Sensitivity NGS Fragment Analysis kit (AATI, DNF-474) or qPCR-based quantification, e.g. using KAPA library quantification kit (Roche, #KK4610))
- Sequence the sample on a NextSeq (single index sequencing, medium throughput, SE150 bp, Illumina) using the standard Illumina sequencing primers.

- ! Addition of 45% PhiX to the samples was necessary to compensate for the remaining low complexity in the first sequencing bases. Adding 10% PhiX resulted in ~50 M barcoded reads, while adding 45% PhiX resulted in ~70 M barcoded reads.
- ! In the original protocol of Vandewalle *et al.* 2015 the samples were run on an Illumina HiSeq 2,500 chip (SE100 bp) (1).
- ! Aim to get ~70 M single-end sequencing reads to allow successful characterization of the ordered library.

#### 2.4. Sequencing data analysis and deconvolution of the mutant positions

The raw Illumina sequencing data was processed using the Galaxy web platform (http://galaxyproject.org/) only (14). The original data analysis protocol of Vandewalle *et al.* 2015 (1) combined the use of the Galaxy platform with the use of CLC Genomics Workbench (http://www.clcbio.com/products/clc-genomics-workbench/). The main analysis outline using the free Galaxy software is given below, the Galaxy tool to fulfill each operation is given in between brackets. The optimized Galaxy workflows and the BioPerl CP-CSeq algorithm can be found in **Section 3** and **Section 4**, the .ga and .pl files can be accessed from the Figshare repository with DOI https://doi.org/10.6084/m9.figshare.c.4507472.

The data analysis protocol serves for Illumina sequencing data for one set of 96 96-well plates. When a library consists of multiple sets of 96 96-well plates, the data analysis must be performed separately for these different sets. Afterwards, the TnLists generated for each separate set of 96 96-well plates can be joined together to create a final list.

##### a. Adaptor trimming and select transposon-specific reads

After Illumina sequencing, the Illumina software automatically demultiplexes the barcodes and generates 40 FastQ files, each one representing the raw sequencing reads of a given Cartesian pool. NextSeq has however 4 physical lanes and produces 4 FastQ files per adaptor instead of 1. Therefore, these 4 FastQ files from NextSeq need to be concatenated to 1 FastQ file per barcode, if the sequencing service facility has not done this already.

Import (**Galaxy tool: Upload file**) the FastQ files on the Galaxy server. In addition, import the genome sequence and annotation files (e.g. *M. bovis* BCG Danish 1331 (NIBSC 07/270, can be accessed from the Figshare repository with DOI https://doi.org/10.6084/m9.figshare.c.4489496 (15)). The quality of the sequencing data can be analyzed with **FastQC** and **MultiQC** in Galaxy.

From these sequences trim the adaptor (3') and primer (5') sequences, select transposon-specific reads and trim the remaining reads (**Galaxy workflow 1: Quality filtering-trimming-clipping transposon sequences**). The workflow selects only those reads containing the 8 bp transposon-specific tag (CAACCTGT) and discards the rest. It trims the remaining reads to a given length, set standard to 25bp. This length of 25 bp is sufficient to confidently map reads to a bacterial genome and not too long to avoid overlap when TA's are close to each other. The trimming step facilitates the calculation of the coverage of each mutant (at their inserted TA in the genome) at later steps.

Optionally, one can analyze the quality of the trimmed sequencing data with **FastQC** and **MultiQC** in Galaxy, to validate that the quality of the sequencing data has improved and the adaptor and primer sequences have been removed.

- ! Longer sequencing fragments, e.g. SE150 bp versus SE100 bp, improve the data analysis as a higher percentage of sequences are retained after trimming leading to a higher percentage of aligned reads and thus more transposon insertions that can be assigned a position.

##### b. Read mapping to the reference genome

Map the reads to the organism's reference genome (**Galaxy workflow 2: Mapping**), which generates a SAM/BAM output for each Cartesian pool (= alignment file with mapping information for each read).

The original protocol by Vandewalle *et al.* 2015 (1) instructs to discard reads that do not uniquely map to the reference genome, as this avoids the uncertain mapping of mutants in duplicated regions of the genome. However, this strategy also discards the transposon insertions for which one of the two reads maps uniquely and the other one not, which are also transposon insertions mapping to a unique region. Adhering to this strategy leads to the exclusion of ~100 transposon insertions targeting such one-sided unique regions.

To include both two-sided and one-sided unique regions in the transposon list, one needs to follow the following strategy. Make a TnList_UNIQUES containing only transposon insertions targeting two-sided unique regions by discarding reads that do not uniquely map to the reference genome (**Galaxy workflow 2a: Mapping uniques_mismatch 1 or 0**) for further data analysis. Make a TnList_ALL containing all transposon insertions targeting one-sided and two-sided unique regions and duplicated regions by including all reads that map in the data analysis (**Galaxy workflow 2b: Mapping all_mismatch 1 or 0**) for further data analysis. Using both lists, one can then extract TnList_DUPLICATED containing all transposon insertions targeting one-sided unique regions and duplicated regions (**Section 2.4.f**). In this TnList coverage profiles that occur once are from transposon insertions targeting one-sided unique regions and coverage profiles that occur more than once are from transposon insertions targeting duplicated regions. To identify the distinct transposon insertions targeting duplicated regions, the duplicated coverage profiles are discerned (TnList_DUPLICATED_condensed). Lastly, the TnList_UNIQUE and TnList_DUPLICATED condensed are merged to create a final TnList.

- ! This strategy seems to double the data-analysis, however identifying the transposon insertions targeting duplicated regions directly on TnList_ALL is discouraged as this wrongly identifies transposon insertions that have an identical coverage profile due to chance as transposon insertions targeting duplicated regions.
- ! For sequencing reads of high quality, the changes are minimal if 0 or 1 mismatches are allowed. For sequencing reads of low quality, the output differs more, depending on allowing 0 or 1 mismatches. Therefore, one needs to evaluate which settings to use, while remembering 'rubbish in, means rubbish out'. Allowing 0 mismatches, means that you are 100% sure about the resulting data. However, for a parental strain for which no close reference genome exists, allowing 1 mismatch could be the best option, to also find the transposon insertions at sites for which the parental strain has a SNP compared to the reference genome.
- ! Terms explained
  - o two-sided unique regions: Fw and Rv reads, each 20 bp, map uniquely to the genome
  - o one-sided unique regions: Fw or Rv reads, not both, map uniquely to the genome
  - o duplicated regions: Fw and Rv reads don't map uniquely to the genome

##### c. Generate the transposon insertion mutant list

Next, we want to generate a list of every transposon-insertion mutant present in the library. First, we need to merge the 40 BAM files of all Cartesian coordinate groups together to one BAM file (**Galaxy tool: Merge SAM files**). From this BAM file, we generate a file containing the coverage for reads that map to the top strand of the genome (**Galaxy workflow 3a: Calculate coverage around TA positions with aligned reads – ForwardReads**), and another one for reads that map to the bottom strand (**Galaxy workflow 3b: Calculate coverage around TA positions with aligned reads – ReverseReads**).

Subsequently, these independent files are joined to create a complete transposon insertion list of the entire library, discarding sites where only forward or reverse reads start (**Galaxy workflow 4a: FOR MERGED FILE_CreateTnList_Forward and Reverse Reads**). Every dinucleotide site where the 5' end of the forward and reverse reads overlap at that dinucleotide site, with a total read coverage >60, is considered a transposon insertion. Generate an interval file (TnList) containing the position of each TA in the genome with a transposon insertion, where each transposon insertion gets a different number.

- ! The Himar1 position preferably integrates at TA dinucleotides (~99,7%), however also a small percentage of transposon insertions at other sites are observed. So when TA dinucleotides are mentioned in the protocol, we broadly mean all transposon insertion sites, including the non-TA dinucleotides.
- ! The visual representation of the merged BAM file can already reveal the location of the distinct transposon insertion mutants in the library. One can also identify the transposon insertions in highly repetitive and duplicated regions of the genome, by visualizing the merged BAM file not containing reads that don't uniquely map to the genome versus the merged BAM file containing those reads.
- ! Overlapping Fw and Rv reads with a total read coverage <60 are filtered out, as the majority are false positives due to aspecific amplification.

Using **Galaxy workflow 5: Add annotation to transposon list**, additional information can be added to this transposon insertion mutant interval file e.g. gene loci and gene names, to ease interpretation of the list during data analysis. This requires an annotation file (in interval format), with columns mentioning chromosome, start, end, gene locus and gene name.

#### d. Calculate coverage file for each Cartesian pool

To calculate the coverage file for each Cartesian pool, run the above workflows (**Galaxy Workflow 3a and 3b**) for each Cartesian pool SAM file. Generate two independent files containing the coverage at each TA dincleotide with reads that map to the top or bottom strand for each pool. Join these independent files to create an interval file for each pool, containing coverage information of each mutant in that pool (**Galaxy workflow 4b: FOR SEPARATE FILES_CreateTnList_Forward and Reverse Reads**).

Next, paste these coverage files together per Cartesian pool group, so pool A till H **(Galaxy Workflow 6a: Join 8 Datasets TAcoverage JoinEnd Start)** and pool 1 till 12 **(Galaxy Workflow 6a: Join 8 Datasets TAcoverage JoinEnd Start)** for both X-Y and Z Pool Plates). This creates 4 interval files in tab-delimited format where each row represents one mutant and where the Cartesian pool coverage is listed per column. Combine both X-Y interval files (1-12 and A-H) together and similarly combine both Z interval files (1-12 and A-H) together **(Galaxy Workflow 7: Join 1-12 and A-H coverage files)**. This creates two tab-delimited files with first 7 columns (column c1 to c7) mutant and gene information, and then 20 columns containing the reads counts of each mutant per Cartesian pool (pool 1-12 in c9 to c20, pool A-H in c22 to c29, c8 and c21 are empty). Save files respectively as 'TnList_XY.txt' and 'TnList_Z.txt'.

Manipulate this tab-delimited format further in e.g. MS Excel. Normalize each read count in these files to the average number of reads per Cartesian pool group. Therefore, first calculate the sum of the coverage counts for each pool. Then calculate the average sum of the coverage count for each Cartesian pool group. Finally, calculate the normalized read counts for each well for each Cartesian pool, by using the mathematical rule of three. E.g. if the total coverage count in Cartesian pool X-Y column 1 is 2.2 million, and the average coverage count across column pool 1 to 12 is 2 million, divide every read count in X-Y column 1 by a factor of 1.1 (2.2 million over 2 million). Save files respectively as 'TnList_XY_normalized.txt' and 'TnList_Z_normalized.txt'. This normalization is an essential step before proceeding with the deconvolution of the mutant coordinates.

Example formulas for the normalization in Excel:

- TnList with transposon insertions in row 2 to 8000. Coverage counts for pool 1-12 in c9 to c20 (I-T) and for pool A-H in c22 to c29 (V-AC)
- Calculate the sum of the coverage counts for each pool in column 9-20 (I-T) and column 22-29 (V-AC): =SUM(coverage transposon insertion 1: coverage transposon insertion n), e.g. =SUM(I2:I800)
- Calculate the average sum of the coverage count for each Cartesian pool group; =ROUND(AVERAGE(pool 1: pool n);0), e.g. =ROUND(AVERAGE(I1:T1);0)
- Calculate the normalized read counts for each well for each Cartesian pool in column (AE-AP) and column (AR-AY), well=IF(well=".";".";ROUND(well/(sum/average sum);0)), e.g. AE2=IF(I2=".";".";ROUND(I2/(I$1/$U$1);0)
- Make a copy of the current tab, select all and past again as numbers.
- Delete the original read counts (column 9-21 (I-AD)) and the first calculation row. Generating an identical list as originally, but with normalized read counts.

  ! Better to perform the normalization without an extreme outlier in the average. However, it is best to investigate for each case, which option gives a better result (leaving the outlier in or out from the average sum for normalization) when applying the CP-CSeq algorithm.

#### e. CP-CSeq algorithm

Run the CP-CSeq algorithm (**CP-CSeq_BioPerlScript**) to deconvolute the mutant coordinates for each transposon insertion for which there are sufficient data for that transposon insertion in the dataset. The algorithm requires the two tabular (.txt) input files ('TnList_XY_normalized.txt' and 'TnList_Z_normalized.txt'). Put the .txt files in the same folder as the CP-CSeq_BioPerlScript.pl file and execute the algorithm, which will result in the generation of the output files in that same folder. The complete BioPerl CP-CSeq algorithm, with explanatory comments, can be found in **Section 4**.

A variable "librarySet" allows the user to specify for which set of 96 96-well plates the output is generated, e.g. set I, set II, set III, … .

Output files from the CP-CSeq algorithm:

- "Summary.txt". The file contains the summary statistics.
- "TnListOrderedByPlate.txt" contains the assigned position(s) for each transposon insertion ordered by plate. The file lists if the wells are filled with one unique mutant (UNIQUE) or if the wells contain more than one mutant (MZS with 2,3, …) (according to the protocol). The file contains 5 columns: (c1) transposon insertion-number, (c2) locus, (c3) XY coordinate, (c4) Z coordinate, (c5) information about the number of mutants assigned to a well.
- "TnListOrderedByMutant.text" contains the assigned position(s) for each transposon insertion ordered by mutant. The file lists if the automatic deduction of position of the mutant was unique, Zok_unique, Zok_nonunique, XYok_unique, XYok_nonunique or heuristically determined. The file contains 5-n columns: (c1) information about how the position was deducted, (c2) transposon insertion-number, (c3) locus, (c4) XY coordinate, (c5) Z coordinate, (c6-cn) additional XY or Z coordinates
- "FinalTnList.txt" contains the coordinates for each mutant (= row) in the list, provided that sufficient data for that mutant is present in the dataset. This file is a cleaned and reordered version of the file "TnListOrderedByMutant.text". The file contains 4-n columns: (c1) transposon insertion-number, (c2) locus, (c3) information about how the position was deducted, (c4) position, (c5-cn) alternative positions for the mutant. The position information is built as followed: Set-x1.Plate-x2.Well-x3 (e.g. Set-I.Plate-23.Well-A3), with x1 the Roman number for the library set, x2 the plate number, x3 the well coordinates.

#### f. Finalizing the TnLists and making them user-friendly

The output of the CP-CSeq algorithm doesn't retain the information of the genes that the transposon insertions target. Run **Galaxy Workflow 8: Add chr/start/end/name and coverages to FinalTnList**, to add this gene information and the coverages to the TnList. It is recommended to add the complete coverage profile for the 40 pools to the TnList, as this allows to cross-check the CP-CSeq algorithm's output for each transposon insertion.

Likewise, it is recommended to calculate the total TA coverage for the pools, by taking the sum of the TA coverage for each pool. Subsequently, color coding can be added (e.g. via conditional formatting in Excel) to allow easy evaluation of the position assignment for the transposon insertion, i.e. whether the position assignment is well-supported by sequencing data. Possible color coding: <300 (white) low coverage, 300-1200 (green) medium coverage and >1200 (dark green) high coverage (dark green).

As mentioned in **Section 2.4.b**, the full data-analysis protocol should be run twice to generate TnList_UNIQUE (containing only transposon insertions targeting two-sided unique regions by discarding reads that do not uniquely map to the reference genome) and TnList_ALL (containing all transposon insertions targeting one-sided and two-sided unique regions and duplicated regions by including all reads that map). **Galaxy workflow 9: Extracting TnList_DUPLICATED** can be used to identify the transposon insertions targeting one-sided unique and duplicated regions. The workflow achieves this by comparing the TnList_ALL and TnList_UNIQUES and extracting the distinct lines. In addition, it filters out transposon insertions with a total TA coverage <60 after which the file is sorted based on the coverage columns, to allow the identification of unique lines based on the unique coverage profile. Two output files are created: one in which only the unique lines are kept, but the number of occurrences of each line is printed (TnList_DUPLICATED_condensed) and one in which the duplicated lines are grouped together (TnList_DUPLICATED_grouped). Subsequently, the TnList_UNIQUE and TnList_DUPLICATED_condensed can be merged to create a final TnList, including all transposon insertions (targeting one-sided and two-sided unique regions and duplicated regions) appearing once in the list. After this step one needs to sort the TnList on transposon insertion_Start and assign a new transposon insertion number. If the complete library consisted out of different sets of 96 96-well plates, then also specify the specific set of 96 96-well plates in the transposon insertion number, e.g. transposon insertion-Setx-x. Then, in a final step, one can merge the TnLists for these different sets into one TnList, while retaining the information in which set each transposon insertion was found.

Additional interesting information to add to the TnList, includes:

- Duplication status of the region in which the transposon insertion occurs: one-sided unique region, two-sided unique region, duplicated region
- Dinucleotide in which the transposon was inserted
- Additional gene information: strand (+/-), gene length, gene product information, number of TA per gene

- *M. bovis* BCG Pasteur and *M. tb* orthologue gene loci.
- Tn_% after start codon, which indicates (in %) how far after the start codon the transposon is inserted. Which is calculated by the Excel formula: =IF(Strand=".";".";IF(Strand="+";100*(Tn_Start-Gene_Start)/Length_Gene; 100-100*(Tn_Start-Gene_Start)/Length_Gene))
- Gene essentiality of their *M. tb* orthologs derived from publications (16–19)
- The TnList_DUPLICATED_grouped can be added to the Excel file in a separate tab to allow the user to find the other regions a transposon insertion targeting a duplicated region targets. As a transposon insertion targeting a duplicated region is only included once in the final TnList.

Information from an interval file can be added to the TnList in Galaxy (**Galaxy tool: Join the intervals of two datasets side-by-side**, with min overlap 1). Be aware that rows with transposon insertions targeting overlapping genes get duplicated in this manner. If needed these can be removed in Galaxy (**Galaxy tool: Sort data in ascending or descending order**, Version 1.1.0, sort on transposon insertion column, ascending order, natural sort, only keep unique lines as output) or in Excel (remove duplicates based on transposon insertion column).

Recommendations for choosing mutants:

1) look at the Tn_% after Start codon, which indicates (in %) how far after the start codon the transposon is inserted (1% = near the start codon; 100% = near the stop codon), at …% of the ORF. To have a knock-out mutant you would want the transposon inserted not far after the start codon or in front of the start-codon in the promoter region.

2) look at the location assignment, you would want mutants that were positioned well in the library. The location assignment 'unique' is recommended, then 'Z/XYok_unique', 'Z/XYok_nonunique' and finally 'heuristic'.

3) look at the total TA coverage, which gives you the total coverage over all the pools for each mutant. Mutants with high coverage are preferred as their position assignment is better supported by data. TA coverage <300 (white) low coverage, 300-1200 (green) medium coverage and >1200 (dark green) high coverage (dark green).

4) look at the TA coverage for each pool. To double check the location assignment for the mutant. An outlier pool can skew the data and influence the assignments.

Some statistics can be extracted from the output file Summary.txt of the CP-CSeq_BioPerlScript. A different range of statistics can be calculated on the final TnLists e.g. using pivot tables in Excel. For some calculations the composed columns need to be uncomposed (e.g. position).

### 3. Galaxy workflows

| **Galaxy workflow 1: Quality filtering-trimming-clipping tn sequences** | |
|---|---|
| **Step** | **Annotation** |
| Step 1: Input dataset<br>FastQ files: *select at runtime* -> e.g. trimmed_KB_I_A001 to A0040 | one FastQ file for each barcode |
| Step 2: Trim Galore!<br>**Is this library paired- or single-end?** Single-end<br>**Reads in FASTQ format:** Output dataset 'output' from step 1<br>**Adapter sequence to be trimmed:** User defined adapter sequence<br>**Adapter sequence to be trimmed off:** AGATCGGAAGAGCACACGTCTGAACT<br>**Remove N bp from the 3' end**: Not available.<br>**Trim Galore! advanced settings:** Full parameter list<br>**Trim low-quality ends from reads in addition to adapter removal (Enter phred quality score threshold):** 20<br>**Overlap with adapter sequence required to trim a sequence:** 1<br>**Maximum allowed error rate**: 0.1<br>**Discard reads that became shorter than length N:** 15<br>**Instructs Trim Galore! to remove N bp from the 5' end of read 1:** Not available.<br>**Instructs Trim Galore! to remove N bp from the 5' end of read 2 (Only for paired-end reads):** Not available.<br>**Generate a report file:** False<br>**specify if you would like to retain unpaired reads:** Do not output unpaired reads<br>**RRBS specific settings:** Use defaults (no RRBS) | Trim adapter sequences using Trim Galore! |
| Step 3: Reverse-Complement<br>**Library to reverse-complement:** Output dataset 'trimmed_reads_single' from step 2 | |
| Step 4: Trim Galore!<br>**Is this library paired- or single-end?** Single-end<br>**Reads in FASTQ format:** Output dataset 'output' from step 3<br>**Adapter sequence to be trimmed:** User defined adapter sequence<br>**Adapter sequence to be trimmed off:** GCTGATAAGTCCCCGGTCTC<br>**Remove N bp from the 3' end:** Not available.<br>**Trim Galore! advanced settings:** Full parameter list<br>**Trim low-quality ends from reads in addition to adapter removal (Enter phred quality score threshold):** 20<br>**Overlap with adapter sequence required to trim a sequence:** 1<br>**Maximum allowed error rate:** 0.1<br>**Discard reads that became shorter than length N:** 15<br>**Instructs Trim Galore! to remove N bp from the 5' end of read 1:** Not available.<br>**Instructs Trim Galore! to remove N bp from the 5' end of read 2 (Only for paired-end reads):** Not available.<br>**Generate a report file:** False<br>**specify if you would like to retain unpaired reads:** Do not output unpaired reads<br>**RRBS specific settings:** Use defaults (no RRBS) | Trim primer sequences (TL028) using Trim Galore! |
| Step 5: Clip<br>**Library to clip:** Output dataset 'trimmed_reads_single' from step 4<br>**Minimum sequence length (after clipping, sequences shorter than this length will be discarded):** 25<br>**Source:** Enter custom sequence<br>**Enter custom clipping sequence :** ACAGGTTG<br>**enter non-zero value to keep the adapter sequence and x bases that follow it:** 0<br>**Discard sequences with unknown (N) bases:** Yes<br>**Output options:** Output only clipped sequences (i.e. sequences which contained the adapter) | Select reads with contain the 8 bp tn-specific tag and discards the rest by using the tool clip adapter sequences (cuts from 3') |
| Step 6: Reverse-Complement<br>**Library to reverse-complement:** Output dataset 'output' from step 5 | |
| Step 7: Trimmomatic<br>**Single-end or paired-end reads?:** Single-end<br>**Input FASTQ file:** Output dataset 'output' from step 6<br>**Perform initial ILLUMINACLIP step?:** False<br>**Select Trimmomatic operation to perform:** Cut the read to a specified length (CROP)<br>**Number of bases to keep from the start of the read:** 25 | Trim remaining reads to 25 bp. With the crop operation from the trimmomatic tool |

| |
|---|
| Final output: **trimmed_input name** (e.g. trimmed_KB_I_A001 to A0040), all the rest of the outputs are hidden and carry the input name |
| **Output name**: Trimmed_x |

| **Galaxy workflow 2a: Mapping uniques_mismatch 1 or 0** | |
|---|---|
| **Step** | **Annotation** |
| Step 1: Input dataset<br>**input:** *select at runtime* | Select the reference genome (fasta file) |
| Step 2: Input dataset<br>**input**: *select at runtime* | Select the trimmed FastQ files |
| Step 3: Map with Bowtie for Illumina<br>**Will you select a reference genome from your history or use a built-in index?** Use one from the history<br>**Select the reference genome:** Output dataset 'output' from step 1<br>**Choose whether to use Default options for building indices or to Set your own:** Default<br>**Is this library mate-paired?** Single-end<br>**FASTQ file:** Output dataset 'output' from step 2<br>**Bowtie settings to use:** Full parameter list<br>**Skip the first n reads (-s)**: 0<br>**Only align the first n reads (-u)**: -1<br>**Trim n bases from high-quality (left) end of each read before alignment (-5):** 0<br>**Trim n bases from low-quality (right) end of each read before alignment (-3):** 0<br>**Alignment mode:** Maq-like: quality-aware, limit mismatches in seed (-n)<br>**Maximum number of mismatches permitted in the seed (-n):** 1 or 0<br>**Maximum permitted total of quality values at all mismatched read positions (-e):** 70<br>**Seed length (-l):** 28<br>**Whether or not to round to the nearest 10 and saturating at 30 (--nomaqround):** Round to nearest 10<br>**Choose whether or not to attempt to align against the forward reference strand (--nofw):** Align against the forward reference strand<br>**Choose whether or not to attempt to align against the reverse-complement reference strand (--norc):** Align against the reverse-complement reference strand<br>**Whether or not to make Bowtie guarantee that reported singleton alignments are 'best' in terms of stratum and in terms of the quality values at the mismatched positions (--best):** Do not use best<br>**Whether or not to try as hard as possible to find valid alignments when they exist (-y):** Do not try hard<br>**Maximum number of backtracks permitted when aligning a read (--maxbts):** 125<br>**Whether or not to report all valid alignments per read (-a):** Report all valid alignments<br>**Suppress all alignments for a read if more than n reportable alignments exist (-m):** 1<br>**Write all reads with a number of valid alignments exceeding the limit set with the -m option to a file (--max):** False<br>**Write all reads that could not be aligned to a file (--un):** False<br>**Override the offrate of the index to n (-o):** -1<br>**Seed for pseudo-random number generator (--seed):** -1<br>**Save the bowtie mapping statistics to the history:** True<br>**Suppress the header in the output SAM file (--sam-nohead):** False | |
| Step 4: SortSam<br>**Select SAM/BAM dataset or dataset collection:** Output dataset 'output' from step 3<br>**Sort order:** Coordinate<br>**Select validation stringency:** Lenient | |
| **Output name**: mapped_unique_1 or 0 MM | |

| **Galaxy workflow 2b: Mapping all_mismatch 1 or 0** | |
|---|---|
| **Step** | **Annotation** |
| Step 1: Input dataset<br>**input:** *select at runtime* | Select the reference genome (fasta file) |
| Step 2: Input dataset<br>**input**: *select at runtime* | Select the trimmed FastQ files |
| Step 3: Map with Bowtie for Illumina<br>**Will you select a reference genome from your history or use a built-in index?** Use one from the history<br>**Select the reference genome:** Output dataset 'output' from step 1 | |

| |
|---|
| **Choose whether to use Default options for building indices or to Set your own:** Default<br>**Is this library mate-paired?** Single-end<br>**FASTQ file:** Output dataset 'output' from step 2<br>**Bowtie settings to use:** Full parameter list<br>**Skip the first n reads (-s)**: 0<br>**Only align the first n reads (-u)**: -1<br>**Trim n bases from high-quality (left) end of each read before alignment (-5):** 0<br>**Trim n bases from low-quality (right) end of each read before alignment (-3):** 0<br>**Alignment mode:** Maq-like: quality-aware, limit mismatches in seed (-n)<br>**Maximum number of mismatches permitted in the seed (-n):** 1 or 0<br>**Maximum permitted total of quality values at all mismatched read positions (-e):** 70<br>**Seed length (-l):** 28<br>**Whether or not to round to the nearest 10 and saturating at 30 (--nomaqround):** Round to nearest 10<br>**Choose whether or not to attempt to align against the forward reference strand (--nofw):** Align against the forward reference strand<br>**Choose whether or not to attempt to align against the reverse-complement reference strand (--norc):** Align against the reverse-complement reference strand<br>**Whether or not to make Bowtie guarantee that reported singleton alignments are 'best' in terms of stratum and in terms of the quality values at the mismatched positions (--best):** Do not use best<br>**Whether or not to try as hard as possible to find valid alignments when they exist (-y):** Do not try hard<br>**Maximum number of backtracks permitted when aligning a read (--maxbts):** 125<br>**Whether or not to report all valid alignments per read (-a):** Report all valid alignments<br>**Suppress all alignments for a read if more than n reportable alignments exist (-m):** -1<br>**Write all reads with a number of valid alignments exceeding the limit set with the -m option to a file (--max):** False<br>**Write all reads that could not be aligned to a file (--un):** False<br>**Override the offrate of the index to n (-o):** -1<br>**Seed for pseudo-random number generator (--seed):** -1<br>**Save the bowtie mapping statistics to the history:** True<br>**Suppress the header in the output SAM file (--sam-nohead):** False |
| Step 4: SortSam<br>**Select SAM/BAM dataset or dataset collection:** Output dataset 'output' from step 3<br>**Sort order:** Coordinate<br>**Select validation stringency:** Lenient |
| **Output name**: mapped_all_1 or 0 MM |

| **Galaxy workflow 3a: Calculate coverage around TA positions with aligned reads - ForwardReads** | |
|---|---|
| **Step** | **Annotation** |
| Step 1: Input dataset<br>**input:** *select at runtime* | SAM file with alignments |
| Step 2: BAM-to-SAM<br>**BAM File to Convert:** Output dataset 'output' from step 1<br>**Header options:** Include header in SAM output (-h) | |
| Step 3: Cut<br>**Cut columns:** c2,c4,c6<br>**Delimited by:** Tab<br>**From:** Output dataset 'output' from step 2 | Remove superfluous columns. Keep columns: FLAG (reads mapped to sense or antisense strand - 0 or 16), POS (Startposition of read in genome) and CIGAR string (contains e.g. read length info) |
| Step 4: Trim<br>**Input dataset:** Output dataset 'out_file1' from step 3<br>**Trim this column only:** 3<br>**Trim from the beginning up to this position:** 1<br>**Remove everything from this position to the end:** -1<br>**Is input dataset in FASTQ format?:** No<br>**Ignore lines beginning with these characters:** Nothing selected. | Remove last character of CIGAR string (-M; remaining characters = read length) |
| Step 5: Filter<br>**Filter:** Output dataset 'out_file1' from step4<br>**With following condition:** c3==25<br>**Number of header lines to skip:** 0 | Remove rows containing a 'read length' (c3) that is not 25. This step is necessary if non-standard CIGAR strings were present (standard = 25M); otherwise subsequent tools may generate an error. **Be aware some mapping tools generate a 'string' instead of 'number' for this column, then you need to change c3==25 by** |

| | |
|---|---|
| | **c3==’25’.** |
| Step 6: Filter<br>**Filter:** Output dataset 'out_file1' from step 5<br>**With following condition:** c1==16<br>**Number of header lines to skip:** 0 | Only retains rows with Flag = 16; this keep reads mapped to (-) strand (end with 'TA') |
| Step 7: Compute<br>**Add expression:** (c2+c3)-2<br>**as a new column to:** Output dataset 'out_file1' from step 6<br>**Round result?:** YES | Compute on reads mapped to (-) strand only: (Position in genome + Read length) - 2 ('TA') = position of 'T' of 'TA' dinucleotide |
| Step 8: Group<br>**Select data:** Output dataset 'out_file1' from step 7<br>**Group by column:** 4<br>**Ignore case while grouping?:** False<br>**Ignore lines beginning with these characters:** Nothing selected.<br>**Operations**<br>**Operation 1**<br>**Type:** Count<br>**On column:** 1<br>**Round result to nearest integer?** NO | (-) reads. Group by position of 'TA' in genome and count prevalence of these positions => this equals the coverage at TA positions in the genome, but only when the read starts at the TA |
| Step 9: Sort<br>**Sort Dataset:** Output dataset 'out_file1' from step 8<br>**on column:** 1<br>**with flavor:** Numerical sort<br>**everything in:** Ascending order<br>**Column selections**<br>**Number of header lines to skip**: 0 | |
| Step 10: Add column<br>**Add this value:** Chromosome<br>**to Dataset:** Output dataset 'out_file1' from step 9<br>**Iterate?** NO | |
| Step 11: Compute<br>**Add expression:** c1+1<br>**as a new column to:** Output dataset 'out_file1' from step 10<br>**Round result?** YES | |
| Step 12: Cut<br>**Cut columns:** c3,c1,c4,c2<br>**Delimited by:** Tab<br>**From:** Output dataset 'out_file1' from step 11 | |
| **Output name**: TA_ForwardReads_x | |

| **Galaxy workflow 3b: Calculate coverage around TA positions with aligned reads - ReverseReads** | |
|---|---|
| **Step** | **Annotation** |
| Step 1: Input dataset<br>**input:** *select at runtime* | SAM file with alignments |
| Step 2: BAM-to-SAM<br>**BAM File to Convert:** Output dataset 'output' from step 1<br>**Header options:** Include header in SAM output (-h) | |
| Step 3: Cut<br>**Cut columns:** c2,c4,c6<br>**Delimited by:** Tab<br>**From:** Output dataset 'output' from step 2 | Remove superfluous columns. Keep columns: FLAG (reads mapped to sense or antisense strand - 0 or 16), POS (Startposition of read in genome) and CIGAR string (contains e.g. read length info) |
| Step 4: Trim<br>**Input dataset**<br>Output dataset 'out_file1' from step 3<br>**Trim this column only:** 3<br>**Trim from the beginning up to this position:** 1<br>**Remove everything from this position to the end:** -1<br>**Is input dataset in FASTQ format?:** No | Remove last character of CIGAR string (-M; remaining characters = read length) |

| | |
|---|---|
| **Ignore lines beginning with these characters:** Nothing selected. | |
| Step 5: Filter<br>**Filter:** Output dataset 'out_file1' from step 4<br>**With following condition:** c3==25<br>**Number of header lines to skip:** 0 | Remove rows containing a 'read length' (c3) that is not 25. This step is necessary if non-standard CIGAR strings were present (standard = 25M); otherwise subsequent tools may generate an error. **Be aware some mapping tools generate a 'string' instead of 'number' for this column, then you need to change c3==25 by c3=='25'.** |
| Step 6: Filter<br>**Filter:** Output dataset 'out_file1' from step 5<br>**With following condition:** c1==0<br>**Number of header lines to skip:** 0 | Only retains rows with Flag = 0; this keep reads mapped to (+) strand (start with 'TA') |
| Step 7: Group<br>**Select data:** Output dataset 'out_file1' from step 5<br>**Group by column:** 2<br>**Ignore case while grouping?:** False<br>**Ignore lines beginning with these characters:** Nothing selected.<br>**Operations**<br>**Operation 1**<br>**Type:** Count<br>**On column:** 1<br>**Round result to nearest integer?**NO | (+) reads. Group by position of 'TA' in genome and count prevalence of these positions => this equals the coverage at TA positions in the genome, but only when the read starts at the TA |
| Step 8: Sort<br>**Sort Dataset:** Output dataset 'out_file1' from step 7<br>**on column:** 1<br>**with flavor:** Numerical sort<br>**everything in:** Ascending order<br>**Column selections**<br>**Number of header lines to skip**: 0 | |
| Step 9: Add column<br>**Add this value:** Chromosome<br>**to Dataset:** Output dataset 'out_file1' from step 9<br>**Iterate?**NO | |
| Step 10: Compute<br>**Add expression:** c1+1<br>**as a new column to:** Output dataset 'out_file1' from step 9<br>**Round result?**YES | |
| Step 11: Cut<br>**Cut columns:** c3,c1,c4,c2<br>**Delimited by:** Tab<br>**From:** Output dataset 'out_file1' from step 10 | |
| **Output name**: TA_ReverseReads_x | |

| **Galaxy workflow 4a: FOR MERGED FILE_CreateTnList_Forward and Reverse Reads** | |
|---|---|
| **Step** | **Annotation** |
| Step 1: Input dataset<br>**input:** *select at runtime* | Input list of possible tn insertions forward reads - Chromosome (c1), start (c2), stop (c3), coverage on TA (c4) |
| Step 2: Input dataset<br>**input:** *select at runtime* | Input list of possible tn insertions reverse reads - Chromosome (c1), start (c2), stop (c3), coverage on TA (c4) |
| Step 3: Join two Datasets<br>**Join:** Output dataset 'output' from step 1<br>**using column:** 2<br>**with:** Output dataset 'output' from step 2<br>**and column:** 2<br>**Keep lines of first input that do not join with second input:** No<br>**Keep lines of first input that are incomplete:** No<br>**Fill empty columns:** No<br>**Keep the header lines:** No | |
| Step 4: Compute | |

| | |
|---|---|
| **Add expression:** c4+c8<br>**as a new column to:** Output dataset 'out_file1' from step 3<br>**Round result?**YES | |
| Step 5: Filter<br>**Filter:** Output dataset 'out_file1' from step 4<br>**With following condition:** c9>=60<br>**Number of header lines to skip:** 0 | Filter out overlapping Fw and Rv reads with a total read coverage <60. Most of them are false positives due to aspecific amplification, which we want to limit in our final library. |
| Step 6: Add column<br>**Add this value:** TnInsertion<br>**to Dataset:** Output dataset 'out_file1' from step 4<br>**Iterate?**YES | |
| Step 7: Cut<br>**Cut columns:** c1,c2,c3,c10,c9<br>**Delimited by:** Tab<br>**From:** Output dataset 'out_file1' from step 5 | Output = interval file containing chromosome (c1), Tn start (c2), Tn stop (c3), TnInsertion number (c4), total coverage on TA (Fw and Rev reads starting on TA only) (c5) |
| **Output name**: TnList_TotalTAcoverage_x | |

| **Galaxy workflow 4b: FOR SEPARATE FILES_CreateTnList_Forward and Reverse Reads** | |
|---|---|
| **Step** | **Annotation** |
| Step 1: Input dataset<br>**input:** *select at runtime* | Input list of possible tn insertions forward reads - Chromosome (c1), start (c2), stop (c3), coverage on TA (c4) |
| Step 2: Input dataset<br>**input:** *select at runtime* | Input list of possible tn insertions reverse reads - Chromosome (c1), start (c2), stop (c3), coverage on TA (c4) |
| Step 3: Join two Datasets<br>**Join:** Output dataset 'output' from step 1<br>**using column:** 2<br>**with:** Output dataset 'output' from step 2<br>**and column:** 2<br>**Keep lines of first input that do not join with second input:** No<br>**Keep lines of first input that are incomplete:** No<br>**Fill empty columns:** No<br>**Keep the header lines:** No | |
| Step 4: Compute<br>**Add expression:** c4+c8<br>**as a new column to:** Output dataset 'out_file1' from step 3<br>**Round result?**YES | |
| Step 5: Cut<br>**Cut columns:** c1,c2,c3,c9<br>**Delimited by:** Tab<br>**From:** Output dataset 'out_file1' from step 4 | Output = interval file containing chromosome (c1), Tn start (c2), Tn stop (c3), total coverage on TA (Fw and Rev reads starting on TA only) (c4) |
| **Output name**: TnList_TotalTAcoverage_x | |

| **Galaxy workflow 5: Add annotation to tn list** | |
|---|---|
| **Step** | **Annotation** |
| Step 1: Input dataset<br>Tn List: *select at runtime* | interval file Tn List (5 columns) chromosome (c1), Tn start (c2), Tn stop (c3), TnInsertion number (c4), total coverage on TA (Fw and Rev reads starting on TA only) (c5) |
| Step 2: Input dataset<br>Annotation file: *select at runtime* | annotation file (txt file organized as interval file, c1 (chrom), c2 (start), c3(end), other columns as wanted, text needs to be left aligned, no titles) |
| Step 3: Join<br>**Join:** Output dataset 'output' from step 1<br>**with:** Output dataset 'output' from step 2<br>**with min overlap:** 1<br>**Return:** All records of first dataset (fill null with ".") | Join based on Tn List file, keep all the records of the Tn List file |
| Step 4: Cut<br>**Cut columns:** c1-c5,c9-c10<br>**Delimited by:** Tab<br>**From:** Output dataset 'output' from step 3 | Cut wanted columns. Output = interval file with with 7 columns: chromosome (c1), Tn start (c2), Tn stop (c3), TnInsertion number (c4), total coverage on TA (Fw and Rev reads starting on TA only) (c5), locus gene (c6), name gene (c7) |

| | |
|---|---|
| Step 5: Sort<br>**Sort Query:** Output dataset 'out_file1' from step 4<br>**Number of header lines:** 0<br>**Column selections**<br>**Column selections 1**<br>**on column:** 4<br>**in:** Ascending order<br>**Flavor:** Natural/Version sort (-V)<br>**Output unique values:** True<br>**Ignore case:** False | In the step Join TnInsertions targeting overlapping genes got duplicated. For the further steps it is better to remove them. Otherwise they get duplicated again every time files are joined, leading to 8x the same TnInsertion in the final TnList. We remove these duplicates by performing a sort on c4 and only retaining the unique values. If this step is not performed here it can also be performed afterwards by removing the duplicates in Excel. |
| **Output name**: TnList_TotalTAcoverage_x_annotated<br>**Remark**: if the annotation file contains more columns you may need to adapt the cut step | |

| **Galaxy workflow 6a: Join 8 Datasets TAcoverage JoinEnd Start**<br>Workflow joins start (c2) of TnList with start (c2) of different file containing TA coverage info in c4. Cuts first 7 columns of final file + TA coverage info of all 8 joined datasets | |
|---|---|
| **Step** | **Annotation** |
| Step 1: Input dataset<br>**Input_TnList:** *select at runtime* | Input the interval file TnList_TotalTAcoverage_x_annotated containing 7 columns (otherwise ajust cut columns at the end) with start and stop in column 2 and 3. |
| Step 2: Input dataset<br>**Input_Dataset_A:** *select at runtime* | Dataset A, coverage file containing 4 columns: c1, c2 (start), c3 (stop), c4 (TAcoverage). |
| Step 3: Input dataset<br>**Input_Dataset_B:** *select at runtime* | Dataset B, idem as previous |
| Step 4: Input dataset<br>**Input_Dataset_C:** *select at runtime* | Dataset C, idem as previous |
| Step 5: Input dataset<br>**Input_Dataset_D:** *select at runtime* | Dataset D, idem as previous |
| Step 6: Input dataset<br>**Input_Dataset_E:** *select at runtime* | Dataset E, idem as previous |
| Step 7: Input dataset<br>**Input_Dataset_F:** *select at runtime* | Dataset F, idem as previous |
| Step 8: Input dataset<br>**Input_Dataset_G:** *select at runtime* | Dataset G, idem as previous |
| Step 9: Input dataset<br>**Input_Dataset_H:** *select at runtime* | Dataset H, idem as previous |
| Step 10: Join two Datasets<br>**Join:** Output dataset 'output' from step 1<br>**using column:** 2<br>**with:** Output dataset 'output' from step 2<br>**and column:** 2<br>**Keep lines of first input that do not join with second input:** Yes<br>**Keep lines of first input that are incomplete:** Yes<br>**Fill empty columns:** Yes<br>**Only fill unjoined rows:** Yes<br>**Fill Columns by:** Single fill value<br>**Fill value:** .<br>**Keep the header lines:** No | |
| Step 11: Join two Datasets<br>**Join:** Output dataset 'out_file1' from step 10<br>**using column:** 2<br>**with:** Output dataset 'output' from step 3<br>**and column:** 2<br>**Keep lines of first input that do not join with second input:** Yes<br>**Keep lines of first input that are incomplete:** Yes<br>**Fill empty columns:** Yes<br>**Only fill unjoined rows:** Yes<br>**Fill Columns by:** Single fill value<br>**Fill value:** .<br>**Keep the header lines:** No | |

| |
|---|
| <u>Step 12: Join two Datasets</u><br>**Join:** Output dataset 'out_file1' from step 11<br>**using column:** 2<br>**with:** Output dataset 'output' from step 4<br>**and column:** 2<br>**Keep lines of first input that do not join with second input:** Yes<br>**Keep lines of first input that are incomplete:** Yes<br>**Fill empty columns:** Yes<br>**Only fill unjoined rows:** Yes<br>**Fill Columns by:** Single fill value<br>**Fill value:** .<br>**Keep the header lines:** No |
| <u>Step 13: Join two Datasets</u><br>**Join:** Output dataset 'out_file1' from step 12<br>**using column:** 2<br>**with:** Output dataset 'output' from step 5<br>**and column:** 2<br>**Keep lines of first input that do not join with second input:** Yes<br>**Keep lines of first input that are incomplete:** Yes<br>**Fill empty columns:** Yes<br>**Only fill unjoined rows:** Yes<br>**Fill Columns by:** Single fill value<br>**Fill value:** .<br>**Keep the header lines:** No |
| <u>Step 14: Join two Datasets</u><br>**Join:** Output dataset 'out_file1' from step 13<br>**using column:** 2<br>**with:** Output dataset 'output' from step 6<br>**and column:** 2<br>**Keep lines of first input that do not join with second input:** Yes<br>**Keep lines of first input that are incomplete:** Yes<br>**Fill empty columns:** Yes<br>**Only fill unjoined rows:** Yes<br>**Fill Columns by:** Single fill value<br>**Fill value:** .<br>**Keep the header lines:** No |
| <u>Step 15: Join two Datasets</u><br>**Join:** Output dataset 'out_file1' from step 14<br>**using column:** 2<br>**with:** Output dataset 'output' from step 7<br>**and column:** 2<br>**Keep lines of first input that do not join with second input:** Yes<br>**Keep lines of first input that are incomplete:** Yes<br>**Fill empty columns:** Yes<br>**Only fill unjoined rows:** Yes<br>**Fill Columns by:** Single fill value<br>**Fill value:** .<br>**Keep the header lines:** No |
| <u>Step 16: Join two Datasets</u><br>**Join:** Output dataset 'out_file1' from step 15<br>**using column:** 2<br>**with:** Output dataset 'output' from step 8<br>**and column:** 2<br>**Keep lines of first input that do not join with second input:** Yes<br>**Keep lines of first input that are incomplete:** Yes<br>**Fill empty columns:** Yes<br>**Only fill unjoined rows:** Yes<br>**Fill Columns by:** Single fill value<br>**Fill value:** .<br>**Keep the header lines:** No |
| <u>Step 17: Join two Datasets</u><br>**Join:** Output dataset 'out_file1' from step 16<br>**using column:** 2 |

| | |
|---|---|
| **with:** Output dataset 'output' from step 9<br>**and column:** 2<br>**Keep lines of first input that do not join with second input:** Yes<br>**Keep lines of first input that are incomplete:** Yes<br>**Fill empty columns:** Yes<br>**Only fill unjoined rows:** Yes<br>**Fill Columns by:** Single fill value<br>**Fill value:** .<br>**Keep the header lines:** No | |
| Step 18: Cut<br>**Cut columns:**<br>c1,c2,c3,c4,c5,c6,c7,c11,c15,c19,c23,c27,c31,c35,c39<br>**Delimited by:** Tab<br>**From:** Output dataset 'out_file1' from step 17 | Output = interval file TnList containing chromosome (c1), Tn start (c2), Tn stop (c3), TnInsertion number (c4), total coverage on TA (Fw and Rev reads starting on TA only) (c5), locus gene (c6), name gene (c7), TA coverage dataset A-H (c8-c15) |
| **Output name**: TnList_TotalTAcoverage_x_A-H | |

| **Galaxy workflow 6b: Join 12 Datasets TAcoverage JoinEnd Start**<br>Annotation: Workflow joins start (c2) of TnList with start (c2) of different file containing TA coverage info in c4. Cuts first 7 columns of final file + TA coverage info of all 12 joined datasets | |
|---|---|
| **Step** | **Annotation** |
| Step 1: Input dataset<br>**Input_TnList:** *select at runtime* | Input the interval file Tn List TnList_TotalTAcoverage_x_annotated containing 7 columns (otherwise ajust cut columns at the end) with start and stop in column 2 and 3. |
| Step 2: Input dataset<br>**Input_Dataset_1:** *select at runtime* | Dataset 1, coverage file containing 4 columns: c1, c2 (start), c3 (stop), c4 (TAcoverage). |
| Step 3: Input dataset<br>**Input_Dataset_2:** *select at runtime* | Dataset 2, idem as previous |
| Step 4: Input dataset<br>**Input_Dataset_3:** *select at runtime* | Dataset 3, idem as previous |
| Step 5: Input dataset<br>**Input_Dataset_4:** *select at runtime* | Dataset 4, idem as previous |
| Step 6: Input dataset<br>**Input_Dataset_5:** *select at runtime* | Dataset 5, idem as previous |
| Step 7: Input dataset<br>**Input_Dataset_6:** *select at runtime* | Dataset 6, idem as previous |
| Step 8: Input dataset<br>**Input_Dataset_7:** *select at runtime* | Dataset 7, idem as previous |
| Step 9: Input dataset<br>**Input_Dataset_8:** *select at runtime* | Dataset 8, idem as previous |
| Step 10: Input dataset<br>**Input_Dataset_9:** *select at runtime* | Dataset 9, idem as previous |
| Step 11: Input dataset<br>**Input_Dataset_10:** *select at runtime* | Dataset 10, idem as previous |
| Step 12: Input dataset<br>**Input_Dataset_11:** *select at runtime* | Dataset 11, idem as previous |
| Step 13: Input dataset<br>**Input_Dataset_12:** *select at runtime* | Dataset 12, idem as previous |
| Step 14: Join two Datasets<br>**Join:** Output dataset 'output' from step 1<br>**using column:** 2<br>**with:** Output dataset 'output' from step 2<br>**and column:** 2<br>**Keep lines of first input that do not join with second input:** Yes<br>**Keep lines of first input that are incomplete:** Yes<br>**Fill empty columns:** Yes<br>**Only fill unjoined rows:** Yes<br>**Fill Columns by:** Single fill value<br>**Fill value:** .<br>**Keep the header lines:** No | |
| Step 15: Join two Datasets | |

| |
|---|
| **Join:** Output dataset 'out_file1' from step 14<br>**using column:** 2<br>**with:** Output dataset 'output' from step 3<br>**and column:** 2<br>**Keep lines of first input that do not join with second input:** Yes<br>**Keep lines of first input that are incomplete:** Yes<br>**Fill empty columns:** Yes<br>**Only fill unjoined rows:** Yes<br>**Fill Columns by:** Single fill value<br>**Fill value:** .<br>**Keep the header lines:** No |
| Step 16: Join two Datasets<br>**Join:** Output dataset 'out_file1' from step 15<br>**using column:** 2<br>**with:** Output dataset 'output' from step 4<br>**and column:** 2<br>**Keep lines of first input that do not join with second input:** Yes<br>**Keep lines of first input that are incomplete:** Yes<br>**Fill empty columns:** Yes<br>**Only fill unjoined rows:** Yes<br>**Fill Columns by:** Single fill value<br>**Fill value:** .<br>**Keep the header lines:** No |
| Step 17: Join two Datasets<br>**Join:** Output dataset 'out_file1' from step 16<br>**using column:** 2<br>**with:** Output dataset 'output' from step 5<br>**and column:** 2<br>**Keep lines of first input that do not join with second input:** Yes<br>**Keep lines of first input that are incomplete:** Yes<br>**Fill empty columns:** Yes<br>**Only fill unjoined rows:** Yes<br>**Fill Columns by:** Single fill value<br>**Fill value** .<br>**Keep the header lines:** No |
| Step 18: Join two Datasets<br>**Join:** Output dataset 'out_file1' from step 17<br>**using column:** 2<br>**with:** Output dataset 'output' from step 6<br>**and column:** 2<br>**Keep lines of first input that do not join with second input:** Yes<br>**Keep lines of first input that are incomplete:** Yes<br>**Fill empty columns:** Yes<br>**Only fill unjoined rows:** Yes<br>**Fill Columns by:** Single fill value<br>**Fill value:** .<br>**Keep the header lines:** No |
| Step 19: Join two Datasets<br>**Join:** Output dataset 'out_file1' from step 18<br>**using column:** 2<br>**with:** Output dataset 'output' from step 7<br>**and column:** 2<br>**Keep lines of first input that do not join with second input:** Yes<br>**Keep lines of first input that are incomplete:** Yes<br>**Fill empty columns:** Yes<br>**Only fill unjoined rows:** Yes<br>**Fill Columns by:** Single fill value<br>**Fill value:** .<br>**Keep the header lines:** No |
| Step 20: Join two Datasets<br>**Join:** Output dataset 'out_file1' from step 19<br>**using column:** 2<br>**with:** Output dataset 'output' from step 8 |

| |
|---|
| **and column:** 2<br>**Keep lines of first input that do not join with second input:** Yes<br>**Keep lines of first input that are incomplete:** Yes<br>**Fill empty columns:** Yes<br>**Only fill unjoined rows:** Yes<br>**Fill Columns by:** Single fill value<br>**Fill value:** .<br>**Keep the header lines:** No |
| Step 21: Join two Datasets<br>**Join:** Output dataset 'out_file1' from step 20<br>**using column:** 2<br>**with:** Output dataset 'output' from step 9<br>**and column:** 2<br>**Keep lines of first input that do not join with second input:** Yes<br>**Keep lines of first input that are incomplete:** Yes<br>**Fill empty columns:** Yes<br>**Only fill unjoined rows:** Yes<br>**Fill Columns by:** Single fill value<br>**Fill value:** .<br>**Keep the header lines:** No |
| Step 22: Join two Datasets<br>**Join:** Output dataset 'out_file1' from step 21<br>**using column:** 2<br>**with:** Output dataset 'output' from step 10<br>**and column:** 2<br>**Keep lines of first input that do not join with second input:** Yes<br>**Keep lines of first input that are incomplete:** Yes<br>**Fill empty columns:** Yes<br>**Only fill unjoined rows:** Yes<br>**Fill Columns by:** Single fill value<br>**Fill value:** .<br>**Keep the header lines:** No |
| Step 23: Join two Datasets<br>**Join:** Output dataset 'out_file1' from step 22<br>**using column:** 2<br>**with:** Output dataset 'output' from step 11<br>**and column:** 2<br>**Keep lines of first input that do not join with second input:** Yes<br>**Keep lines of first input that are incomplete:** Yes<br>**Fill empty columns:** Yes<br>**Only fill unjoined rows:** Yes<br>**Fill Columns by:** Single fill value<br>**Fill value:** .<br>**Keep the header lines:** No |
| Step 24: Join two Datasets<br>**Join:** Output dataset 'out_file1' from step 23<br>**using column:** 2<br>**with:** Output dataset 'output' from step 12<br>**and column:** 2<br>**Keep lines of first input that do not join with second input:** Yes<br>**Keep lines of first input that are incomplete:** Yes<br>**Fill empty columns:** Yes<br>**Only fill unjoined rows:** Yes<br>**Fill Columns by:** Single fill value<br>**Fill value:** .<br>**Keep the header lines:** No |
| Step 25: Join two Datasets<br>**Join:** Output dataset 'out_file1' from step 24<br>**using column:** 2<br>**with:** Output dataset 'output' from step 13<br>**and column:** 2<br>**Keep lines of first input that do not join with second input:** Yes<br>**Keep lines of first input that are incomplete:** Yes |

| | |
|---|---|
| **Fill empty columns:** Yes<br>**Only fill unjoined rows:** Yes<br>**Fill Columns by:** Single fill value<br>**Fill value:** .<br>**Keep the header lines:** No | |
| Step 26: Cut<br>**Cut columns:**<br>c1,c2,c3,c4,c5,c6,c7,c11,c15,c19,c23,c27,c31,c35,c39,c43,c47,c51,c55<br>**Delimited by:** Tab<br>**From:** Output dataset 'out_file1' from step 25 | Output = interval file containing chromosome (c1), Tn start (c2), Tn stop (c3), TnInsertion number (c4), total coverage on TA (Fw and Rev reads starting on TA only) (c5), locus gene (c6), name gene (c7), TA coverage dataset 1-12 (c8-c19) |
| **Output name**: TnList_TotalTAcoverage_x_1-12 | |

| **Galaxy workflow 7: Join 1-12 and A-H coverage files** | |
|---|---|
| **Step** | **Annotation** |
| Step 1: Input dataset<br>**input:** *select at runtime* | TnList_TotalTAcoverage_x_annotated |
| Step 2: Input dataset<br>**input:** *select at runtime* | TnList_TotalTAcoverage_x_1-12 |
| Step 3: Input dataset<br>**input:** *select at runtime* | TnList_TotalTAcoverage_x_A-H |
| Step 4: Add column<br>**Add this value:** Empty.<br>**to Dataset:** Output dataset 'output' from step 1<br>**Iterate?** NO | Add empty column |
| Step 5: Join two Datasets<br>**Join:** Output dataset 'out_file1' from step 4<br>**using column:** 2<br>**with:** Output dataset 'output' from step 2<br>**and column:** 2<br>**Keep lines of first input that do not join with second input:** No<br>**Keep lines of first input that are incomplete:** No<br>**Fill empty columns:** No<br>**Keep the header lines:** No | Join with coverage file pools 1-12 |
| Step 6: Add column<br>**Add this value:** Empty.<br>**to Dataset:** Output dataset 'output' from step 1<br>**Iterate?** NO | Add empty column |
| Step 7: Join two Datasets<br>**Join:** Output dataset 'out_file1' from step 6<br>**using column:** 2<br>**with:** Output dataset 'output' from step 3<br>**and column:** 2<br>**Keep lines of first input that do not join with second input:** No<br>**Keep lines of first input that are incomplete:** No<br>**Fill empty columns:** No<br>**Keep the header lines:** No | Join with coverage file pools 1-12 |
| Step 8: Cut<br>**Cut columns:**<br>c1,c2,c3,c4,c5,c6,c7,c8,c16,c17,c18,c19,c20,c21,c22,c23,c24,c25,c26,c27,c28,c36,c37,c38,c39,c40,c41,c42,c43<br>**Delimited by:** Tab<br>**From:** Output dataset 'out_file1' from step 7 | Cut wanted columns. Output = an interval file with 7 columns mutant and gene information (c1 to c7), c8=empty, 12 columns containing read counts of each mutant per Cartesian pool 1-12 (c9 to c20), c21=empty, 8 columns containing read counts of each mutant per Cartesian pool A-H (c22 to c29) |
| **Output name**: TnList_x | |

| **Galaxy workflow 8: Add chr/start/end/name and coverages to FinalTnList** | |
|---|---|
| **Step** | **Annotation** |
| Step 1: Input dataset<br>**input:** *select at runtime* | TnList_TotalTAcoverage_x_annotated (7 columns) |
| Step 2: Input dataset<br>**input:** *select at runtime* | FinalTnList (7 columns) |

| | |
|---|---|
| Step 3: Input dataset<br>**input:** *select at runtime* | TnList_Z_normalized.txt (29 columns) |
| Step 4: Input dataset<br>**input:** *select at runtime* | TnList_XY_normalized.txt (29 columns) |
| Step 5: Join two Datasets<br>**Join:** Output dataset 'output' from step 1<br>**using column:** 4<br>**with:** Output dataset 'output' from step 2<br>**and column**: 1<br>**Keep lines of first input that do not join with second input :** Yes<br>**Keep lines of first input that are incomplete:** Yes<br>**Fill empty columns**: Yes<br>**Only fill unjoined rows:** No<br>**Fill Columns by:** Single fill value<br>**Fill value :** .<br>**Keep the header lines:** No | Join with FinalTnList.txt |
| Step 6: Cut<br>**Cut columns:** c1-c4,c6-c7,c10-c14<br>**Delimited by:** Tab<br>**From:** Output dataset 'out_file1' from step 5 | Cut wanted columns from the file. Output = an interval file with 6 columns mutant and gene information (c1 to c6) and 5 columns position information. |
| Step 7: Join two Datasets<br>**Join:** Output dataset 'output' from step 6<br>**using column:** 4<br>**with:** Output dataset 'output' from step 3<br>**and column**: 4<br>**Keep lines of first input that do not join with second input :** Yes<br>**Keep lines of first input that are incomplete:** Yes<br>**Fill empty columns**: Yes<br>**Only fill unjoined rows:** No<br>**Fill Columns by:** Single fill value<br>**Fill value :** .<br>**Keep the header lines:** No | Join with coverage for Z coordinate (TnList_Z_normalized.txt) |
| Step 8: Cut<br>**Cut columns:** c1-c11,c19-c40<br>**Delimited by:** Tab<br>**From:** Output dataset 'out_file1' from step 7 | Cut wanted columns from the file. Output = an interval file with 6 columns mutant and gene information (c1 to c6) and 5 columns position information (c7 to c11) and the rest of the columns with the coverage information for the Z coordinate (c12-c33) |
| Step 9: Join two Datasets<br>**Join:** Output dataset 'output' from step 8<br>**using column:** 4<br>**with:** Output dataset 'output' from step 4<br>**and column**: 4<br>**Keep lines of first input that do not join with second input :** Yes<br>**Keep lines of first input that are incomplete:** Yes<br>**Fill empty columns**: Yes<br>**Only fill unjoined rows:** No<br>**Fill Columns by:** Single fill value<br>**Fill value :** .<br>**Keep the header lines:** No | Join with coverage for XY coordinate (TnList_XY_normalized.txt) |
| Step 10: Cut<br>**Cut columns:** c1-c33,c41-c62<br>**Delimited by:** Tab<br>**From:** Output dataset 'out_file1' from step 9 | Cut wanted columns from the file. Output = an interval file with 6 columns mutant and gene information (c1 to c6) and 5 columns position information (c7 to c11) and the rest of the columns with the coverage information for the Z and XY coordinates (c12-c55) |
| **Output name**: FinalTnList! | |

| **Galaxy workflow 9: Extracting TnList_DUPLICATED** | |
|---|---|
| **Step** | **Annotation** |
| Step 1: Input dataset<br>**TnList_ALL:** *select at runtime* | TnList containing all TnInsertions targeting one-sided and two-sided unique regions and duplicated regions. All reads that map to the genome were included in the analysis to |

| | |
|---|---|
| | generate this list. |
| <u>Step 2: Input dataset</u><br>**TnList_UNIQUE:** *select at runtime* | TnList_UNIQUE containing all TnInsertions targeting two-sided unique regions. Reads that do not map uniquely to the genome were discarded during the data-analysis to create this list. |
| <u>Step 3: Compare two Datasets</u><br>**Compare** Output dataset 'output' from step 1<br>**Using column** 2<br>**against** Output dataset 'output' from step 2<br>**and column** 2<br>**To find** Non Matching rows of 1st dataset | Extract TnList_DUPLICATED containing TnInsertions targeting duplicated regions and one-sided unique regions. Creating TnList_DUP of 55 columns. Output = an interval file with 6 columns mutant and gene information (c1 to c6) and 5 columns position information (c7 to c11) and the rest of the columns with the coverage information for the Z and XY coordinates (c12-c55). |
| <u>Step 4: Filter</u><br>**Filter** Output dataset 'out_file1' from step 3<br>**With following condition** c12>=60<br>**Number of header lines to skip** 0 | Deletes all TnInsertions with a total TA coverage <60 |
| <u>Step 5: Sort</u><br>**Sort Query** Output dataset 'out_file1' from step 4<br>**Number of header lines** 0<br>**Column selections**<br>**Column selections 1**<br>on column: 12<br>in: Ascending order<br>Flavor: Fast numeric sort (-n)<br>**Column selections 2**<br>on column: 13<br>in: Ascending order<br>Flavor: Fast numeric sort (-n)<br>**Column selections 3**<br>on column: 14<br>in: Ascending order<br>Flavor: Fast numeric sort (-n)<br>**Column selections 4**<br>on column: 15<br>in: Ascending order<br>Flavor: Fast numeric sort (-n)<br>**Column selections 5**<br>on column: 16<br>in: Ascending order<br>Flavor: Fast numeric sort (-n)<br>**Column selections 6**<br>on column: 17<br>in: Ascending order<br>Flavor: Fast numeric sort (-n)<br>**Column selections 7**<br>on column: 18<br>in: Ascending order<br>Flavor: Fast numeric sort (-n)<br>**Column selections 8**<br>on column: 19<br>in: Ascending order<br>Flavor: Fast numeric sort (-n)<br>**Column selections 9**<br>on column: 20<br>in: Ascending order<br>Flavor: Fast numeric sort (-n)<br>**Column selections 10**<br>on column: 21<br>in: Ascending order<br>Flavor: Fast numeric sort (-n)<br>**Column selections 11**<br>on column: 22<br>in: Ascending order | The TnList is sorted for the TA coverages |

| |
|---|
| Flavor: Fast numeric sort (-n)<br>**Column selections 12**<br>on column: 23<br>in: Ascending order<br>Flavor: Fast numeric sort (-n)<br>**Column selections 13**<br>on column: 24<br>in: Ascending order<br>Flavor: Fast numeric sort (-n)<br>**Column selections 14**<br>on column: 26<br>in: Ascending order<br>Flavor: Fast numeric sort (-n)<br>**Column selections 15**<br>on column: 27<br>in :Ascending order<br>Flavor: Fast numeric sort (-n)<br>**Column selections 16**<br>on column: 28<br>in: Ascending order<br>Flavor: Fast numeric sort (-n)<br>**Column selections 17**<br>on column: 29<br>in: Ascending order<br>Flavor: Fast numeric sort (-n)<br>**Column selections 18**<br>on column: 30<br>in: Ascending order<br>Flavor: Fast numeric sort (-n)<br>**Column selections 19**<br>on column: 31<br>in: Ascending order<br>Flavor: Fast numeric sort (-n)<br>**Column selections 20**<br>on column: 32<br>in: Ascending order<br>Flavor: Fast numeric sort (-n)<br>**Column selections 21**<br>on column: 33<br>in: Ascending order<br>Flavor: Fast numeric sort (-n)<br>**Column selections 22**<br>on column: 35<br>in: Ascending order<br>Flavor: Fast numeric sort (-n)<br>**Column selections 23**<br>on column: 36<br>in: Ascending order<br>Flavor: Fast numeric sort (-n)<br>**Column selections 24**<br>on column: 37<br>in: Ascending order<br>Flavor: Fast numeric sort (-n)<br>**Column selections 25**<br>on column: 38<br>in: Ascending order<br>Flavor: Fast numeric sort (-n)<br>**Column selections 26**<br>on column: 39<br>in: Ascending order<br>Flavor: Fast numeric sort (-n)<br>**Column selections 27**<br>on column: 40 |

| | |
|---|---|
| in: Ascending order<br>Flavor: Fast numeric sort (-n)<br>**Column selections 28**<br>on column: 41<br>in: Ascending order<br>Flavor: Fast numeric sort (-n)<br>**Column selections 29**<br>on column: 42<br>in: Ascending order<br>Flavor: Fast numeric sort (-n)<br>**Column selections 30**<br>on column: 43<br>in: Ascending order<br>Flavor: Fast numeric sort (-n)<br>**Column selections 31**<br>on column: 44<br>in: Ascending order<br>Flavor: Fast numeric sort (-n)<br>**Column selections 32**<br>on column: 45<br>in: Ascending order<br>Flavor: Fast numeric sort (-n)<br>**Column selections 33**<br>on column: 46<br>in: Ascending order<br>Flavor: Fast numeric sort (-n)<br>**Column selections 34**<br>on column: 48<br>in: Ascending order<br>Flavor: Fast numeric sort (-n)<br>**Column selections 35**<br>on column: 49<br>in: Ascending order<br>Flavor: Fast numeric sort (-n)<br>**Column selections 36**<br>on column: 50<br>in: Ascending order<br>Flavor: Fast numeric sort (-n)<br>**Column selections 37**<br>on column: 51<br>in: Ascending order<br>Flavor: Fast numeric sort (-n)<br>**Column selections 38**<br>on column: 52<br>in: Ascending order<br>Flavor: Fast numeric sort (-n)<br>**Column selections 39**<br>on column: 53<br>in: Ascending order<br>Flavor: Fast numeric sort (-n)<br>**Column selections 40**<br>on column: 54<br>in: Ascending order<br>Flavor: Fast numeric sort (-n)<br>**Column selections 41**<br>on column: 55<br>in: Ascending order<br>Flavor: Fast numeric sort (-n)<br>**Output unique values:** False<br>**Ignore case:** False | |
| Step 6: Unique lines<br>**File to scan for unique values:** Output dataset 'outfile' from step 5 | All duplicated lines are deleted from the TnList_DUP, while counting the number of occurrences of each unique line. |

| | |
|---|---|
| **Do you want to group each unique group?** No<br>**Counting number of occurrences:** True<br>**Only print duplicate lines:** False<br>**Only print unique lines:** False<br>**Ignore differences in case when comparing:** False<br>**Avoid comparing the first N fields:** 11 | |
| Step 7: Unique lines<br>**File to scan for unique values:** Output dataset 'outfile' from step 5<br>**Do you want to group each unique group?** Yes<br>Output all lines, and delimit each unique group<br>Output a delimiter around each group of unique items<br>**Ignore differences in case when comparing:** False<br>**Avoid comparing the first N fields:** 11 | All TnInsertions are grouped according to their coverage profile. Matching TnInsertions that target duplicated regions together. Creating TnList_DUP_grouped. |
| Step 8: Cut<br>**Cut columns:** c2-c5,c1,c6-c56<br>**Delimited by**: Tab<br>**From**: Output dataset 'outfile' from step 6 | The columns are reorganized in the TnList. Creating TnList_DUP_condensed of 56 columns Output = an interval file with 6 columns mutant and gene information (c1 to c7) and 5 columns position information (c8 to c12) and the rest of the columns with the coverage information for the Z and XY coordinates (c13-c56) |
| **Output names**: TnList_DUP, TnList_DUP_grouped, TnList_DUP_condensed | |

### 4. BioPerl CP-CSeq algorithm

```
##################################################################
# CP-CSeq protocol.
# Yvan Saeys wrote the BioPerl algorithm for the automated location assignment (K. Vandewalle et al., Nature Communications, 2015)(1). This algorithm differs from the original one by exchanging "AP" for "XY" and "TP" for "Z". A variable "librarySet" was added to be able to differentiate between different library sets. In addition, the output was renamed and reorganized. Previously, the TnList and the statistical summary were printed to the terminal and only the file "uniquesnew.text" or now "TnListOrderedByPlate.txt" was generated as output. Now the algorithm also generates the two files "TnListOrderedByMutant.txt" and "Summary.txt". A piece of code (compiled by Laurent Schindfessel) was added to clean up and reorder "TnListOrderedByMutant.txt" in order to create "FinalTnList.txt", in which the location of the mutants can be more easily discerned. In the subroutine, the new arguments 'exaequo' were introduced for the cases where there are equal values in the coverages (compiled by Laurent Schindfessel).

#!/usr/local/bin/perl -w

open(FH, '>', 'TnListOrderedByMutant.txt') or die "cannot open file";
select FH;

use strict;
use warnings;

#variable to declare with which library set you are working if you have more than one library set (one set consists of 1x 96 96-well plates). Default the variable is set at "I", if you have for example a second set you can put this variable at "II". This allows you to merge the two library sets together afterwards.
my $librarySet = "I";

#input files: the two tabular (.txt) input files (TnList_XY_normalized.txt and TnList_Z_normalized.txt) which were generated, each containing 29 columns in total, 7 columns with mutant and gene information (c1 to c7), c8=empty, 12 columns containing read counts of each mutant per Cartesian pool 1-12 (c9 to c20), c21=empty, 8 columns containing read counts of each mutant per Cartesian pool A-H (c22 to c29). These files need to be located in the same map as the algorithm is when you perform.
my $XYfile="TnList_XY_normalized.txt";
my $Zfile="TnList_Z_normalized.txt";

#Some thresholds are declared
#Threshold below which to remove counts
my $min_threshold=15;
#Other matching thresholds
```

```
my $split_threshold=1000;
my $lower_ratio=3; # if this ratio is set to 1, then subroutine might not make a distinction between equal coverages
my $upper_ratio=1.5; # if this ratio is set to 1, then subroutine might not make a distinction between equal coverages

my %XYokhash;
my %Zokhash;
my $totalidentifications=0;

open(XY,"<$XYfile");
open(Z,"<$Zfile");

#First skip the header lines
$_=<XY>;
$_=<Z>;

my $numok=0;
my $numXYok=0;
my $numunique=0;
my $numZok=0;

my $numcase1=0;
my $impossible=0;
my $rest=0;

my %stack_unique;
my %stack_XYok;
my %stack_Zok;
my %stack_heuristic;

#For every row in the XY file split the row on tab and define where the column TnInsertion and Locus is located. Then the same is done for the same row in Z file. Following this, a sanity check is performed to see if the TnInsertion and Locus are the same for the same row in the two files. If everything is okay a count is added to $numok.
while(<XY>){
  my $line=$_;
  chomp($line);
  my @valsXY=split/\t/,$line;
  my $TnInsertion_XY=$valsXY[3];
  my $Locus_BCG_XY=$valsXY[5];

  $_=<Z>;
  $line=$_;
  chomp($line);
  my @valsZ=split/\t/,$line;
  my $TnInsertion_Z=$valsZ[3];
  my $Locus_BCG_Z=$valsZ[5];

  #Sanity check to see if they are the same
  if( ($TnInsertion_XY eq $TnInsertion_Z) &&
      ($Locus_BCG_XY eq $Locus_BCG_Z)){
     #All ok
     $numok++;
      }
   else{
     die;
   }

#Now the calculations can be performed

#In the following code, one checks if the coverage is integer and exceeds the min_threshold. If this is true the coverage is kept in the appropriate array. Four different arrays are created. @XY1 is an empty array for maximum 12 values for the 12 columns (9-21) of the XY plate, @XY2 is empty array for maximum 8 values for the 8 rows (22-29) of the XY plate. Analogously, we have @Z1 and @Z2 for the Z plate. In the ideal case, only one value remains for each
```

array of a plate (e.g. @XY1=1), meaning that the position of the mutant can be uniquely determined in the plate. This is checked and if this is true $XYok or $Zok is put at 1.

```
  #First do the XY check
  my $XYok=0;
  my @XY1;
  for(my $i=0;$i<12;$i++){
    if(is_integer($valsXY[8+$i])){
   if($valsXY[8+$i]>$min_threshold){
        push(@XY1,$valsXY[8+$i])
      }
    }
  }
  my @XY2;
  for(my $i=0;$i<8;$i++){
    if(is_integer($valsXY[21+$i])){
   if($valsXY[21+$i]>$min_threshold){
        push(@XY2,$valsXY[21+$i])
      }
    }
  }

  if( (scalar(@XY1)==1) &&
      (scalar(@XY2)==1)){
   $XYok=1; #ideal case in which only one value remains in each array
  }

  #Now do the Z check in the same way
  my $Zok=0;
  my @Z1;
  for(my $i=0;$i<12;$i++){
    if(is_integer($valsZ[8+$i])){
   if($valsZ[8+$i]>$min_threshold){
        push(@Z1,$valsZ[8+$i])
      }
    }
  }
  my @Z2;
  for(my $i=0;$i<8;$i++){
    if(is_integer($valsZ[21+$i])){
   if($valsZ[21+$i]>$min_threshold){
        push(@Z2,$valsZ[21+$i])
      }
    }
  }

  if( (scalar(@Z1)==1) &&
      (scalar(@Z2)==1)){
   $Zok=1; #ideal case in which only one value remains in each array
  }

#Now we know whether we have one unique value in @XY1, @XY2, @Z1 and @Z2. We can go over the different cases to assign the positions
  ##Case 1##: at least one of the 4 arrays has less than one count
  #Once there is no coverage in 1 array (@XY1, @XY2, @Z1 or @Z2) for a mutant, we can impossibly detect the location for that mutant
  #Count number of cases for which we can impossibly detect the plate
  if( (scalar(@XY1) < 1) or
      (scalar(@XY2) < 1) or
      (scalar(@Z1) < 1) or
      (scalar(@Z2) < 1)){
   $impossible++;
   }
   ##Case 2##: all the 4 arrays have one count ($XYok=1 and $Zok=1)
```

```
   #If every array has one value and the identification of the mutant is unique
  elsif($XYok && $Zok){

   $totalidentifications++;
   $numunique++;

   (my $XYid1, my $XYid2, my $Zid1, my $Zid2)=get_plate_id(\@valsXY,\@valsZ,$XY1[0],$XY2[0],$Z1[0],$Z2[0],0,0,0,0);

   print "Unique\t".($TnInsertion_XY)."\t".( $Locus_BCG_Z)."\tXY $XYid1".($XYid2)."\t"."Z $Zid1".($Zid2)."\n";
   push(@{$stack_unique{$XYid1.$XYid2}{$Zid1.$Zid2}},"$TnInsertion_XY\t$Locus_BCG_Z");
  }
  ##Case 3##: unique identification for XY ($XYok=1), not for Z (several values for Z)
  #If we have one value for @XY1 and @XY2 than XY is OK, subsequently we need to look at the different possibilities for the Z coordinate.
  elsif($XYok){
   my $numpos=0;

   #First sort the values in @Z1 and @Z2, from big to small. The biggest values are put first in the array and are also the most probable
   my @Z1s=sort{$b<=>$a} @Z1;
   my @Z2s=sort{$b<=>$a} @Z2;

   ###Case 3a###: 1 value in @Z1, several values in @Z2
   #Check if first element of Z2s is comparable, and the next element is significantly less
   #Check if the first element is > split_threshold, if yes check if the fraction of the first and second element is > upper_ratio (least stringent), if yes we have a XYok_unique identification, if no we have a XYok_nonunique identification.
   #If the first element is not bigger than the split_threshold, it is checked if the fraction of the first and second element is > lower_ratio (more stringent), if yes we have a XYok_unique identification, if no we have a XYok_nonunique identification.
   if((scalar(@Z1s)==1) && (scalar(@Z2s)>1)){
      my $fraction=$Z2s[0]/$Z2s[1];
   if($Z2s[0]>=$split_threshold){
     if($fraction >  $upper_ratio){
      $numpos=1;
         (my $XYid1, my $XYid2, my $Zid1, my $Zid2)=get_plate_id(\@valsXY,\@valsZ,$XY1[0],$XY2[0],$Z1s[0],$Z2s[0],0,0,0,0);

         print "XYok_unique\t".($TnInsertion_XY)."\t".( $Locus_BCG_Z)."\tXY $XYid1".($XYid2)."\t"."Z $Zid1".($Zid2)."\n";
         push(@{$stack_XYok{$XYid1.$XYid2}{$Zid1.$Zid2}},"$TnInsertion_XY\t$Locus_BCG_Z");
         $totalidentifications++;
         $numXYok++;
        }
     else{
       $numpos=scalar(@Z2s);
      if($numpos < 5){
          print "XYok_nonunique\t".($TnInsertion_XY)."\t".( $Locus_BCG_Z);
      for(my $i=0;$i<$numpos;$i++){
        (my $XYid1, my $XYid2, my $Zid1, my $Zid2)=get_plate_id(\@valsXY,\@valsZ,$XY1[0],$XY2[0],$Z1s[0],$Z2s[$i],0,0,0,$i);
        print "\tXY $XYid1".($XYid2)."\t"."Z $Zid1".($Zid2);
      }
      print "\n";
      $totalidentifications++;
      $numXYok++;
          }
      else{
        $impossible++;
      }
     }
      }
```

```
    else{
     if($fraction > $lower_ratio){
       $numpos=1;
         (my $XYid1, my $XYid2, my $Zid1, my $Zid2)=get_plate_id(\@valsXY,\@valsZ,$XY1[0],$XY2[0],$Z1s[0],$Z2s[0],0,0,0,0);

         print "XYok_unique\t".($TnInsertion_XY)."\t".( $Locus_BCG_Z)."\tXY $XYid1".($XYid2)."\t"."Z $Zid1".($Zid2)."\n";
         push(@{$stack_XYok{$XYid1.$XYid2}{$Zid1.$Zid2}},"$TnInsertion_XY\t$Locus_BCG_Z");
         $totalidentifications++;
         $numXYok++;

        }
      else{
      $numpos=scalar(@Z2s);
      if($numpos < 5){
             print "XYok_nonunique\t".($TnInsertion_XY)."\t".( $Locus_BCG_Z);
       for(my $i=0;$i<$numpos;$i++){
         (my $XYid1, my $XYid2, my $Zid1, my $Zid2)=get_plate_id(\@valsXY,\@valsZ,$XY1[0],$XY2[0],$Z1s[0],$Z2s[$i],0,0,0,$i);
          print "\tXY $XYid1".($XYid2)."\t"."Z $Zid1".($Zid2);
       }
       print "\n";
       $totalidentifications++;
       $numXYok++;
           }
      else{
        $impossible++;
      }
      }
    }
    }
    ###Case 3b###: several values in @Z1, one value in @Z2
    #Check if first element of Z1s is comparable, and the next element is significantly less
    #Check if the first element is > split_threshold, if yes check if the fraction of the first and second element is > upper_ratio (least stringent), if yes we have a XYok_unique identification, if no we have a XYok_nonunique identification.
    #If the first element is not bigger than the split threshold, it is checked if the fraction of the first and second element is > lower_ratio (more stringent), if yes we have a XYok_unique identification, if no we have a XYok_nonunique identification.
    elsif((scalar(@Z2s)==1) && (scalar(@Z1s)>1)){
      my $fraction=$Z1s[0]/$Z1s[1];
    if($Z1s[0]>=$split_threshold){
     if($fraction >  $upper_ratio){
      $numpos=1;
           (my $XYid1, my $XYid2, my $Zid1, my $Zid2)=get_plate_id(\@valsXY,\@valsZ,$XY1[0],$XY2[0],$Z1s[0],$Z2s[0],0,0,0,0);

           print "XYok_unique\t".($TnInsertion_XY)."\t".( $Locus_BCG_Z)."\tXY $XYid1".($XYid2)."\t"."Z $Zid1".($Zid2)."\n";
           push(@{$stack_XYok{$XYid1.$XYid2}{$Zid1.$Zid2}},"$TnInsertion_XY\t$Locus_BCG_Z");
           $totalidentifications++;
           $numXYok++;

        }
     else{
      $numpos=scalar(@Z1s);
      if($numpos < 5){
            print "XYok_nonunique\t".($TnInsertion_XY)."\t".( $Locus_BCG_Z);
      for(my $i=0;$i<$numpos;$i++){
        (my $XYid1, my $XYid2, my $Zid1, my $Zid2)=get_plate_id(\@valsXY,\@valsZ,$XY1[0],$XY2[0],$Z1s[$i],$Z2s[0],0,0,$i,0);
         print "\tXY $XYid1".($XYid2)."\t"."Z $Zid1".($Zid2);
```

```
        }
        print "\n";
        $totalidentifications++;
        $numXYok++;
            }
       else{
          $impossible++;
       }

      }
        }
     else{
       if($fraction > $lower_ratio){
         $numpos=1;
            (my $XYid1, my $XYid2, my $Zid1, my $Zid2)=get_plate_id(\@valsXY,\@valsZ,$XY1[0],$XY2[0],$Z1s[0],$Z2s[0],0,0,0,0);

            print "XYok_unique\t".($TnInsertion_XY)."\t".( $Locus_BCG_Z)."\tXY $XYid1".($XYid2)."\t"."Z $Zid1".($Zid2)."\n";
            push(@{$stack_XYok{$XYid1.$XYid2}{$Zid1.$Zid2}},"$TnInsertion_XY\t$Locus_BCG_Z");
            $totalidentifications++;
            $numXYok++;
          }
       else{
       $numpos=scalar(@Z1s);
       if($numpos < 5){
              print "XYok_nonunique\t".($TnInsertion_XY)."\t".( $Locus_BCG_Z);
        for(my $i=0;$i<$numpos;$i++){
          (my $XYid1, my $XYid2, my $Zid1, my $Zid2)=get_plate_id(\@valsXY,\@valsZ,$XY1[0],$XY2[0],$Z1s[$i],$Z2s[0],0,0,$i,0);
           print "\tXY $XYid1".($XYid2)."\t"."Z $Zid1".($Zid2);
        }
        print "\n";
        $totalidentifications++;
        $numXYok++;
            }
       else{
          $impossible++;
       }
       }
     }
    }
    ###Case 3c###: several values in @Z1 AND in @Z2
    #Check for both if the first (biggest) value is >split_threshold AND >upper_ratio OR < split_threshold AND >lower_ratio. If true we have XYok_unique identification, if false we have an impossible case.
     elsif((scalar(@Z1s)>1) && (scalar(@Z2s)>1)){
        my $fraction1=$Z1s[0]/$Z1s[1];
        my $fraction2=$Z2s[0]/$Z2s[1];
        my $pass1=0;
     my $pass2=0;
     if($Z1s[0]>=$split_threshold){
       if($fraction1 > $upper_ratio){
        $pass1=1;
          }
        }
     else{
       if($fraction1 > $lower_ratio){
         $pass1=1;
           }
     }
     if($Z2s[0]>=$split_threshold){
       if($fraction2 > $upper_ratio){
        $pass2=1;
```

```
      }
    }
  else{
   if($fraction2 > $lower_ratio){
     $pass2=1;
      }
  }
     if($pass1 && $pass2){
    $numpos=1;
       (my $XYid1, my $XYid2, my $Zid1, my $Zid2)=get_plate_id(\@valsXY,\@valsZ,$XY1[0],$XY2[0],$Z1s[0],$Z2s[0],0,0,0,0);

       print "XYok_unique\t".($TnInsertion_XY)."\t".( $Locus_BCG_Z)."\tXY $XYid1".($XYid2)."\t"."Z $Zid1".($Zid2)."\n";
       push(@{$stack_XYok{$XYid1.$XYid2}{$Zid1.$Zid2}},"$TnInsertion_XY\t$Locus_BCG_Z");
       $totalidentifications++;
       $numXYok++;

  }
  else{
    $numpos=scalar(@Z1s)*scalar(@Z2s);
    $impossible++;
  }
 }
 else{
 die "Other case ".(scalar(@Z1s))." ".(scalar(@Z2s))." "."\n";
 }
  if(defined($XYokhash{$numpos})){
  $XYokhash{$numpos}=$XYokhash{$numpos}+1;
  }
  else{
  $XYokhash{$numpos}=1;
  }
}
##Case 4##: unique identification for Z ($Zok=1), not for XY (several values for XY)
#If we have one value for @Z1 and @Z2 than Z is OK, subsequently we need to look at the different possibilities for the XY coordinate.
elsif($Zok){

  my $numpos=0;

  #First sort the values in @XY1 and @XY2, from big to small. The biggest values are put first in the array and are also the most probable
  my @XY1s=sort{$b<=>$a} @XY1;
  my @XY2s=sort{$b<=>$a} @XY2;

  ###Case 4a###: 1 value in @XY1, several values in @XY2
  #Check if first element of XY2s is comparable, and the next element is significantly less
  #Check if the first element is > split_threshold, if yes check if the fraction of the first and second element is > upper_ratio (least stringent), if yes we have a Zok_unique identification, if no we have a Zok_nonunique identification.
  #If the first element is not bigger than the split threshold, it is checked if the fraction of the first and second element is > lower_ratio (more stringent), if yes we have a Zok_unique identification, if no we have a Zok_nonunique identification.
  if((scalar(@XY1s)==1) && (scalar(@XY2s)>1)){
     my $fraction=$XY2s[0]/$XY2s[1];
  if($XY2s[0]>=$split_threshold){
   if($fraction >  $upper_ratio){
    $numpos=1;
    (my $XYid1, my $XYid2, my $Zid1, my $Zid2)=get_plate_id(\@valsXY,\@valsZ,$XY1s[0],$XY2s[0],$Z1[0],$Z2[0],0,0,0,0);

```

```
            print "Zok_unique\t".($TnInsertion_XY)."\t".( $Locus_BCG_Z)."\tXY $XYid1".($XYid2)."\t"."Z $Zid1".($Zid2)."\n";
       push(@{$stack_Zok{$XYid1.$XYid2}{$Zid1.$Zid2}},"$TnInsertion_XY\t$Locus_BCG_Z");
       $totalidentifications++;
       $numZok++;
         }
      else{
        $numpos=scalar(@XY2s);
        if($numpos < 5){
            print "Zok_nonunique\t".($TnInsertion_XY)."\t".( $Locus_BCG_Z);
       for(my $i=0;$i<$numpos;$i++){
         (my $XYid1, my $XYid2, my $Zid1, my $Zid2)=get_plate_id(\@valsXY,\@valsZ,$XY1s[0],$XY2s[$i],$Z1[0],$Z2[0],0,$i,0,0);
          print "\tXY $XYid1".($XYid2)."\t"."Z $Zid1".($Zid2);
       }
       print "\n";
       $totalidentifications++;
       $numZok++;
           }
      else{
         $impossible++;
       }
      }
       }
    else{
      if($fraction > $lower_ratio){
        $numpos=1;
        (my $XYid1, my $XYid2, my $Zid1, my $Zid2)=get_plate_id(\@valsXY,\@valsZ,$XY1s[0],$XY2s[0],$Z1[0],$Z2[0],0,0,0,0);

           print "Zok_unique\t".($TnInsertion_XY)."\t".( $Locus_BCG_Z)."\tXY $XYid1".($XYid2)."\t"."Z $Zid1".($Zid2)."\n";
        push(@{$stack_Zok{$XYid1.$XYid2}{$Zid1.$Zid2}},"$TnInsertion_XY\t$Locus_BCG_Z");
        $totalidentifications++;
        $numZok++;

         }
      else{
      $numpos=scalar(@XY2s);
           if($numpos < 5){
            print "Zok_nonunique\t".($TnInsertion_XY)."\t".( $Locus_BCG_Z);
       for(my $i=0;$i<$numpos;$i++){
         (my $XYid1, my $XYid2, my $Zid1, my $Zid2)=get_plate_id(\@valsXY,\@valsZ,$XY1s[0],$XY2s[$i],$Z1[0],$Z2[0],0,$i,0,0);
          print "\tXY $XYid1".($XYid2)."\t"."Z $Zid1".($Zid2);
       }
       print "\n";
       $totalidentifications++;
       $numZok++;
           }
      else{
         $impossible++;
       }
       }
    }
    }
    ###Case 4b###: several values in @XY1, one value in @XY2
    #Check if first element of XY1s is comparable, and the next element is significantly less
    #Check if the first element is > split_threshold, if yes check if the fraction of the first and second element is > upper_ratio (least stringent), if yes we have a Zok_unique identification, if no we have a Zok_nonunique identification.
```

```
   #If the first element is not bigger than the split threshold, it is checked if the fraction of the first and second element > lower_ratio (more stringent), if yes we have a Zok_unique identification, if no we have a Zok_nonunique identification.
   elsif((scalar(@XY2s)==1) && (scalar(@XY1s)>1)){
      my $fraction=$XY1s[0]/$XY1s[1];
   if($XY1s[0]>=$split_threshold){
     if($fraction >  $upper_ratio){
      $numpos=1;
      (my $XYid1, my $XYid2, my $Zid1, my $Zid2)=get_plate_id(\@valsXY,\@valsZ,$XY1s[0],$XY2s[0],$Z1[0],$Z2[0],0,0,0,0);

           print "Zok_unique\t".($TnInsertion_XY)."\t".( $Locus_BCG_Z)."\tXY $XYid1".($XYid2)."\t"."Z $Zid1".($Zid2)."\n";
      push(@{$stack_Zok{$XYid1.$XYid2}{$Zid1.$Zid2}},"$TnInsertion_XY\t$Locus_BCG_Z");
      $totalidentifications++;
      $numZok++;

        }
     else{
      $numpos=scalar(@XY1s);
           if($numpos < 5){
            print "Zok_nonunique\t".($TnInsertion_XY)."\t".( $Locus_BCG_Z);
      for(my $i=0;$i<$numpos;$i++){
        (my $XYid1, my $XYid2, my $Zid1, my $Zid2)=get_plate_id(\@valsXY,\@valsZ,$XY1s[$i],$XY2s[0],$Z1[0],$Z2[0],$i,0,0,0);
         print "\tXY $XYid1".($XYid2)."\t"."Z $Zid1".($Zid2);
      }
      print "\n";
      $totalidentifications++;
      $numZok++;
          }
      else{
         $impossible++;
      }
     }
      }
   else{
     if($fraction > $lower_ratio){
       $numpos=1;
        (my $XYid1, my $XYid2, my $Zid1, my $Zid2)=get_plate_id(\@valsXY,\@valsZ,$XY1s[0],$XY2s[0],$Z1[0],$Z2[0],0,0,0,0);

           print "Zok_unique\t".($TnInsertion_XY)."\t".( $Locus_BCG_Z)."\tXY $XYid1".($XYid2)."\t"."Z $Zid1".($Zid2)."\n";
      push(@{$stack_Zok{$XYid1.$XYid2}{$Zid1.$Zid2}},"$TnInsertion_XY\t$Locus_BCG_Z");
      $totalidentifications++;
      $numZok++;

        }
      else{
      $numpos=scalar(@XY1s);
           if($numpos < 5){
            print "Zok_nonunique\t".($TnInsertion_XY)."\t".( $Locus_BCG_Z);
      for(my $i=0;$i<$numpos;$i++){
        (my $XYid1, my $XYid2, my $Zid1, my $Zid2)=get_plate_id(\@valsXY,\@valsZ,$XY1s[$i],$XY2s[0],$Z1[0],$Z2[0],$i,0,0,0);
         print "\tXY $XYid1".($XYid2)."\t"."Z $Zid1".($Zid2);
      }
      print "\n";
      $totalidentifications++;
      $numZok++;
          }
      else{
```

```
          $impossible++;
       }
       }
     }
    }
    ###Case 4c###: several values in @XY1 AND in @XY2
     #Check for both if the first (biggest) value is >split_threshold AND >upper_ratio OR < split_threshold AND >lower_ratio. If true we have Zok_unique identification, if false we have an impossible case.
     elsif((scalar(@XY1s)>1) && (scalar(@XY2s)>1)){
        my $fraction1=$XY1s[0]/$XY1s[1];
        my $fraction2=$XY2s[0]/$XY2s[1];
        my $pass1=0;
     my $pass2=0;
     if($XY1s[0]>=$split_threshold){
       if($fraction1 > $upper_ratio){
        $pass1=1;
          }
        }
     else{
       if($fraction1 > $lower_ratio){
          $pass1=1;
           }
      }
     if($XY2s[0]>=$split_threshold){
       if($fraction2 > $upper_ratio){
        $pass2=1;
          }
        }
     else{
       if($fraction2 > $lower_ratio){
          $pass2=1;
           }
      }
        if($pass1 && $pass2){
       $numpos=1;
          (my $XYid1, my $XYid2, my $Zid1, my $Zid2)=get_plate_id(\@valsXY,\@valsZ,$XY1s[0],$XY2s[0],$Z1[0],$Z2[0],0,0,0,0);

          print "Zok_unique\t".($TnInsertion_XY)."\t".( $Locus_BCG_Z)."\tXY $XYid1".($XYid2)."\t"."Z $Zid1".($Zid2)."\n";
       push(@{$stack_Zok{$XYid1.$XYid2}{$Zid1.$Zid2}},"$TnInsertion_XY\t$Locus_BCG_Z");
       $totalidentifications++;
       $numZok++;

      }
     else{
        $numpos=scalar(@XY1s)*scalar(@XY2s);
        $impossible++;
      }
     }
    else{
    die "Other case ".(scalar(@XY1s))." ".(scalar(@XY2s))." "."\n";
    }
     if(defined($Zokhash{$numpos})){
     $Zokhash{$numpos}=$Zokhash{$numpos}+1;
      }
     else{
     $Zokhash{$numpos}=1;
      }
   }
   ##Case 5##: no unique identification, not for XY (several values for XY) and not for Z (several values for Z). Both $XYok>1 and $Zok>1. We need some heuristics.
   else{
```

```
    #For XY1 and XY2. Sort the values in @XY1 and @XY2, from big to small. Then check the first element of XY1 and XY2 to check if they meet the thresholds (like for case c)
    my @XY1s=sort{$b<=>$a} @XY1;
    my @XY2s=sort{$b<=>$a} @XY2;

    if((scalar(@XY1s)==1) && (scalar(@XY2s)>1)){
        my $fraction=$XY2s[0]/$XY2s[1];
    if($XY2s[0]>=$split_threshold){
      if($fraction >   $upper_ratio){
       $XYok=1;
          }
        }
    else{
      if($fraction > $lower_ratio){
         $XYok=1;
           }
    }
    }
    elsif((scalar(@XY2s)==1) && (scalar(@XY1s)>1)){
        my $fraction=$XY1s[0]/$XY1s[1];
    if($XY1s[0]>=$split_threshold){
      if($fraction >   $upper_ratio){
       $XYok=1;
          }
        }
    else{
      if($fraction > $lower_ratio){
         $XYok=1;
           }
    }
   }
    elsif((scalar(@XY1s)>1) && (scalar(@XY2s)>1)){
        my $fraction1=$XY1s[0]/$XY1s[1];
        my $fraction2=$XY2s[0]/$XY2s[1];
        my $pass1=0;
    my $pass2=0;
    if($XY1s[0]>=$split_threshold){
      if($fraction1 > $upper_ratio){
       $pass1=1;
          }
        }
    else{
      if($fraction1 > $lower_ratio){
         $pass1=1;
           }
    }
    if($XY2s[0]>=$split_threshold){
      if($fraction2 > $upper_ratio){
       $pass2=1;
          }
        }
    else{
      if($fraction2 > $lower_ratio){
         $pass2=1;
           }
    }
        if($pass1 && $pass2){
       $XYok=1;
    }
   }

    #For Z1 and Z2. Sort the values in @Z1 and @Z2, from big to small. Then check the first element of Z1 and Z2 to check if they meet the thresholds (like for case c)
```

```
 my @Z1s=sort{$b<=>$a} @Z1;
 my @Z2s=sort{$b<=>$a} @Z2;

 if((scalar(@Z1s)==1) && (scalar(@Z2s)>1)){
     my $fraction=$Z2s[0]/$Z2s[1];
 if($Z2s[0]>=$split_threshold){
   if($fraction >   $upper_ratio){
    $Zok=1;
      }
     }
 else{
   if($fraction > $lower_ratio){
      $Zok=1;
       }
 }
 }
 elsif((scalar(@Z2s)==1) && (scalar(@Z1s)>1)){
     my $fraction=$Z1s[0]/$Z1s[1];
 if($Z1s[0]>=$split_threshold){
   if($fraction >   $upper_ratio){
    $Zok=1;
      }
     }
 else{
   if($fraction > $lower_ratio){
      $Zok=1;
       }
 }
}
 elsif((scalar(@Z1s)>1) && (scalar(@Z2s)>1)){
     my $fraction1=$Z1s[0]/$Z1s[1];
     my $fraction2=$Z2s[0]/$Z2s[1];
     my $pass1=0;
 my $pass2=0;
 if($Z1s[0]>=$split_threshold){
   if($fraction1 > $upper_ratio){
    $pass1=1;
      }
     }
 else{
   if($fraction1 > $lower_ratio){
      $pass1=1;
       }
 }
 if($Z2s[0]>=$split_threshold){
   if($fraction2 > $upper_ratio){
    $pass2=1;
      }
     }
 else{
   if($fraction2 > $lower_ratio){
      $pass2=1;
       }
 }
     if($pass1 && $pass2){
    $Zok=1;
 }
}
#If all thresholds are met for XY1, XY2, Z1 and Z2 we have determined the position heuristically.
 if($XYok and $Zok){
   $numcase1++;
   $totalidentifications++;
```

```
      (my $XYid1, my $XYid2, my $Zid1, my $Zid2)=get_plate_id(\@valsXY,\@valsZ,$XY1s[0],$XY2s[0],$Z1s[0],$Z2s[0],0,0,0,0);

      print "Heuristic\t".($TnInsertion_XY)."\t".( $Locus_BCG_Z)."\tXY $XYid1".($XYid2)."\t"."Z $Zid1".($Zid2)."\n";
      push(@{$stack_heuristic{$XYid1.$XYid2}{$Zid1.$Zid2}},"$TnInsertion_XY\t$Locus_BCG_Z");

    }
    else{
      $impossible++;
    }

  }

}
close XY;
close Z;
#The calculations are performed

open(SUMFILE,">Summary.txt");
print SUMFILE "$numok good lines\n"; #total number of lines (tn insertions)
print SUMFILE "$impossible impossible cases\n";
print SUMFILE "$totalidentifications total identifications\n";
print SUMFILE "with the following details:\n";
print SUMFILE "\t$numunique unique identifications\n";
print SUMFILE "\t$numXYok identifications 'XY ok'\n";
print SUMFILE "\t$numZok identifications 'Z ok'\n";
print SUMFILE "\t$numcase1 heuristic identifications\n";

close SUMFILE;

#foreach my $key(sort {$a<=>$b} keys %XYokhash){
#  my $val=$XYokhash{$key};
#  print "$key -> $val\n";
#}

#Print all unique identifications (1 well 1 mutant) to file. In this file the TnInsertions are ordered by plate. The file indicates if the wells are filled with one unique mutant or if the wells contain more than one mutant (according to the protocol).
open(OUT,">TnListOrderedByPlate.txt");
foreach my $key1 (keys %stack_unique){
 foreach my $key2 (keys %{$stack_unique{$key1}}){
     my $numvals=scalar(@{$stack_unique{$key1}{$key2}});
     if($numvals==1){
   my $TnI=${$stack_unique{$key1}{$key2}}[0];
   print OUT "$TnI\tXY: $key1\tZ: $key2\tUNIQUE\n";
     }
     else{
    for(my $i=0;$i<$numvals;$i++){
      my $TnI=${$stack_unique{$key1}{$key2}}[$i];
      print OUT "$TnI\tXY: $key1\tZ: $key2\tMZS with $numvals\n";
       }
     }
 }
}
close OUT;

close FH;

# output files from the above code
# - "TnListOrderedByPlate.txt" In this file the TnInsertions are ordered by plate. The file indicates if the wells are filled with one unique mutant or if the wells contain more than one mutant (according to the protocol). The file contains 5 columns: (c1) TnInsertion-number, (c2) locus, (c3) XY coordinate, (c4) Z coordinate, (c5) information about uniqueness of the well
```

```
# - "Summary.txt". The file contains the summary statistics.
# - "TnListOrderedByMutant.text" In this file the TnInsertions are ordered by mutant. The list indicates if the automatic deduction of position of the mutant was unique, Zok_unique, Zok_nonunique, XYok_unique, XYok_nonunique or heuristically determined. The file contains 5-n columns: (c1)information about how the position was deducted, (c2) TnInsertion-number, (c3) locus, (c4) XY coordinate, (c5) Z coordinate, (c6-cn) additional XY or Z coordinates

# end CP-CSeq protocol
##################################################################

##################################################################
# Addition by Katlyn Borgers to create a clean and reordered output (compiled by Laurent Schindfessel)

# read output file
my $filename = 'TnListOrderedByMutant.txt';
open(my $fh, '<:encoding(UTF-8)', $filename)
 or die "Could not open file '$filename' $!";

# open file to write new output
open(OUTGOOD, '>', 'FinalTnList.txt') or die "cannot open file";

# loop through every row
while (my $row = <$fh>) {
 # remove last character, which is newline "\n"
 my $shorterRow = substr($row, 0, -1);

 # split string in components, using fixed structure
 my @splitted = split(/\s+/, $shorterRow );

 # reorder the first columns
 my $stringTemp = @splitted[1]."\t".@splitted[2]."\t".@splitted[0];

 # loop over each Z- and XY-column of txt file, starting at 4, going to infinity

  for(my $iter = 6; $iter < @splitted; $iter = $iter + 4 ) {
     # extract character that determines row of plate
    my $rowChar = substr(@splitted[$iter], -1);
    # extract number that determines column of plate
    my $colNumb = substr(@splitted[$iter ],0,-1);
    # calculate plate number
    my $plateNumber = (ord($rowChar)-ord("A"))*12 + $colNumb ;

    # reverse order of XY
    my $goodXY = substr(@splitted[$iter-2], -1).substr(@splitted[$iter-2],0,-1);

    # put output together
    $stringTemp = $stringTemp."\tSet-".$librarySet.".Plate-".($plateNumber).".Well-".$goodXY ;
  }

 print OUTGOOD $stringTemp."\n";
 #print OUTGOOD "$shorterRow"."\t".($plateNumber )."\n";
}
close OUTGOOD;
close $fh;

# output file from the above code
# - "FinalTnList.txt". This output file contains coordinates for each mutant (= row) in the list, provided that sufficient data for that mutant is present in the dataset This file is a cleaned and reordered version of the file "TnListOrderedByMutant.text". The file contains 4-n columns: (c1)TnInsertion-number,(c2)locus,(c3)information about how the position was deducted,(c4)position, (c5-cn) alternative positions for the mutant. The position information is built as followed: Set-x1.Plate-x2.Well-x3 (e.g. Set-I.Plate-23.Well-A3), with x1 the Roman number for the library set, x2 the plate number, x3 the well coordinates.
```

```
# end additions by Katlyn Borgers
##################################################################

##################################################################
# Subroutines for the CP-CSeq protocol

sub is_integer {
  defined $_[0] && $_[0] =~ /^[+-]?\d+$/;
}

# New arguments 'exaequo' were introduced for the cases where there are equal values in the coverages. Due to the way this program is implemented, the function get_plate_id does not know how the coverage arrays are sorted in the program. So when there is equal coverage for a non-unique case two times the same solution is given, instead of the two different ones. Optimally, the sorting in this program should be restructured, but by creating the new 'exaequo' arguments the problem is solved. The arguments 'exaequo' allow to select a value associated with a, multiple times occurring, coverage.
# Explanation of the code: Suppose a certain coverage needs to be found, this function will return the first equal value in case exaequo = 0. If exaequo = 1, it will return the second equal value, exaequo = 2 returns the third equal value, etc. This corresponds with Perl counting from zero. If there is no equal coverage corresponding to the exaequo value, this function will return the last equal value.
# Note: first, second and third correspond to the ordering in the arrays $vXY and $vZ. By implementing exaequo for all parameters, this function can be used to return all equal values if used in a for loop.
sub get_plate_id{
  (my $vXY,my $vZ, my $XY1,my $XY2,my $Z1,my $Z2, my $exaequo_XY1, my $exaequo_XY2, , my $exaequo_Z1, , my $exaequo_Z2 )=@_;
  my @valsXY=@{$vXY};
  my @valsZ=@{$vZ};

  my @letters=("A","B","C","D","E","F","G","H");

  my $XYid1;
  my $XYid2;
  my $Zid1;
  my $Zid2;

  #Write out this identification
  my $found_counter = 0;
  for(my $i=0;$i<12;$i++){
  if($valsXY[8+$i] eq $XY1){
      $XYid1=1+$i;
      if($found_counter eq $exaequo_XY1){
        last;
      }
      $found_counter++;
    }
  }

  $found_counter = 0;
  for(my $i=0;$i<8;$i++){
    if($valsXY[21+$i] eq $XY2){
      $XYid2=$letters[$i];
      if($found_counter eq $exaequo_XY2){
        last;
      }
      $found_counter++;
    }
  }

  $found_counter = 0;
  for(my $i=0;$i<12;$i++){
    if($valsZ[8+$i] eq $Z1){
     $Zid1=1+$i;
     if($found_counter eq $exaequo_Z1){
```

```
            last;
        }
        $found_counter++;
      }
    }

    $found_counter = 0;
    for(my $i=0;$i<8;$i++){
      if($valsZ[21+$i] eq $Z2){
        $Zid2=$letters[$i];
        if($found_counter eq $exaequo_Z2){
            last;
        }
        $found_counter++;
      }
    }
   return(($XYid1,$XYid2,$Zid1,$Zid2));
}

# end subroutines
###################################################################
```

### References with supplementary information

7. Gupta R, Barkan D, Redelman-Sidi G, Shuman S, Glickman MS. 2011. Mycobacteria exploit three genetically distinct DNA double-strand break repair pathways. Mol Microbiol 79.

13. Hosomichi K, Mitsunaga S, Nagasaki H, Inoue I. 2014. A Bead-based Normalization for Uniform Sequencing depth (BeNUS) protocol for multi-samples sequencing exemplified by HLA-B. BMC Genomics 15.
